# Cross-species analysis of GNB1 I80T encephalopathy: conserved developmental, epileptic and neuronal transcriptome signatures

**DOI:** 10.64898/2026.08.03.742477

**Authors:** Haritha P. Reddy, Vigneshwar Ranjan, Merav Klo, Guy Shapira, Haim Bassan, Gilad Harel, Gali Heimer, Bruria Ben Zeev, Tatiana Rabinski, Gad D. Vatine, Yakey Yaffe, Ben M. Maoz, Lior Bikovski, Noam Shomron, Daniel Yakubovich, Moran Rubinstein, Nathan Dascal

## Abstract

GNB1 encephalopathy (GNB1E) is a rare neurodevelopmental disorder caused by mutations in GNB1 gene encoding the G protein subunit Gβ1. Mechanisms linking these variants to neurological dysfunction remain unclear. We investigated the prevalent p.Ile80Thr (I80T) variant using combined clinical, cellular, and *in vivo* approaches. Longitudinal evaluation of a GNB1E patient revealed developmental delay, progressive peripheral spasticity, and epilepsy with Spike-Wave Activation in Sleep. Heterozygous knock-in *Gnb1*^I80T/+^ mice exhibited disease-relevant phenotypes, including impaired early development, mild adult motor and cognitive deficits and epileptiform cortical spike-and-wave discharges. Transcriptomic analysis identified 323 genes concordantly dysregulated in mouse cortex and cortical human neuronal cultures from patient-derived induced pluripotent cells. This gene set was enriched for ion-channel function, epilepsy-associated genes, and G_s_/adenylyl cyclase signaling pathway. Our integrated analysis establishes the first cross-species model for GNB1E, suggests common neurological mechanisms and molecular pathways linked to GNB1E, and provides a framework for mechanistic and therapeutic studies.

**Teaser:** Conserved human/mouse neurological and transcriptomic signatures in *GNB1* encephalopathy.

## Introduction

*GNB1* encephalopathy (GNB1E) is a rare neurodevelopmental disorder caused by more than 40 pathogenic variants in *GNB1*, which encodes the β1 subunit of heterotrimeric G proteins (Gβ1). *GNB1* mutations were initially identified as oncogenic drivers in hematological malignancies, with mutations at residues K57, I80, and K89 shown to activate proliferative signaling pathways and confer resistance to targeted therapies (*1*). Subsequently, predominantly *de novo* germline *GNB1* variants were identified as the cause of GNB1E, a severe neurodevelopmental disorder characterized by global developmental delay, intellectual disability, hypotonia, dystonia, epilepsy, and movement disorders (*2–5*). Disease-associated variants frequently affect functionally important residues within the WD40 β-propeller domain of Gβ1 (*4, 5*). Despite increasing recognition of GNB1E and expansion of its clinical spectrum, the molecular and neuronal mechanisms linking specific *GNB1* variants to disease remain incompletely understood.

Gβ1 is a core component of heterotrimeric G proteins, which link G-protein-coupled receptors (GPCRs) to diverse intracellular signaling pathways (*6–8*). In the inactive state, GDP-bound Gα associates with the Gβγ heterodimer to form the heterotrimeric G-protein complex. Upon GPCR activation, the receptor promotes GDP-GTP exchange on Gα, leading to conformational changes and reorganization of the G-protein complex that enable Gα and Gβγ to regulate downstream effectors. Gβγ is ubiquitous and regulates plethora of cellular processes including ion channels, synaptic proteins and signaling enzymes, profoundly regulating neuronal excitability and synaptic function (*9–13*). Disease-associated GNB1 variants are predicted to disrupt critical protein interactions within the G-protein signaling complex and alter downstream signaling (*14–18*). Functional studies have demonstrated that GNB1E-associated variants can alter GPCR coupling and Gβγ-dependent regulation of G-protein-activated inwardly rectifying potassium (GIRK) channels, implicating dysregulated ion-channel signaling in neuronal dysfunction (*3, 17–19*). Our previous studies extended these mechanistic observations to a *Gnb1*^K78R/+^ mouse model, which recapitulated developmental and epileptic phenotypes associated with GNB1E. Pharmacological modulation of GIRK channel activity suppressed spike-and-wave discharges (SWDs) in vivo and restored network activity in vitro, highlighting Gβγ-dependent ion-channel signaling as a potential therapeutic target (*19, 20*).

Despite incremental progress, the mechanisms by which variant-associated perturbations in G protein signaling give rise to broader molecular and functional changes in the nervous system remain poorly understood. Elucidating these mechanisms is particularly challenging owing to the diversity of Gβγ-regulated cellular processes, and different pathogenic variants are expected to differentially affect specific signaling pathways. This concept is supported by the marked phenotypic heterogeneity observed among affected individuals (*5*) and studies in heterologous cellular models (*3, 17–19*). However, efforts to understand genotype–phenotype relationships remain limited by the scarcity of systematically collected clinical and animal model data. At present, only a preliminary patient registry maintained by a patient advocacy organization, with about 100 entries, is available (*21*), and no comprehensive natural history database exists. Furthermore, previous studies have largely examined clinical cohorts or individual experimental models in isolation, and the molecular consequences of GNB1 mutations across human and in vivo disease models remain poorly understood.

Here, we used an integrated multimodal cross-species approach to characterize the phenotypic and molecular consequences of the most prevalent GNB1E-associated variant, p.Ile80Thr (I80T), in a human patient and a new *Gnb1*^I80T/+^ mouse model. The I80T variant was reported in more than 20% of affected individuals (*5, 22*). Clinically, it is representative of the disorder, being consistently associated across independent patient cohorts with its core manifestations, including developmental delay, epilepsy, and motor and movement abnormalities (*2, 4, 23–26*).

Our study reveals developmental, behavioral, and electroencephalographic abnormalities in *Gnb1*^I80T/+^ mice that are consistent with the clinical manifestations observed during the longitudinal evaluation of a patient carrying the GNB1 I80T variant. Remarkably, the concordance between patient and mouse phenotypes was paralleled by convergent transcriptomic alterations in the mouse cortex and in neuronal cultures derived from patient-derived induced pluripotent stem cells (iPSCs). Together, these findings bridge clinical observations and the new experimental disease model through shared molecular signatures, providing insights into the conserved pathways that underlie GNB1 encephalopathy.

## Results

To approach the mechanisms of GNB1 I80T Encephalopathy, we integrated a longitudinal patient case study with developmental, behavioral and electrophysiological investigation of the newly generated *Gnb1*^I80T/+^ mouse model, along with comparative transcriptomic (RNAseq) analyses of *Gnb1*^I80T/+^ mouse brain cortex and neuronal cultures produced *in vitro* from patient-derived human iPSCs (hiPSCs) (Fig. 1A).

**Figure 1:**
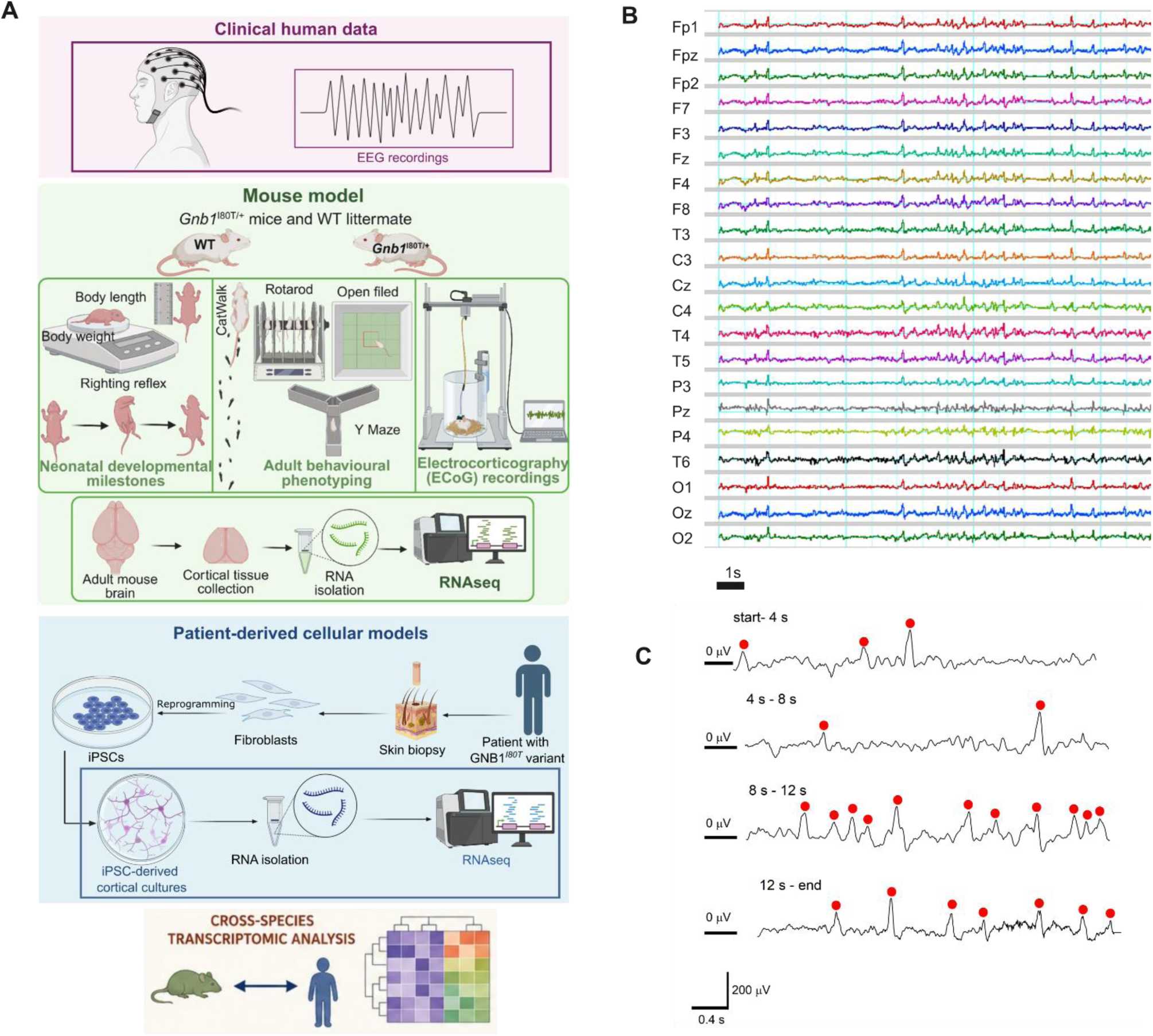
The general scheme of the study and a representative EEG recording of case patient. (**A**) Graphical scheme of the study parts and stages. (**B, C**) A representative EEG recording from the case patient at the age of 11 years and 6 months. EEG was recorded utilizing 10-20 electrode placement system. Sampling frequency was 256 Hz. **B** shows all recorded channels during an eye-closed interval. The record length is 17 s. **C** is a zoom on channel T3 on higher time and amplitude resolution scales. Spikes corresponding to epileptic activity are marked with red circles.

### Patient clinical description

The patient is a 17.5-year-old female with a pathogenic *de novo* GNB1 variant (chr1.1737942A>G; c.239T>C; p.Ile80Thr, I80T), identified and confirmed by trio whole-exome sequencing. A brief initial report about the patient was described by Hemati et al. 2018 (*4*) (patient 5) at the age of 8.5 years. The patient was born at 37 weeks’ gestation by cesarean section with a birth weight of 2700 g and normal Apgar scores. Family history was unremarkable. The neonatal period manifested with marked hypotonia, excessive sleepiness and weak cry. Hypotonia persisted during infancy, accompanied by head nodding, nystagmus, and global developmental delay (*4*). Toward the end of the first year, peripheral tone gradually increased, head nodding resolved, and nystagmus markedly improved.

During early childhood, the patient developed marked limb spasticity pyramidal signs, dystonia and bradykinesia. Development was characterized by severe motor delay, expressive language impairment, feeding difficulties, and relatively preserved nonverbal cognitive abilities. She received early intervention and entered special education at 18 months, when her developmental level corresponded to approximately 9 months. No clinical seizures occurred during infancy, and awake EEG at 12 months was normal.

However, after age of 6 years, the EEG demonstrated frequent independent bilateral centrotemporal epileptiform discharges, increasing during sleep to 50–80% of the recording (Fig. 1B, C; Supplementary Figure 1) and fulfilling the clinical criteria for spike-wave activation in sleep (SWAS). Parents reported brief staring episodes (not captured on EEG). Sulthiame did not improve the EEG but was associated with subjective improvement in language. At 8.5 years, she experienced status epilepticus with generalized clonic seizures and right eye deviation during febrile illness, prompting initiation of valproic acid. No further generalized seizures occurred, although intermittent staring episodes, nystagmus, and truncal myoclonic jerks, particularly during febrile illnesses, persisted and may have represented brief seizures. Treatment with levetiracetam, clobazam, ethosuximide, lamotrigine, and cannabis oil (CBD/THC 20:1) were sequentially introduced and failed to substantially reduce epileptiform EEG activity. From age 12 years, she remained on sulthiame and cannabis oil, and by 16 years EEG abnormalities improved, probably spontaneously since the pharmacotherapy remained unchanged (Supplementary Figure 1). For further details on pharmacotherapy, see Supplementary Fig. 1 legend.

Motor course was characterized by progressive spasticity, dystonia, bradykinesia and oromotor dyspraxia, right hip subluxation, and bilateral patella alta. She received four courses of botulinum toxin injections. Oral baclofen and diazepam were ineffective, and trihexyphenidyl was declined. At 8.5 years, she could sit independently, stand with support, and walk using a walker. Despite intensive rehabilitation and CBD oil treatment, she gradually regressed and lost ambulation.

Brain MRI at 6 and 21 months showed relative thickening of the corpus callosum and hippocampi, prominence of the frontal and temporal horns, and later at 6.5 years, additional mild posterior periventricular white matter hyperintensities. Spine MRI, echocardiography, ECG, abdominal ultrasound, extensive metabolic investigations including CSF neurotransmitters, kidney, thyroid, and liver function tests were unremarkable.

At 17.5 years, she is wheelchair-dependent, requires complete assistance with daily living activities, attends a special education school, has dysarthria with intermittent drooling, reads with assistance, and communicates using augmentative and alternative communication.

### Characterization of the Gnb1^I80T/+^ mouse

To model the GNB1 encephalopathy-associated I80T variant, we generated a *Gnb1*^I80T^ knock-in mouse line using CRISPR/Cas9. The mutation was confirmed by PCR and Sanger sequencing (Supplementary Figure 2). Homozygous *Gnb1*^I80T/I80T^ mice were not viable, and heterozygous *Gnb1*^I80T/+^ mice showed reduced viability on the C57BL/6NJ background. Therefore, subsequent experiments were performed using mice repeatedly backcrossed toward the FVB background for 8-10 generations (Supplementary Figure 2A). *Gnb1^I80T/+^* mice and wild-type (WT) littermate controls were used to characterize disease-associated phenotypes *in vivo*. Mice underwent assessment of neonatal developmental milestones, adult behavioral phenotyping, and electrocorticography (ECoG) recordings. In addition, cortical tissue was collected for transcriptomic profiling by RNA sequencing (RNAseq).

#### Gnb1^I80T/+^ mice show early developmental delay

To assess early physical and sensorimotor development, *Gnb1*^I80T/+^ pups and WT littermates were evaluated from postnatal day (P)4 to P18. *Gnb1*^I80T/+^ pups were visibly smaller than WT littermates and showed significantly reduced body weight throughout early postnatal development, accompanied by reduced body length (Fig. 2A-C). Sex-dependent differences in body weight are shown in Supplementary Figure 3A,F. Body length was similar in males and females during the neonatal period (Supplementary Figure 3B). Female *Gnb1*^I80T/+^ pups showed delayed eye opening compared with WT females (Supplementary Figure 3C).

**Figure 2.**
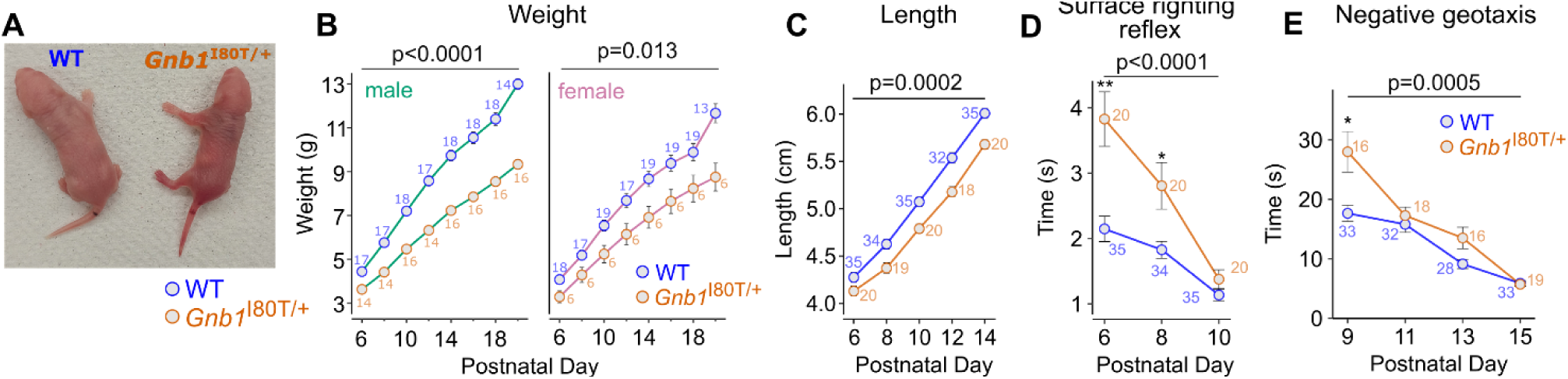
*Gnb1*^I80T/+^ mice exhibit developmental delay throughout the neonatal period. (**A**) Representative image of WT (left) and *Gnb1*^I80T/+^ (right) pups at postnatal day 6 (P6) belonging to the same litter. (**B**–**E**) Assessment of developmental milestones from P6 to P20. (**B**) Body weight (sex-stratified; see Supplementary Figure 3A for sex effects). (**C**) Body length. (**D**, **E**) Motor reflexes: surface righting latency and negative geotaxis (latency to reorient 180° upward from a downward-facing starting position). Error bars represent mean ± SEM; sample sizes are indicated beside each data point. Statistics: (**B**–**E**) mixed-effects analysis with Šídák’s multiple comparisons test. *\*p* < 0.05; *\*\*p* < 0.01.

Assessment of developmental milestones revealed a transient delay in sensorimotor maturation. *Gnb1*^I80T/+^ pups showed impaired surface righting reflex performance during the early postnatal period, particularly up to P8, but this deficit was not sustained at later times (Fig. 2D). Mutant pups also exhibited delayed negative geotaxis performance at P9, with no persistent deficit thereafter (Fig. 2E). There were no differences in both surface righting and negative geotaxis between males and females, both WT and *Gnb1*^I80T/+^ (Supplementary Figure 3E,D).

Together, these findings indicate that male and female *Gnb1*^I80T/+^ mice exhibit early postnatal growth impairment and transient delays in developmental sensorimotor milestones, recapitulating aspects of neurodevelopmental delay observed in GNB1 encephalopathy.

#### Gnb1^I80T/+^ mice exhibit frequent spike-and-wave discharges and abnormal cortical oscillatory activity

Some human GNB1 patients were reported to undergo recurrent spontaneous convulsive seizures, whereas in many others such seizures were not observed (*2, 4, 5*) but, as in our patient, significant epileptiform activity was revealed in EEG (Fig. 1, Supplementary Figure 1). We did not observe clear convulsive seizures in the *Gnb1*^I80T/+^ mouse colony. However, in ECoG recordings in freely behaving adult mice aged 4 to 8 weeks, we observed unambiguous abnormal cortical excitability in *Gnb1*^I80T/+^ mice in the form of bilateral spike-and-wave discharges (SWDs), that were completely absent in WT littermates (Fig. 3, Supplementary Figure 4). These discharges showed the characteristic abrupt onset and termination of repetitive spike-and-wave complexes and closely resembled the absence-like SWDs previously described in the *Gnb1*^K78R/+^ mouse model (*19, 31*).

**Figure 3:**
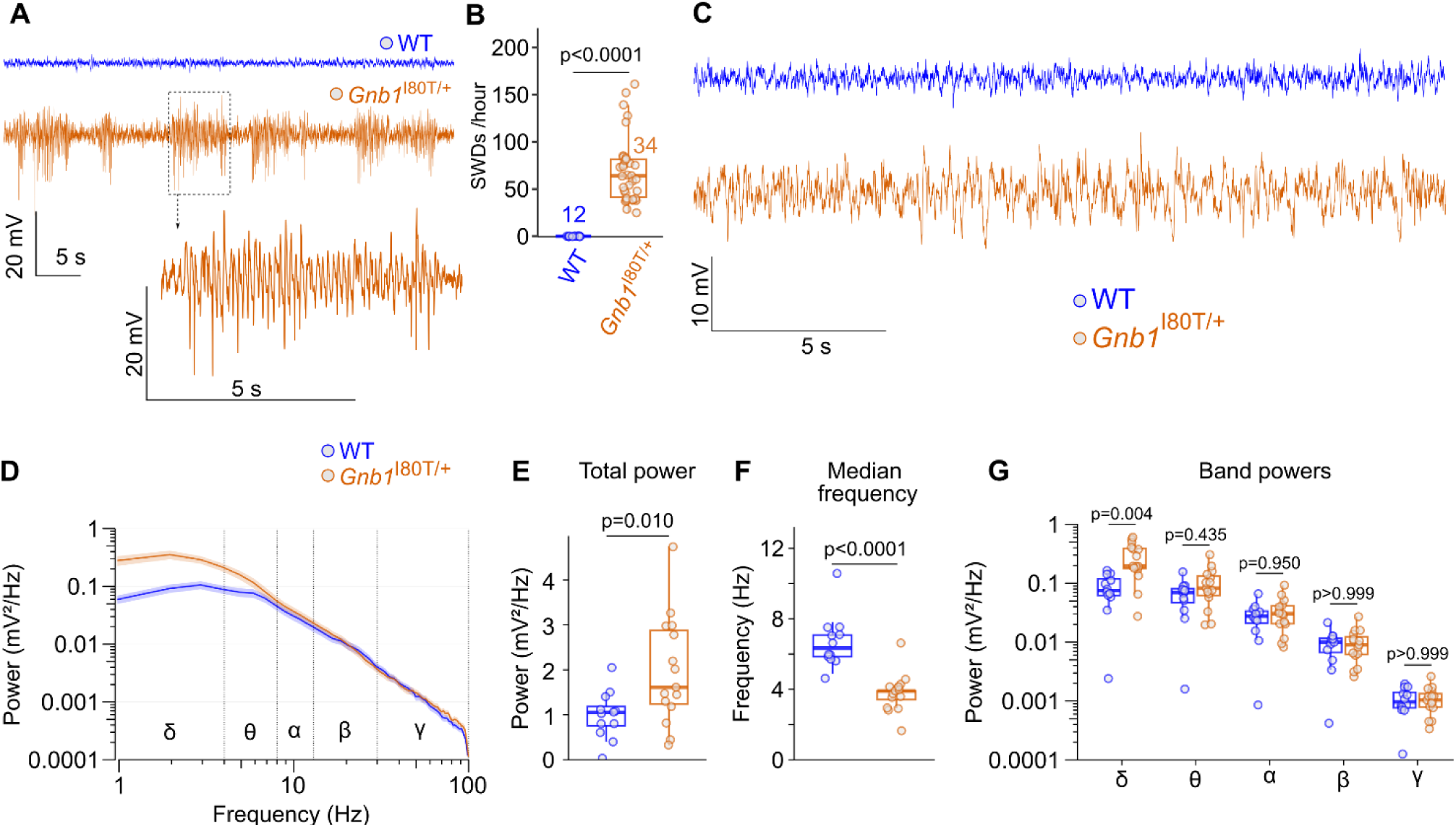
Spike-wave discharges and increased power in the δ band in the background in *Gnb1*^I80T/+^ mice. (**A**) Representative ECoG traces of WT and *Gnb1*^I80T/+^ mice obtained from right somatosensory cortex. WT mice show no SWDs. (**B**) Total number of SWDs recorded in WT and *Gnb1*^I80T/+^ mice during one hour following a 30-minutes habituation period (see Supplementary Figure 4 for further details). Boxes represent median and interquartile range; whiskers extend to 1.5 × IQR; individual data points represent single animals. WT (blue circles): *n* = 6 males, 6 females, P28–54; *Gnb1*^I80T/+^: *n* = 20 males, 14 females, P34–56. Statistics: Mann-Whitney test. (**C-G**) Analysis of ECoG background activity of 4–8 week old mice (P28–P55) from record segments where no SWDs were detected. (**C**) Example segments of ECoG without SWDs in wild-type (WT) and *Gnb1*^I80T/+^ mice. (**D**) Power spectral density (PSD). (**E**) Total power. Statistics: Welch’s t-test. (**F**) median frequency. Mann-Whitney test. (**G**) Total power within frequency bands: δ-0.9–3.9 Hz; θ-4.8–7.8 Hz; α-8.7–12.7 Hz; β-13.6–29.3 Hz; γ-30.2–99.6 Hz (two-way ANOVA with Šídák’s multiple comparisons test). Throughout **E-G**, boxes represent median and interquartile range; whiskers extend to 1.5 × IQR; individual data points represent single animals. WT: *n* = 6 males, 6 females; *Gnb1*^I80T/+^: *n* = 6 males, 9 females. Only highest quality records were selected for PSD analysis (D-G), hence the lower the total number of animals analyzed than for SWD (B).

Analysis of ECoG recordings revealed very similar total SWD count, duration, and SWD spike content in female and male *Gnb1*^I80T/+^ mice (Supplementary Figure 4B). Therefore, for subsequent analysis we pooled female and male ECoG data. SWDs occurred at a frequency of ∼70 SWD/hour, with an average duration of 3.16 seconds and average of 15 spikes per SWD (Fig. 3B, Supplementary Figure 4B). Together, these findings indicate that *Gnb1*^I80T/+^ mice develop robust non-convulsive epileptiform activity characterized by frequent SWDs.

Further ECoG analysis showed that, in addition to epileptiform discharges, *Gnb1*^I80T/+^ mice exhibited persistent abnormalities in background cortical activity throughout the recording period (Fig. 4D-G). To quantify these alterations, power spectrum analysis was performed on artifact-free epochs of quiet wakefulness (see Methods). When analyzed separately, the differences between mutant and WT background activity parameters reached statistical significance in females but not in males (Supplementary Fig. 5), but we note the relatively limited sample size. Because all spectral characteristics of the background activity were not different between males and females, both in WT and *Gnb1*^I80T/+^ mice (Supplementary Figure 5D-F), for the final analysis we pooled data from both sexes (Fig. 3). Mutant mice showed a significant increase in total ECoG power compared with WT controls (Fig. 3E) along with a selective increase in the delta-frequency band, whereas theta, alpha, beta, and gamma band power were unchanged (Fig. 3D,G). Analysis of normalized band powers confirmed that the spectral shift in *Gnb1*^I80T/+^ mice was driven by a significant enrichment of delta-frequency oscillations (Supplementary Figure 6). Consistent with this shift toward delta frequency oscillations, the median frequency was significantly reduced in *Gnb1*^I80T/+^ mice (Fig. 3F).

**Figure 4.**
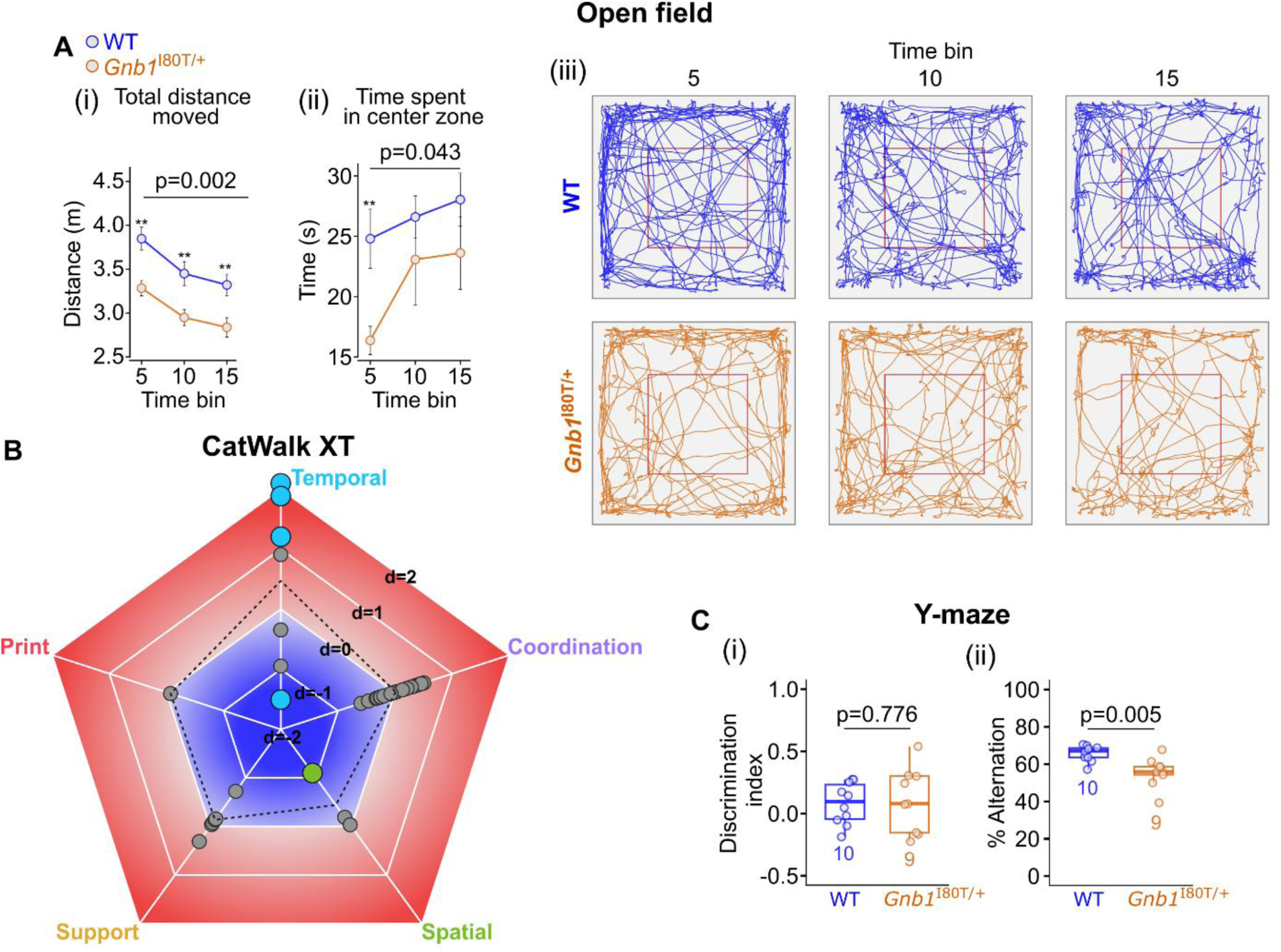
*Gnb1*^I80T/+^ male mice show reduced motor activity and altered gait and open field behavioral parameters. (**A**) Open field test conducted over a 15-minute period: (i) total distance moved, (ii) time spent in the center zone, and (iii) representative traces of WT and *Gnb1*^I80T/+^ mouse locomotion with center zone enclosed in red square. WT: *n* = 15 males; *Gnb1*^I80T/+^: *n* = 13 males; data are shown as mean ± SEM. Statistics: two-way ANOVA with Šídák’s multiple comparisons test. The reduction in total distance over time (5 vs 15 min) was significant in WT (<0.0001) and in *Gnb1*^I80T/+^ (p=0.0049). (**B**) CatWalk XT gait analysis depicted as a pentagon spider plot of Cohen’s d values for all 36 parameters, categorized into five domains (Temporal, Spatial, Coordination, Support, and Print). Each point represents one parameter; points are colored by domain for FDR-significant parameters (FDR < 0.05) and shown in grey for non-significant parameters; the dashed polygon connects the category’s mean Cohen’s d per domain. Statistics: Welch’s t-test or Mann-Whitney U test with Benjamini-Hochberg FDR correction (see Supplementary Figure 8 for individual plots of significant parameters and Supplementary Table 1 for the full parameter list and sample sizes). (**C**) Y-maze test: (i) discrimination index (DI = time spent in novel arm / (time spent in novel arm + time spent in familiar arm)) and (ii) percentage alternation (percentage of successive entries into all three arms without repetition, out of total possible alternations). Boxes represent median and interquartile range; whiskers extend to 1.5× IQR; individual data points represent single animals. WT: *n* = 10 males; *Gnb1*^I80T/+^: *n* = 9 males. Statistics: Welch’s t-test.

Collectively, these findings demonstrate that the *Gnb1*^I80T/+^ variant causes widespread epileptiform encephalographic changes characterized by frequent generalized spike-and-wave discharges, reminiscent of epilepsy observed in GNB1 encephalopathy. This is accompanied by a slowing of background oscillatory cortical activity combined with an increase in total power, as also reported in idiopathic generalized epilepsy (increased delta or delta and theta bands and an increase in total power) (*32–34*).

#### Gnb1I80T/+ male mice display motor and cognitive impairments in early adulthood

To assess behavioral phenotypes of the *Gnb1*^I80T/+^ mouse model, we performed behavioral tests in WT and *Gnb1*^I80T/+^ male and female mice.

The open field test provides a way to systematically assess motivational behavior by quantifying general locomotor activity in a novel environment and visits to the center as an indicator for anxiety-like behavior (*27*). In the open field test, male *Gnb1*^I80T/+^ mice exhibited significantly reduced spontaneous locomotor activity throughout the 15-min recording session (Fig. 4A(i)), similar to the hypokinetic phenotype previously reported in the *Gnb1*^K78R/+^ mouse model (*19*). During the first 5 minutes of exploration, *Gnb1*^I80T/+^ males also spent significantly less time in the center zone, though this difference was not sustained across subsequent recording intervals, suggesting a mild transient anxiety phenotype (*28*) (Fig. 4A(ii)). Notably, *Gnb1*^I80T/+^ females showed no differences in the locomotor activity and time spent in the center zone compared to WT (Supplementary Figure 7A). Furthermore, *Gnb1*^I80T^ females moved significantly greater total distance relative to males (Supplementary Figure 7C).

To examine alterations in motor coordination, we assessed mouse gait using CatWalk XT. Across the 36 parameters tested (Supplementary Table 1), *Gnb1*^I80T/+^ males showed significant impairments in five parameters: four temporal (increased swing duration, increased single stance duration, decreased swing speed, and increased step cycle duration) and one spatial (increased hind paw base of support). Effect sizes for all parameters were quantified as Cohen’s *d* and visualized on a pentagon spider plot, with parameters categorized according to Timotius *et al.* (*29*) (Fig. 4B, details in Supplementary Figure 8A, Supplementary Table 1a). In contrast, *Gnb1*^I80T/+^ females did not show any significant deviations in CatWalk XT tests compared to WT females (Supplementary Figure 8B,C, Supplementary Table 1b).

Gross motor coordination, motor endurance and balance, assessed by the rotarod test, were intact in *Gnb1*^I80T/+^ mice, similar to the findings in *Gnb1*^K78R^ mice (*19*) and similar among males and females (Supplementary Figure 9A). Object memory was assessed by novel object recognition test, and *Gnb1*^I80T/+^ and WT mice showed no difference in preference for the novel object (Supplementary Figure 9B).

Spatial working memory was assessed using the Y-maze alternation test by exploiting the innate tendency of rodents to preferentially explore previously unvisited areas. Impaired alternation indicates a failure to remember the recently visited arm (*30*). *Gnb1*^I80T/+^ male mice displayed significantly reduced spontaneous alternation between maze arms relative to WT (Fig. 4C(ii)), indicating impairment in spatial working memory. Preference for the novel arm itself was not different between *Gnb1*^I80T/+^ and WT males (Fig. 4C(i)). Notably, female *Gnb1*^I80T/+^ mice did not show differences from WT females in Y-maze (Supplementary Figure 9C).

Overall, male *Gnb1*^I80T/+^ mice display a behavioral profile characterized by reduced locomotion in a novel environment, a transient and mild anxiety-like phenotype, indications of impaired spatial working memory, and gait abnormalities, with normal gross motor coordination, balance and object memory. These phenotypical differences between male and female *Gnb1*^I80T/+^ mice imply sex-dependent effects in this mouse model, which remain to be further studied.

### Cross-species transcriptomic analysis: convergent molecular signatures for GNB1 I80T variant

Two iPSC lines were generated from fibroblasts obtained from a patient carrying the heterozygous *GNB1* I80T variant and her healthy sister, who served as a control. Both patient- and control-derived iPSC lines exhibited typical pluripotent stem cell morphology, normal karyotypes, and robust expression of pluripotency markers, as confirmed by immunocytochemistry and flow cytometry (Supplementary Figure 10). The differentiation potential of the iPSC lines into all three germ layers was further confirmed by embryoid body formation and lineage-specific marker expression (Supplementary Figure 10G). Cell line identity was verified by STR profiling, and all lines were negative for mycoplasma contamination.

Control and *GNB1*^I80T^ patient-derived iPSCs were differentiated using an established accelerated cortical differentiation protocol (*35*) and collected on differentiation day 13 for RNA sequencing. Immunocytochemical analysis demonstrated robust expression of the neuronal marker TUJ1 and the early cortical neuronal marker TBR1 in both control and patient-derived cultures (Supplementary Figure 11), confirming successful differentiation toward an early cortical neuronal fate. Both cultures exhibited comparable neuronal morphology and staining patterns, with no obvious differences in differentiation efficiency. According to the original characterization of this differentiation protocol, cultures at this developmental stage are enriched for cortical neurons (∼70–75% TUJ1-positive and ∼60–70% TBR1-positive cells by day 16) and contain minimal glial cells, with remaining cells representing predominantly late neural progenitors and transitioning precursor cells (*35*). These cultures were therefore considered suitable for subsequent transcriptomic analysis. For brevity, we designate these cultures “iPSC-derived neurons”.

To assess transcriptomic changes induced by the GNB1 I80T mutation, we conducted bulk RNA-seq analysis on RNA extracted from hiPSCs and hiPSC-derived cortical neurons from our patient and her healthy sister (as control), and from whole mouse cortex of a 6-week-old *Gnb1*^I80T/+^ female mouse and a littermate WT female mouse as control (Fig. 5A).

**Figure 5.**
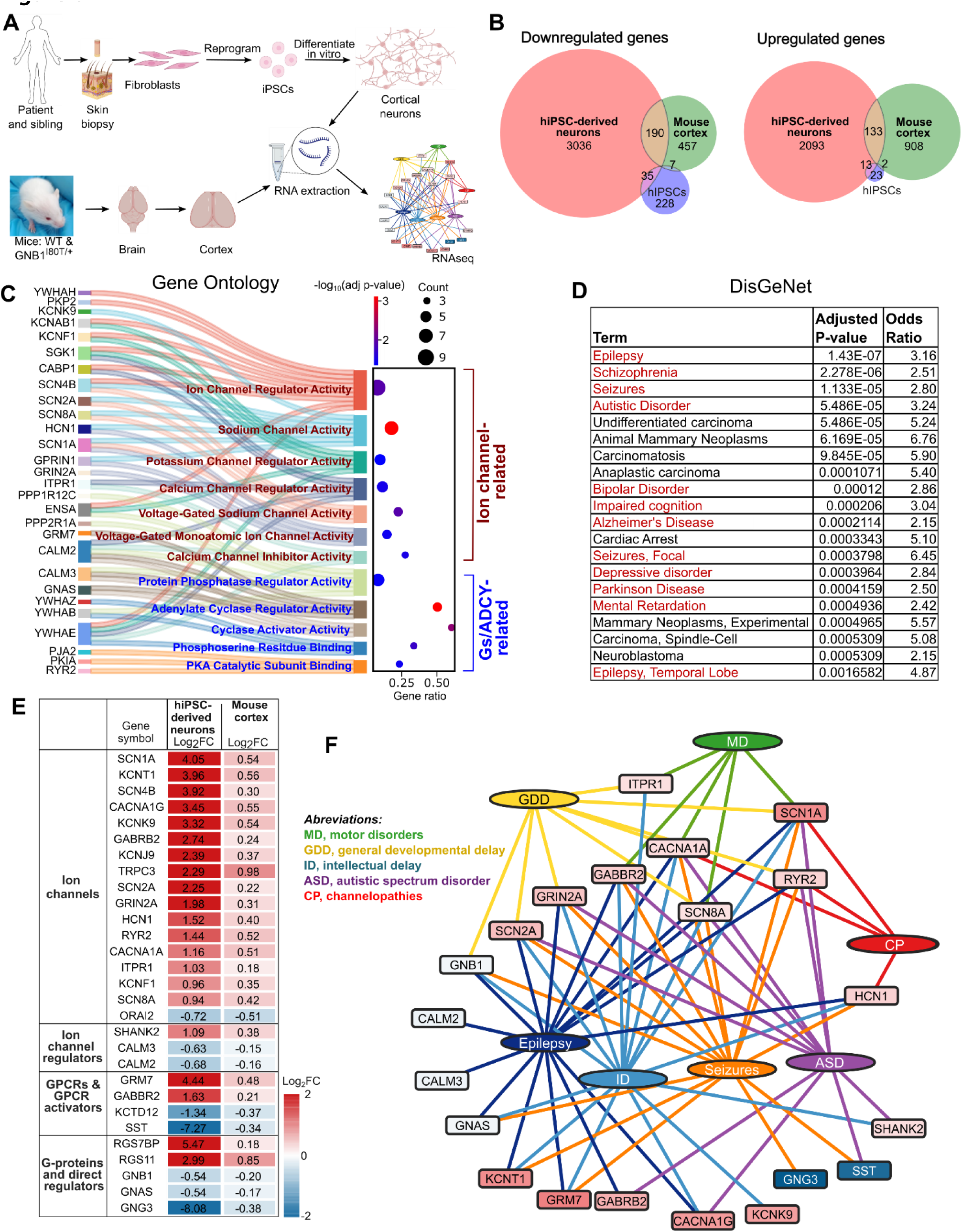
Transcriptomic analysis of differentially expressed genes. **(A)** Scheme of experimental design. (**B**) Venn diagrams (areas are proportional to gene set size) showing significantly overexpressed and underexpressed protein-coding genes (adjusted P-value (FDR) < 0.05) from hiPSC, hiPSC-derived neurons and mouse whole cortex, with intersections representing concordantly dysregulated genes across species and cell types. (**C**) Gene Ontology (Molecular Function) analysis of the 323 concordantly dysregulated genes (the overlapping gene set; Supplementary Table 2) showing all significantly enriched terms with FDR<0.05, split into two categories: ion channel-related (dark red) and Gs/ADCY pathway-related (blue). Genes are linked to each term by Sankey lines. Gene ratio (number of overlapping genes / total genes annotated to the term) is plotted on the right as a dot plot, with dot color representing adjusted p-value and dot size representing gene count. Further details in Supplementary Table 3 and Full result in Data set S4. Plot was produced with SRplot (*74*). (**D**) Top 20 diseases, arranged by FDR (lowest on top), linked to the 323 mouse/human overlapping gene set, identified by DisGeNET overrepresentation analysis (ORA). Full DisGeNet ORA results are in Supplementary Table 4 and in Data Set S5. Dark red font denotes neurological and neurodegenerative disorders. (**E**) The manually curated subset of 29 differentially expressed genes, selected from the 323 mouse/human overlap gene set. Color coding according to log_2_ Fold Change (FC). (**F**) Network diagram of the 29 selected differentially expressed genes, linked to selected disease-associated terms identified by DisGeNET ORA.

Transcriptomic profiling revealed extensive changes in all three sample groups. In patient’s undifferentiated hiPSCs, 389 genes were differentially expressed relative to control hiPSCs, whereas 5,880 were differentially expressed in hiPSC-derived cortical neuronal cultures compared with control cultures (Fig. 5B; Data Sets S1-S3). The substantially greater number of differentially expressed genes (DEGs) and the large magnitude of changes in hiPSC-derived neuronal cultures may reflect their developmental state, corresponding to an early stage of cortical development. At this stage, neurons undergo dynamic transcriptional remodeling driven by neuronal activity, chromatin accessibility, and epigenetic regulation (*36–39*). In comparison, transcriptomic alterations in cortical tissue from *Gnb1*^I80T/+^ mice cortical tissue were less extensive, with 1,736 DEGs relative to the WT control (Fig. 5B; Data Sets S1-S3). This number lies at the upper end of the range reported for transcriptomic studies of mouse models of developmental and epileptic encephalopathies caused by haploinsufficiency or pathogenic variants in voltage-gated sodium channel genes, where the number of DEGs ranges from several tens (*40*) to approximately 1,500 (*41*). These findings indicate that the GNB1E-associated I80T variant is accompanied by widespread transcriptional alterations. Notably, no genes were concordantly differentially expressed across all three sample types, i.e. between undifferentiated hiPSCs and the two neuron-enriched populations (Fig. 5B), indicating that the transcriptional consequences of the I80T variant are highly dependent on cell type and differentiation state.

Downstream analyses therefore focused on hiPSC-derived neurons and mouse cortical tissue, the two datasets most relevant to the neuronal phenotype. We conducted a comparative cross-species analysis to try to identify shared transcriptomic changes relevant to convergent pathological outcomes. 323 differentially expressed genes showed concordant changes (either over- or underexpressed) in mouse cortex and human neurons (hereafter termed 323 overlapping gene set; Fig. 5B, Supplementary Table 2). Notably, GNB1 itself was downregulated in both systems (31% decrease in hiPSC-derived neurons and 21% in mouse cortex; Fig. 5E), suggesting that the I80T variant reduces GNB1 transcript abundance *in vivo* and *in vitro*.

We applied two hypothesis-free approaches to analyze the 323 overlapping gene set. First, Gene Ontology (GO) overrepresentation analysis in Molecular Function domain identified 12 significantly enriched terms (FDR < 0.05; Fig. 5C, Supplementary Table 3, Data Set 4). Remarkably, we noticed that these terms can be segregated into two clear functional clusters. The first comprised ion channel-related terms, including sodium channel activity, ion channel regulator activity, voltage-gated sodium channel activity, calcium channel inhibitor activity, calcium channel regulator activity, voltage-gated monoatomic ion channel activity, and potassium channel regulator activity. The second group comprised genes linked to all stages of the G_s_/adenylyl cyclase (ADCY)-protein kinase A (PKA) cascade (adenylate cyclase regulator activity, cyclase activator activity, phosphoserine residue binding, PKA subunit binding), including the related termination step (protein phosphatase regulator activity). This two-cluster enrichment pattern strongly implicates the dysregulation of ion channel-regulated neuronal excitability and cAMP/PKA signaling as key molecular consequences of the GNB1 I80T mutation.

Second, disease overrepresentation analysis using the DisGeNET gene-disease association library on the overlapping gene set identified 154 significantly enriched disease terms (FDR < 0.05; Supplementary Table 4, Data Set 5). The top 20 enriched terms (Fig. 5D) recapitulating many of the key clinical features of GNB1 encephalopathy including epilepsy, seizures, global developmental delay and intellectual disability, as well as several neurological and neuropsychiatric disorders, supporting the neuronal origin. Cancer-linked genes constituted another significantly enriched group of disease terms, in line with the carcinogenic potential reported for somatic GNB1 I80T mutations (*1*).

To integrate these findings into a targeted gene set, we manually curated the 323 overlapping gene set to include all ion channels, all G protein subunits, all GPCRs and some prominent GPCR direct regulators, and some selected ion channel regulators in the 323 overlapping gene set. Overall, 29 genes were shortlisted comprising 17 ion channels, 3 ion channel regulators, 4 G protein-coupled receptors (GPCRs) and GPCR regulators, and 5 G protein subunits. Notably, among G protein subunits the only Gα that was found in the 323 gene overlapping set was Gα_s_, supporting the importance of G_s_/ADCY/cAMP/PKA pathway (Fig. 5E). The direction and magnitude of expression changes across the human and mouse samples are displayed as a heatmap (Fig. 5E). These 29 genes were further visualized in a network diagram along with the top neurological terms from the DisGeNET analysis (Fig. 5F), highlighting that the shared cross-species transcriptional signature is substantially enriched for genes implicated in neurodevelopmental and neuropsychiatric disorders, including several genes linked to epileptic encephalopathies.

## Discussion

In this study, we sought to define the molecular mechanisms underlying the clinical manifestations of GNB1 Encephalopathy, focusing on its most prevalent and clinically representative variant, I80T. To this end, we combined longitudinal clinical characterization of a patient carrying the I80T variant with phenotyping characterization of a CRISPR/Cas9-generated *Gnb1*^I80T/+^ mouse and cross-species transcriptomic analysis of patient-derived cortical neurons and mouse cortex. The mouse recapitulated key disease features, including early developmental delay, motor abnormalities, and robust epileptiform activity, while transcriptomic analyses converged on two principal molecular pathways: dysregulation of ion-channel function and G_s_/ADCY/cAMP/PKA signaling. These findings establish the *Gnb1*^I80T/+^ mouse model as an adequate platform for future mechanistic and therapeutic studies, extend previous evidence implicating Gβγ-dependent ion-channel dysfunction, and identify dysregulated ADCY/cAMP/PKA signaling as an additional pathway that may contribute to disease pathogenesis.

The patient’s disease history highlights the evolving nature of GNB1E. Early hypotonia and developmental delay progressed to spasticity, dystonia, bradykinesia, and severe motor disability, while epilepsy emerged later with spike-wave activation during sleep (SWAS). This progression is consistent with previous reports but remains poorly defined across GNB1 variants (*4, 5, 42–44*). Importantly, developmental impairment preceded epileptiform activity, suggesting that epilepsy is unlikely to be the primary cause of neurodevelopmental dysfunction, although prolonged epileptiform discharges may have further affected cognitive and language development (*45–47*). The limited response to conventional antiseizure medications further underscores the need for mechanism-based therapies.

Interestingly, this clinical trajectory was reflected in the *Gnb1*^I80T/+^ mouse. Mutant pups displayed impaired growth and delayed sensorimotor maturation, paralleling the patient’s early developmental abnormalities. Adult behavioral deficits were comparatively mild and largely restricted to males, with reduced locomotion, gait abnormalities, and impaired working memory. Although the model does not reproduce the severity of the patient’s movement disorder, it captures key aspects of disease biology.

The most robust phenotype in the mouse was cortical network dysfunction characterized by frequent bilateral spike-and-wave discharges (SWDs), increased cortical power, and slowing of background oscillations. Although behavioral correlates of individual SWDs were not examined, these abnormalities closely resemble the absence-like phenotype reported in the *Gnb1*^K78R/+^ model (*19, 31*). While the patient’s EEG cannot be directly compared with mouse electrocorticography because of differences in species, developmental stage, and recording methodology, both independently demonstrate substantial disturbances of cortical excitability. These parallels support altered network activity as a defining feature of the I80T variant and provide a physiological link between the experimental model and the patient’s refractory epileptic phenotype.

Despite producing different effects on GIRK channel regulation *in vitro*, both the I80T and K78R variants converge on remarkably similar epileptiform phenotypes *in vivo* (*18*). This convergence is consistent with the concept that both gain- and loss-of-function alterations in ion-channel signaling can disrupt excitation-inhibition balance and produce epilepsy (*48–50*). Thus, distinct molecular defects may ultimately converge on common pathological network states, an important consideration for therapeutic development toward genetically heterogeneous forms of GNB1E. (However, we cannot exclude the possibility that the shared phenotype also reflects a common effect of both variants on an as yet uncharacterized Gβγ target).

Cross-species transcriptomic analysis independently reinforced these physiological findings. Although patient-derived cortical neurons and mouse cortex represent distinct experimental systems, both demonstrated concordant dysregulation of 323 genes enriched for ion-channel function and neurological disease. Notably, several genes, including SCN1A, SCN2A, SCN8A, HCN1, KCNT1, GRIN2A, and CACNA1A are established causes of epilepsy and developmental encephalopathy. This conserved molecular signature closely mirrors the patient’s pronounced epilepsy and progressive movement disorder, supporting altered neuronal excitability as an important mechanism underlying GNB1E.

These findings also extend our previous work implicating GIRK channels in GNB1E. Gβγ directly regulates GIRK channels and modulates multiple ion-channel families through direct and indirect mechanisms (*6, 51*). Pharmacological targeting of GIRK signaling suppressed SWDs in the *Gnb1*^K78R/+^ mouse (*18, 19*), whereas the present transcriptomic data suggest that I80T causes broader remodeling of excitability-related pathways (though there is one GIRK gene, KCNJ9 (GIRK3) in our 323 overlapping gene set; Fig. 5E). Because many of the dysregulated ion channels are regulated by GPCR signaling and phosphorylation, altered Gβ1 function is likely to influence neuronal excitability through multiple interconnected mechanisms rather than a single downstream effector.

A second major finding was enrichment of genes involved in Gs/ADCY/cAMP/PKA signaling. Although this pathway has received little attention in GNB1E, it is highly relevant given the central role of heterotrimeric G proteins in regulating adenylyl cyclase activity. While transcriptomic data cannot establish pathway directionality, coordinated changes in GPCRs, G-protein subunits, and downstream signaling components suggest widespread alteration of cAMP-dependent signaling. This molecular signature is particularly compelling in the context of the patient’s prominent dystonia and bradykinesia, as disruption of ADCY5-mediated cAMP signaling causes childhood-onset movement disorders with overlapping clinical features (*52–54*). Future biochemical studies measuring cAMP levels, PKA activity, and downstream phosphorylation will be essential to define the functional consequences of these changes.

Several limitations warrant consideration. Human transcriptomic analyses were derived from a single patient and sibling control, requiring validation in additional patient-derived lines. Likewise, both differentiated neuronal cultures and bulk mouse cortex contain heterogeneous cell populations that may obscure cell-type-specific effects (*55–57*). Single-cell transcriptomics, use of isogenic hiPSC cell lines for control, and biochemical studies will therefore be important for refining these molecular mechanisms, while further behavioral-electrophysiological studies are needed to better define the epileptic phenotype and underlying circuits.

Despite these limitations, our integrated clinical, experimental, and transcriptomic approach provides convergent evidence that altered neuronal excitability is a central feature of GNB1 I80T encephalopathy. Beyond reinforcing the importance of Gβγ-dependent ion-channel regulation, our findings identify ADCY/cAMP/PKA signaling as a previously underappreciated mechanistic axis and establish the *Gnb1*^I80T/+^ mouse as a valuable platform for testing targeted therapeutic strategies. The correspondence between the patient’s evolving neurological phenotype, cortical network dysfunction in the mouse, and the conserved cross-species transcriptional signature strengthens the biological relevance of the identified pathways.

## Materials and Methods

### Human electrophysiological data

Clinical electrophysiological data were obtained from the patient in the form of electroencephalography (EEG) recordings utilizing 10-20 electrode placement system on MicroMed device (Natus, Middleton, WI 53562, USA). Sampling frequency was 256 Hz. Presentation of data was as follows: EEG shown in Fig. 1B,C is the original unprocessed recording from the indicated channels. Electrode configuration in EEG analysis in Supplementary Fig. 1 was done using standard longitudinal bipolar montage (a “banana” montage) (*58*). Electrical difference between two neighboring electrodes arranged in sequential, front-to-back lines (left parasagittal, right parasagittal, left temporal, right temporal chains) are displayed. EEG analysis was done by board certified pediatric neurologist (H.B.).

### Mice

#### Generation of the Gnb1^I80T/+^ mice

*Gnb1^I80T/+^* mice were generated using CRISPR/Cas9 at The Jackson Laboratory (JAX; Bar Harbor, Maine, USA) on a C57BL/6NJ background (JAX stock #005304). A missense mutation was introduced into mouse *Gnb1* to replace isoleucine with threonine at amino acid position 80 (I80T; ATC > ACC). CRISPR guides targeting Gnb1 exon 6 were used together with a single-stranded DNA donor carrying the I80T substitution and a silent guide-blocking mutation (ATC > ATT) four base pairs downstream. Founder animals were screened by PCR and Sanger sequencing, and sequence-confirmed founders were bred to generate N1 animals. Five N1 mice carrying the heterozygous Gnb1-I80T knock-in allele were obtained and used to establish the colony. According to the report from JAX, the heterozygote *Gnb1^I80T/+^* mice on the C57BL/6NJ background showed low viability, therefore we decided to shift the genetic background to FVB. To establish an FVB background that we consider as pure, the original C57BL/6NJ line was crossed with FVB mice (Envigo, Israel) for 8+ generations. For all experiments, mice from generations F8-F10 were used. Wild-type (WT) littermates served as controls. Mice were housed in ventilated cages under controlled environmental conditions (22–23°C, ∼60% humidity, 12 h light:12 h dark cycle) with ad libitum access to standard chow and water. All animal experiments were approved by the Tel Aviv University Institutional Animal Care and Use Committee (Approval number TAU-MD-IL-2211-167-4).

#### Genotyping of mice

DNA was extracted from the tail or ear tissue using the HotShot method (*59*). PCR amplification was performed using the following primers: forward, TTTATCTTTTCTCCAGGCTCCTT; reverse, AGCACAACTGTATGCCAAGC. PCR generated a 200 bp amplicon. Sanger sequencing was subsequently performed to distinguish WT from *Gnb*1^I80T/+^ mice.

#### Developmental milestones

Pups were tattooed on the tail for identification at postnatal day 3 (P3) (*60*). Developmental milestone assessments, including body weight, body length, righting reflex, and negative geotaxis, were conducted on alternating days to minimize handling-related stress (*61*).

Body weight and body length were recorded for each pup. The righting reflex was assessed as the latency required for a pup placed in a supine position to right itself onto all four limbs. Negative geotaxis was evaluated on a cotton-lined cardboard incline as the latency required to rotate 180° from an initial downward-facing position. For each testing session, pups were gently removed individually from the nest and placed on a clean bench protector. The cage lid was immediately replaced to minimize disturbance to the remaining litter. All assessments were completed by a trained experimenter within 3 min per pup, after which pups were promptly returned to the nest.

### Behavioral experiments

All behavioral experiments were performed at The Myers Neuro-Behavioral Core Facility, Tel Aviv University. All behavioral tests were conducted on 6- to 8-week-old mice. Mice were divided into two experimental groups: wild-type (WT) and heterozygous (Het) animals. All mice underwent behavioral testing in the following order: open field (1 day), novel object recognition (2 days), rotarod (1 day), CatWalk XT (1 day), and Y-maze (1 day). As the Myers Neuro-Behavioral Core Facility operates on a reversed light-dark cycle relative to the breeding and weaning facility, mice were transferred to the behavioral core at least one week prior to testing to allow acclimatization. Before each experimental session, mice were habituated to the testing room for a minimum of 30 minutes. A minimum inter-assay interval of 24 h without human interaction was maintained between assays. All behavioral testing was conducted during the dark period between 11:00 AM and 2:00 PM and performed on one mouse at a time. All equipment was thoroughly cleaned with Virusolve+ before and between trials. Except for the rotarod assay, behavior during all tests was recorded using monochrome cameras (Basler acA1300-60gm, Basler AG, Ahrensburg, Germany). Video recordings were analyzed using EthoVision XT 18 software (Noldus Information Technology, Wageningen, The Netherlands) (*62*).

#### Open field

In the open field test, mice were placed in one of the corners of an open-top plexiglass box (L×W: 50×50 cm), and behavior was recorded for 15 min. Activity, defined as the fraction of pixels that changed in the entire arena between consecutive frames (frame rate, 25 Hz), was measured throughout the run duration to evaluate exploratory behavior. The total distance traveled (for general locomotor activity), and the time spent in the center of the arena (for anxiety-like behavior) were measured (*63*).

#### Novel object recognition

The novel object recognition (NOR) test was used to assess visual recognition memory based on rodents’ innate tendency to preferentially explore novel objects over familiar ones (*64*). The NOR paradigm was conducted on the two consecutive days following the open field test, utilizing the same arena. During each stage, mice were placed individually in the center of the arena and allowed to explore freely for 5 min. Twenty-four hours after the open field trial, mice underwent the learning (familiarization) trial, in which two identical familiar objects (tin cola cans) were placed symmetrically in the arena. After an additional 24 h interval, mice underwent the test trial, during which one of the familiar objects was replaced with a novel object (a ceramic coffee mug). Object exploration was recorded, and recognition memory performance was quantified using a discrimination index, calculated as:

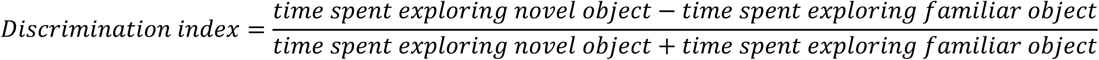

Higher discrimination index values indicate greater preference for the novel object and improved recognition memory. Total exploration time of both objects was also recorded and analyzed to ensure adequate object interaction and to control for potential differences in exploration between animals.

#### Rotarod

Motor coordination and motor learning were evaluated (*65*) using an accelerating rotarod apparatus (Ugo Basile Srl, Gemonio, Italy). Mice were placed on the rod and underwent five consecutive trials in a single session with an inter-trial interval of approximately 1 minute. Each trial began at 4 rpm and accelerated to a maximum of 40 rpm over 5 minutes. The latency to fall from the rod was measured.

#### Catwalk

The gait analysis apparatus (CatWalk XT, Noldus Information Technology, Wageningen, The Netherlands) enables the assessment of voluntary gait and locomotion in mice (*66*). The apparatus consists of a glass walkway (130 × 20 cm), a GigE video camera, and dedicated analysis software for footprint quantification. The walkway is illuminated from above with red light and from the side with green light. Green light undergoes internal reflection within the glass except at paw contact points, enabling footprint visualization. The walkway led to a dark goal box and was enclosed by an adjustable tunnel. During testing, mice were placed at the entrance and allowed to traverse the walkway voluntarily. A compliant run was defined as a crossing with a maximum run duration of 5 seconds and a maximum run variation of 50%. Each mouse completed three compliant runs, and the average value across runs was used for analysis. Between mice, the walkway was cleaned with water and thoroughly wiped dry to remove all residue.

For statistical analysis, paw-specific parameters, including stride length, swing speed, step cycle, stance duration, swing duration, single stance duration, duty cycle, and print area, were averaged across all four paws to yield one representative value per parameter per animal. Additional parameters measuring coordination, stability, and running speed were included, yielding a total of 36 parameters for analysis (full list provided in Supplementary Table 1). Outliers were removed using a 2×IQR criterion. Based on Shapiro-Wilk normality test results, either Welch’s unpaired t-test or Mann-Whitney U test was performed, with multiple testing correction applied using the Benjamini-Hochberg false discovery rate (FDR) method.

#### Y maze

The Y-maze test was used to assess spontaneous exploratory behavior, responsiveness to novelty, and spatial memory function in mice (*67*). The task is based on rodents’ innate preference to explore previously unvisited environments. The apparatus consisted of three identical arms (8 × 30 × 15 cm) arranged at 120° angles. The test comprised two stages separated by a 5-min inter-trial interval. In each stage, mice were individually placed at the edge of a designated start arm. During the first stage (training trial), one arm of the maze was blocked, and mice were allowed to explore the remaining two arms for 5 min. Following this trial, mice were returned to their home cages for 5 min while the maze was cleaned with virusolve and the blocked arm was reopened. During the second stage (test trial), mice were reintroduced to the maze and allowed to explore all three arms freely for 5 min. Time spent in each arm was recorded.

Spatial memory performance was quantified using two measures. First, a discrimination preference index, calculated as:

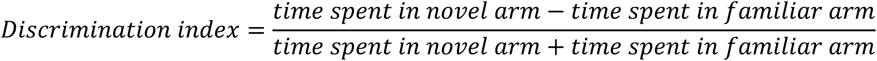

Higher preference index values indicate greater exploration of the novel arm and improved spatial recognition memory. Second, spontaneous alternation was calculated as the percentage of alternations out of the maximum possible alternations:

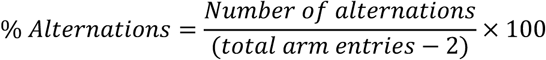

where an alternation was defined as a consecutive visit to all three arms without revisiting the same arm, and the maximum number of possible alternations was calculated by subtracting 2 (the number of arms minus one) from the total number of arm entries. Higher alternation scores indicate greater spontaneous exploration and working memory performance.

### Electrocorticography

#### Electrode implantation

Electrode implantation, recordings and analyses were performed as previously described (*68*). Briefly, mice were anesthetized with ketamine (100 mg/kg), xylazine (10 mg/kg), and carprofen (5 mg/kg) for analgesia. A midline incision was made over the skull, and small burr holes were manually drilled over visually identified regions of the somatosensory cortex and cerebellum using a 30-gauge needle. Fine silver wire electrodes (130 μm bare diameter; 180 μm coated diameter) were implanted bilaterally over visually identified regions of the somatosensory cortex. A reference electrode was positioned over the cerebellum, and a ground electrode was placed subcutaneously behind the neck toward the left shoulder. An EMG electrode was placed subcutaneously over the right shoulder. The electrode connector was secured using dental cement. Mice were allowed a minimum recovery period of 48 hours before recordings commenced.

Video-ECoG recordings (2–3 h duration) were obtained from freely behaving mice during the light cycle between 08:00 and 18:00. Mice were connected to a T8 Headstage (Triangle BioSystems International, Durham, NC, USA), and signals were acquired using PowerLab 8/35 hardware and LabChart 8 software (ADInstruments, Sydney, Australia). Electrical signals were digitized at a sampling rate of 1 kHz. Prior to analysis, both left and right somatosensory cortex signals were referenced by subtracting the cerebellar signal.

#### Spike-Wave Discharge Detection and Analysis

Spike-wave discharge (SWD) detection and analysis were performed on the right somatosensory cortex channel (cerebellar-referenced) using a custom Python-based pipeline applied to ECoG signals bandpass filtered between 7 and 50 Hz. The full pipeline is publicly available on GitHub (https://github.com/VigneshwarRanjan/Spike-Wave-Discharge-Detection-Pipeline-for-Mouse-ECoG). The analysis has been performed using a standard moving average detection procedure. Spike and SWD selection parameters that rendered optimal spike selection with lowest fraction of false events were determined by repetitive comparison of analysis results of representative EEG recordings with the automatic routine versus visual selection of SWDs. The analysis proceeded in four steps: (1) candidate spikes were identified using a sliding z-score algorithm, in which for each data point a z-score was computed relative to the mean and standard deviation of the preceding 3-second window, and deflections exceeding a threshold of z < −4 were flagged as candidate spikes; (2) selection of bursts was done as follows: flagged spikes were analyzed for burst structure based on inter-spike interval (ISI), with three or more consecutive spikes with ISIs ≤ 0.5 s classified as a burst; (3) all candidate SWDs were presented in a custom interactive graphical user interface (GUI) for manual verification, in which each event was visually inspected and classified as a true positive, false positive, or duplicate detection; and (4) all verified SWDs underwent a second manual pass using a dedicated GUI, in which SWD duration was measured by manually placing start and end markers on each event, and spike count was quantified by setting an amplitude threshold, with peaks detected using the find_peaks function from the SciPy signal processing library https://docs.scipy.org/doc/scipy/reference/generated/scipy.signal.find_peaks.html. False negatives (SWDs missed by the automated detection) identified during visual inspection were manually added to the dataset.

#### Background ECoG Analysis

For background frequency band analysis, offline signal processing and analysis were performed using LabChart 8 software (ADInstruments, Castle Hill, Australia). ECoG signals were bandpass filtered between 0.5 and 100 Hz, while EMG signals were processed using a 0.5 Hz high-pass filter. For each recording, five to ten artifact-free 30-second segments resembling quiet wakefulness were selected from both left and right somatosensory cortex channels (each cerebellar-referenced), beginning 0–30 seconds after a period of movement. Both the preceding movement and the subsequent absence of movement within the selected segments were verified using the EMG signal and video recordings, to exclude movement-related artifacts. Power spectral density (PSD) was calculated utilizing LabChart software for each segment using a fast Fourier transform (FFT) with an FFT size of 1,024 points and a Hann (cosine-bell) window with 50% overlap. PSDs were first averaged across segments within each channel, then averaged across the left and right channels to yield a single representative power spectrum per recording. Frequency bands were defined as: delta, 0.9–3.9 Hz; theta, 4.8–7.8 Hz; alpha, 8.7–11.7 Hz; beta, 12.6–29.2 Hz; gamma, 30.2–99.6 Hz.

### Generation and Characterization of Patient-Derived iPSC Lines

The use of human tissue samples for generation of iPSCs and the subsequent analysis was approved by the Haim Sheba Medical Center Ethics (Helsinki) Committee for Human Medicine #6158-19-SMC, and by the patient’s legal guardians (parents) who signed the appropriate Informed Consent statement.

#### Fibroblast Isolation, reprogramming and iPSC generation

Skin punch biopsies (3 mm) were obtained under local anesthesia and with informed consent from the patient carrying the GNB1 variant (heterozygous) and her healthy sister. Biopsies were placed in 6-well plates precoated with 1% gelatin (Sigma) and cultured in DMEM supplemented with 20% fetal bovine serum (FBS) at 37°C and 5% CO₂. Fibroblasts migrating from tissue explants were expanded, passaged weekly, and cryopreserved in liquid nitrogen.

Fibroblasts were expanded to confluency in DMEM containing 15% FBS, dissociated using TrypLE (Gibco), and electroporated (0.5–1 × 10^6 cells) with non-integrating episomal vectors using the Neon Transfection System (Invitrogen). Cells were plated onto mouse embryonic fibroblast (MEF)-coated plates and cultured in DMEM supplemented with 15% FBS and 5 ng/mL basic fibroblast growth factor (bFGF; Thermo Fisher Scientific), with media changes every other day. From day 11 onward, cultures were maintained in NutriStem medium (Biological Industries, Beit-Haemek, Israel) supplemented with 5 ng/mL bFGF. Emerging colonies were manually selected on day 21, transferred to fresh MEF-coated plates for expansion, and subsequently adapted to Matrigel-coated plates for characterization.

#### Immunocytochemistry and flow cytometry analysis

iPSCs were fixed in 4% paraformaldehyde for 20 min at room temperature and washed with DPBS. Cells were blocked in 1% bovine serum albumin (BSA), with 0.1% Triton X-100 added for intracellular staining. Primary antibodies (Table: x) were applied overnight at 4°C, followed by incubation with fluorescent secondary antibodies for 1 h at room temperature. Nuclei were counterstained with DAPI, and images were acquired using an EVOS XL imaging system (Thermo Fisher Scientific).

Expression of pluripotency markers was quantified by fluorescence-activated cell sorting (FACS). Cells were dissociated using TrypLE and washed with DPBS. For intracellular staining, cells were fixed and permeabilized using commercial fixation/permeabilization reagents (Invitrogen). For surface marker analysis, cells were incubated with primary antibodies in 3% FBS/DPBS for 40 min at 4°C. Following washing, cells were incubated with fluorescent secondary antibodies and analyzed using a NovoCyte flow cytometer (Agilent Technologies, Santa Clara, California). Antibodies used are listed in Supplementary Table 5.

#### Differentiation potential and karyotyping

To assess pluripotency, iPSCs were dissociated into single cells and cultured in suspension in NutriStem medium supplemented with 10 ng/mL bFGF and 7 μM ROCK inhibitor (Enzo Life Sciences, Farmingdale, NY) to allow spontaneous embryoid body (EB) formation. EBs were subsequently maintained in differentiation medium (DMEM supplemented with 15% FBS, 1% non-essential amino acids, and 0.1 mM β-mercaptoethanol) for 4-7 days and then transferred onto 0.1% gelatin-coated plates. Cultures were maintained for 21 days with medium replacement twice weekly and evaluated for differentiation into the three germ layers using lineage-specific immunostaining Supplementary Table 5.

Pluripotency was validated by immunocytochemistry (ICC) demonstrating expression of the canonical pluripotency markers TRA-1-60, SSEA-4, OCT4, SOX2, and NANOG (Supplementary Figure 10C,D). Quantitative assessment by fluorescence-activated cell sorting (FACS) further confirmed expression of these markers in 71–99% of cells across the generated iPSC lines. To evaluate differentiation potential, iPSC clones were subjected to embryoid body (EB) formation followed by spontaneous differentiation into the three germ layers. Lineage commitment was validated by ICC using established lineage-specific markers: 68-kDa neurofilament (NF-L) for ectoderm, α-smooth muscle actin (α-SMA) for mesoderm, and α-fetoprotein (AFP) for endoderm. Cell line identity was confirmed by short tandem repeat (STR) profiling through comparison of parental fibroblasts and their corresponding derived iPSC lines, using a panel of standard STR loci. All iPSC lines were routinely tested for mycoplasma contamination using the Hy-Mycoplasma PCR kit (Hylabs) and confirmed to be negative.

Chromosomal integrity was evaluated by G-banding analysis. iPSCs were treated with colcemid (100 ng/mL; Biological Industries), harvested using TrypLE, fixed in methanol:acetic acid (3:1), and processed for cytogenetic analysis.

### Generation and Characterization of Patient-iPSC derived cortical neurons

We followed the cortical differentiation protocol by (*35*). For neural differentiation, cells were disassociated with Accutase and pre-plated as reported at the density of 200,000 cells/cm2 supplemented with 10 μM Y-27632 on matrigel coated plates, and started differentiation the next day when confluent. KSR medium which contained 820 ml of Knockout DMEM, 150 ml Knockout Serum Replacement, 1 mM L-glutamine, 100 μM MEM nonessential amino acids and 0.1 mM β-mercaptoethanol was used to start differentiation. Inhibitors used in LSB+X/P/S/D induction included LDN193189 (250 nM; Stemgent), SB431542 (10 μM; Tocris), XAV939 (5 μM; Tocris), PD0325901 (1 μM in P1S5D; Tocris), SU5402 (5 μM in P1S5D; Biovision), DAPT (10 μM; Tocris). N2 medium with B27 supplement (N2/B27; Life Technologies) was added in increasing 1/3 increment every other day from day 4, until reaching 100% neurobasal/B27/L-Glu containing medium (NB/B27; Life Technologies) supplemented with BDNF (20 ng/ml; R&D), dibutyryl cAMP (0.5 mM; Sigma-Aldrich) and ascorbic acid (0.2 mM; Sigma-Aldrich) (BCA) at day 8. LSB+X were added from day 0-6, and P/S/D were added from day 2-13.

#### Quality Control

Immunostaining with TBR1 (preplate, subplate and layer VI) and TUJ1 as markers of early born cortical neurons, in LSB+X/P/S/D conditions. By 13 days of differentiation, both control and iPSC-core control lines were enriched for TBR1+/TUJ1+ neurons (Supplementary Figure 11).

### RNA Isolation, RNA-Seq libraries and transcriptomic analysis

Total RNA was isolated from hiPSC-derived cortical neurons and mouse cortical tissue using a combined TRIzol-column purification protocol. Briefly, cortical tissue was homogenized in TRIzol Reagent (Invitrogen #15596026). Following phase separation and RNA extraction, samples were further purified using the RNeasy mini kit (Qiagen #74134) to improve RNA purity and remove residual contaminants. RNA concentration and purity were determined using a NanoDrop spectrophotometer, and RNA integrity was evaluated by bleach agarose gel electrophoresis (*69*).

For RNA-seq analysis, we prepared triplicates from RNA samples from hiPSCs and hiPSC-derived cortical neurons from our patient and her healthy sister, from whole mouse cortex of a 6-week-old *Gnb1*^I80T/+^ female mouse and a littermate WT female mouse (18 samples total). Raw reads were processed by adapter trimming and low-quality base removal using fastp (v0.20.1) (*70*), followed by mapping and quantification using STAR (v2.7.2a) (*71*). Downstream analyses were carried out in R. Sample-level quality control, aided by principal component analysis (PCA), identified one sample (neuron_I80T_1) as a clear outlier, which was therefore excluded from all subsequent analyses (Supplementary Figure 12). Count normalization and differential expression analysis were performed using DESeq2 (v1.40.2) (*72*), with p-values corrected for multiple testing using the Benjamini–Hochberg false discovery rate (FDR) method; genes with FDR < 0.05 were considered significantly differentially expressed.

For disease enrichment analysis, genes concordantly and significantly (FDR < 0.05) differentially expressed in both hiPSC-derived neurons and mouse cortical samples were identified, yielding a total of 323 genes (Supplementary Table 2). This dataset was then used for overrepresentation analysis using Enrichr (*73*) with the DisGeNet library to identify significantly enriched disease-associated gene sets.

### Figures and graphics

Figures were produced using InkScape freeware (www.inkscape.org). Images in Figures 1A and 5A were produced with BioRender, upon obtaining the appropriate licenses.

### Statistical analysis

Initial sample-size estimation for initial trials was performed based on previous experience, followed by more accurate estimates using G*Power software (version 3.1) based on effect sizes derived from preliminary experiments. Randomization was not performed, except for developmental milestones assays, as group assignment was determined by PCR-based genotyping prior to testing and was therefore defined by animal’s genotype (WT or *Gnb1*^I80T/+^) rather than arbitrary allocation.

Data are presented as mean ± standard error of mean (SEM). Outliers were removed using ROUT method (Q = 1%). Statistical analyses were performed using GraphPad Prism 10, and details of statistical tests and corresponding *P*-values for individual panels are provided in figures or figure legends. Comparisons between WT and mutant mice were generally performed using an unpaired two-tailed Student’s *t*-test for normally distributed data and the Mann–Whitney *U* test for non-normally distributed data. Statistical significance was defined as *P* < 0.05.

## Acknowledgments

We thank Prof. Miguel Chilon Rodriguez (all d’Hebron Institute of Research / Universitat Autonoma de Barcelona, Spain) for critical reading of the manuscript, and Prof. Wayne Frankel (Columbia University, New York, USA) for advice on mouse model.

## Authors contributions

Conceptualization: H.P.R., V.R., M.K., N.D

Data curation: H.P.R, V.R, G.S, H.B, G.H, B.B.H, G.H, T. R, Y.Y, L.B

Formal analyses: H.P.R, V.R, G.S, G.H, T. R, Y.Y, L.B, N.S, D.Y, M.R, N.D

Investigation: H.P.R, V.R, G.S, H.B, G.H, T. R, G.D.V, Y.Y, L.B

Methodology: H.P.R, V.R, G.S, H.B, G.H, T. R, G.D.V, Y.Y, L.B, N.S, D.Y, M.R, N.D

Project administration: N.D. Supervision: M.R., L.B., N.S., N.D.

Writing – original draft: H.P.R, V.R., H.B.

Writing – review & editing: H.P.R, V.R., M.K., G.S, H.B., G.H., B.B-Z., T.R., G.D.V., Y.Y., L.B., B.M.M., L.B., N.S., D.Y., M.R., N.D.

## Data sharing

All reported data are presented in Results and Supplementary Materials. Analysis routines are freely available on GitHub: https://github.com/VigneshwarRanjan/Spike-Wave-Discharge-Detection-Pipeline-for-Mouse-ECoG. Supplementary data sets are at https://drive.google.com/drive/folders/1D-W3LL0gSn7yWZ8h1ynLq-y29KaCLDea?usp=sharing. Original recordings and summary files will be made available upon reasonable request, excluding the human patient data.

