## Supplementary material for "Cross-species analysis of GNB1 I80T encephalopathy: conserved developmental, epileptic and neuronal transcriptome signatures": Suplementary Material

#### Supplementary Figures

##### *Supplementary Figure 1*

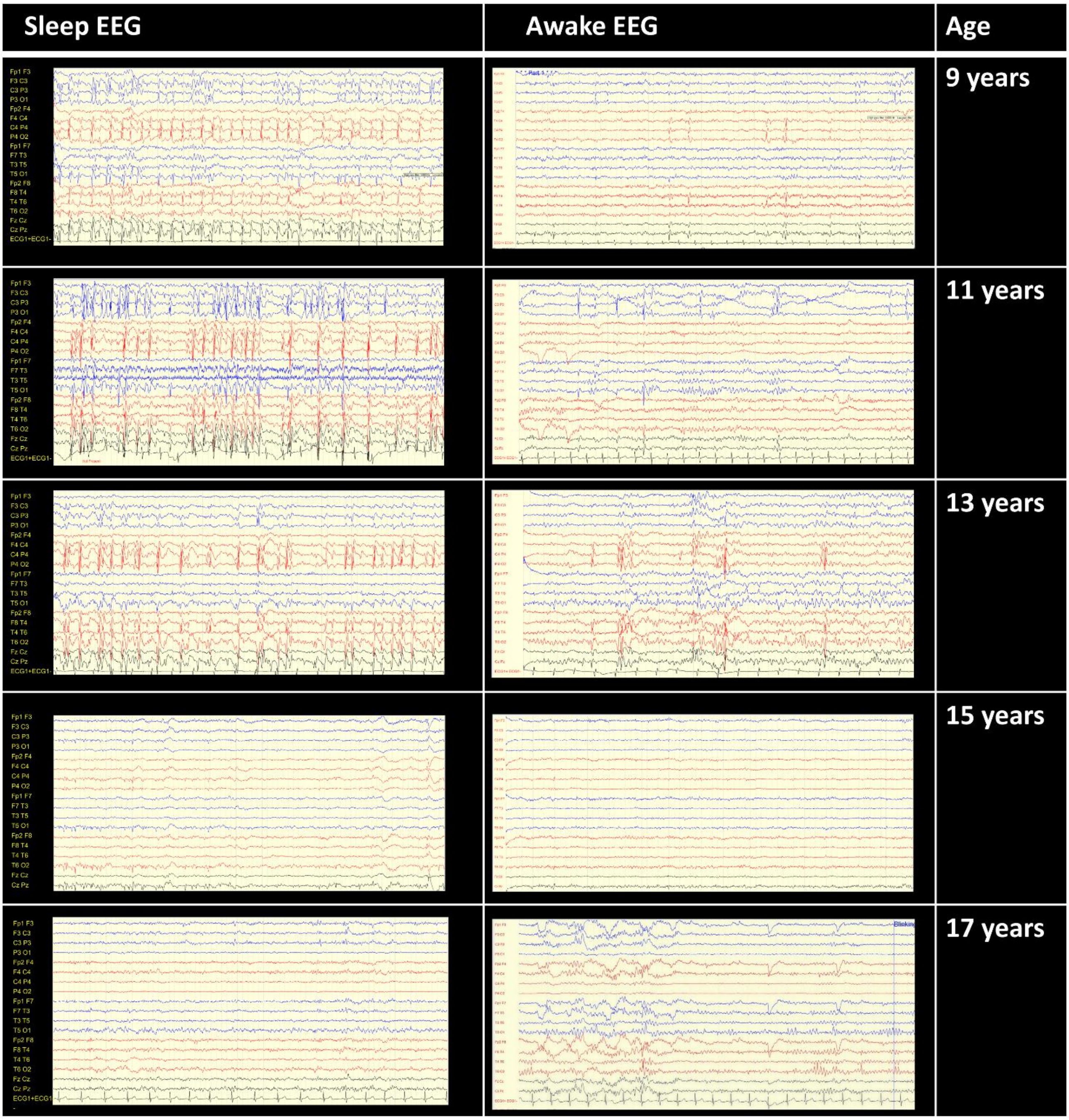


**Supplementary Figure 1. The sleep- and age-dependent changes in patients EEG recordings obtained at different ages using the international 10–20 electrode placement system.** The antiseizure medications (ASMs) administered at the time of each recording are also indicated. At 9 years of age (ASMs: sulthiame, ethosuximide, clobazam), 11 years (ASMs: sulthiame, ethosuximide, cannabis oil), and 13 years (ASMs: sulthiame, cannabis oil), EEGs consistently demonstrated frequent independent right and left centrotemporal epileptiform sharp-wave discharges that increased markedly during sleep, occupying approximately 50–80% of the recording, with a substantial reduction during wakefulness. At 15 years of age (ASMs: sulthiame, cannabis oil), there was a marked reduction in the frequency and amplitude of centrotemporal epileptiform activity during both sleep and wakefulness. By 17 years of age (ASMs: sulthiame, cannabis oil), centrotemporal epileptiform activity had decreased further, with only intermittent left centrotemporal sharp waves during sleep and near-complete resolution of epileptiform activity during wakefulness.

#### *Supplementary Figure 2*

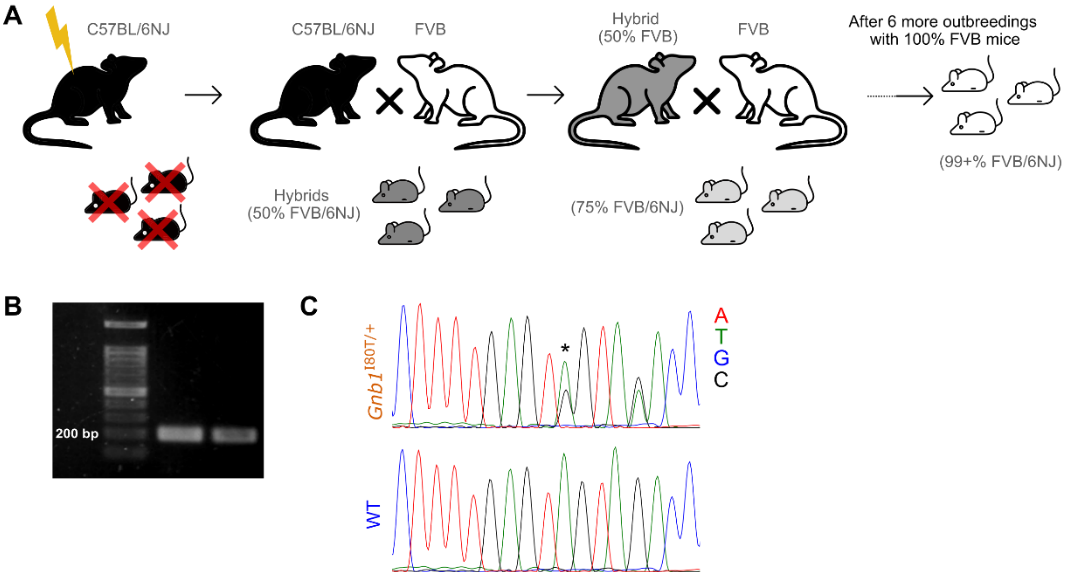


**Supplementary Figure 2. Breeding scheme and the genotyping method of *Gnb1*$^{I80T}$ mice.** (**A**) Schematic of the generational cross-breeding scheme of C57BL/6NJ *Gnb1*$^{I80T/+}$ mice with WT FVB/NJ mice to achieve >99% FVB/NJ *Gnb1*$^{I80T/+}$ congenic mice. The images of mice were downloaded from freely available internet sites: https://www.nashpestmanagement.com/ https://www.commonwealthext.com/ (large mouse) and https://www.flaticon.com/free-icon/mouse_2683099?related_id=2683103&origin=search (small mouse). (**B**) Representative DNA agarose gel showing the 200 bp PCR product obtained from tail-derived genomic DNA, used for routine genotyping of *Gnb1*$^{I80T/+}$ and WT littermates. (**C**) Sanger sequencing chromatograms comparing *Gnb1*$^{I80T/+}$ (top) and WT (bottom) mice at the *Gnb1* locus (Chr4, GRCm38), confirming the heterozygous point mutation c.239T>C (p.Ile80Thr) indicated by *. Overlapping peaks at the mutation site in the *Gnb1*$^{I80T/+}$ trace are indicative of heterozygosity. Note the additional c.243C>T mutation four base pairs 3' to the desired c.239T>C mutation. This is a silent guide-blocking mutation (ATC > ATT) that does not alter the protein sequence.

#### *Supplementary Figure 3*

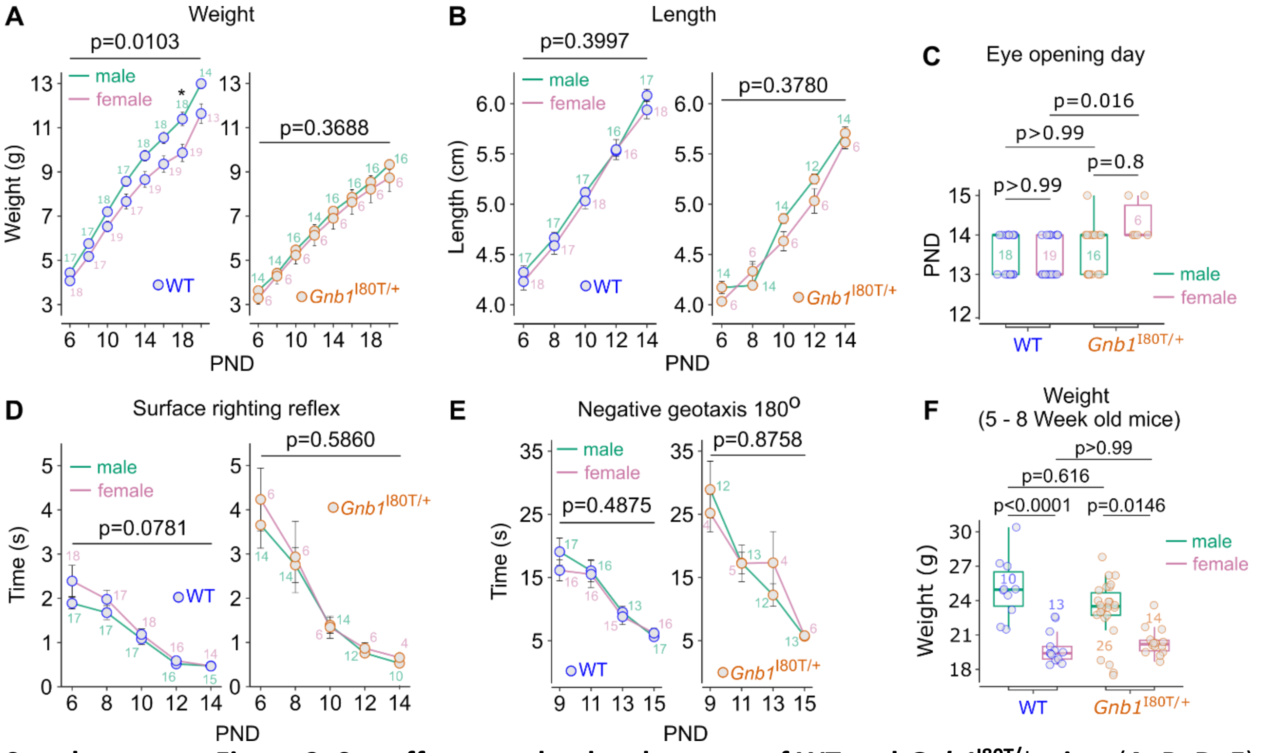


**Supplementary Figure 3. Sex effects on the development of WT and *Gnb1*$^{I80T/+}$ mice.** (**A**, **B**, **D**, **E**) Assessment of developmental milestones from P6 to P20: (**A**) body weight, (**B**) body length, (**D**) surface righting reflex, and (**E**) negative geotaxis (latency to reorient 180° upward from a downward-facing starting position). Line colors distinguish sex; error bars represent mean ± SEM; sample sizes are indicated beside each data point. (**C**) Eye-opening day. (**F**) Body weight comparison of mice at 5–8 weeks (P36–P57) of age. (**C**, **F**) Boxes represent median and interquartile range; whiskers extend to 1.5× IQR; box outline colors distinguish sex; individual data points represent single animals colored by genotype. Statistics: (**A**, **B**, **D**, **E**) mixed-effects analysis with Šídák's multiple comparisons test; (**C**, **F**) Kruskal-Wallis test with Dunn's multiple comparisons test.

*Supplementary Figure 4*

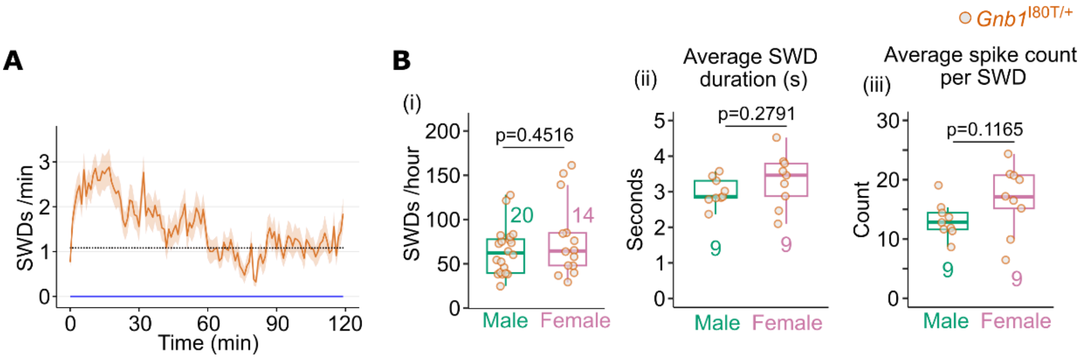


**Supplementary figure 4: Analysis of spike-wave discharges over time and stratification by sex.** (**A**) Number of SWDs per minute over a 2-hour recording period. Black line, drawn by hand, shows the SWD level that was assigned as stabilized baseline. (A linear fit line drawn between 60 and 120 s was nearly horizontal with a slope of 0.0071; not shown). WT (blue line): *n* = 4–6 males, 4–6 females; *Gnb1*$^{I80T}$ (dark orange line): *n* = 20 males, 14 females. (**B**) Analysis of sex effects on the spike-wave discharges in *Gnb1*$^{I80T}$ mice: (i) Total number of SWDs recorded during one hour following a 30-minute habituation period – Mann-Whitney test; (ii) average SWD duration – Welch's t-test; (iii) Average number of spikes per SWD – Welch's t-test. Boxes represent median and interquartile range; whiskers extend to 1.5× IQR; box outlines distinguish sex. Individual data points represent single animals colored by genotype, sample sizes are indicated below each box.

***Supplementary Figure 5***

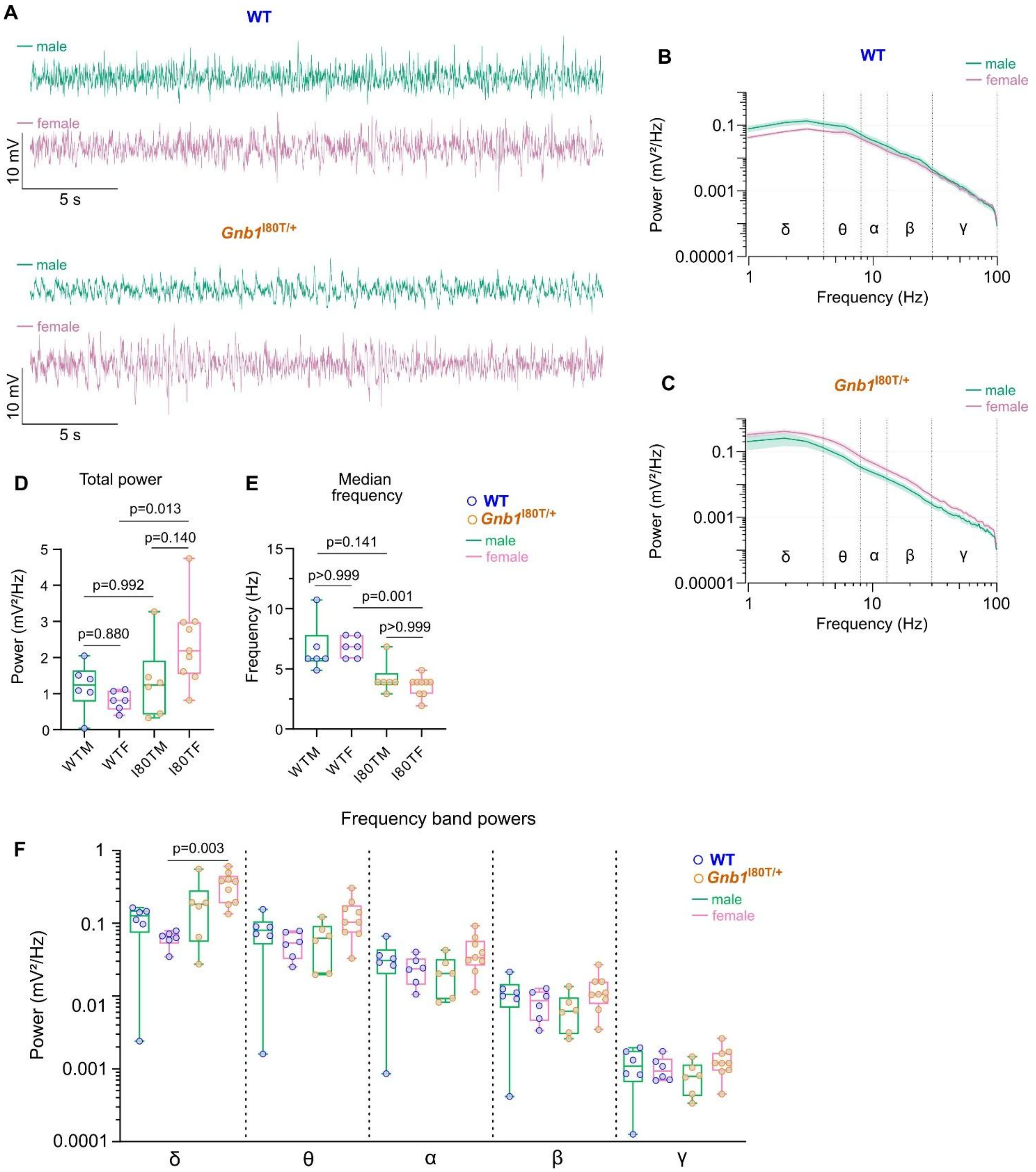


**Supplementary figure 5: Analysis of sex effects on background ECoG activity**. (**A**) Example ECoG traces of WT and *Gnb1*$^{I80T}$ male and female mice. (**B-F**) Analysis of sex effects within WT and *Gnb1*$^{I80T}$ genotypes: (**B, C**) power spectral density; (**D**) total power; (**E**) median frequency; (**F**) Power within frequency bands. Boxes represent interquartile range with median represented as a line within the box; whiskers from minimum to maximum value (two-way RM ANOVA: p=0.007; significance mentioned only for comparisons with p<0.05). Data points represent single animals. Statistics: (**D**) Welch's t-test; (**E**) Mann-Whitney test; (**F**) two-way ANOVA with Šídák's multiple comparisons test. WT: *n* = 6 males, 6 females; *Gnb1*$^{I80T}$: *n* = 6 males, 9 females.

#### *Supplementary Figure 6*

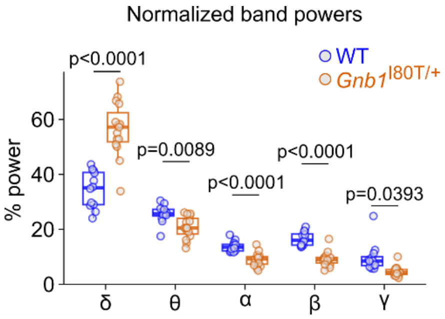


**Supplementary figure 6: Normalized frequency band powers in *Gnb1*$^{I80T}$ and WT mice.** Frequency band powers were normalized to total power and presented as % of total. Analysis: two-way ANOVA with Šídák's multiple comparisons test. Frequency bands: δ-0.9–3.9 Hz; θ-4.8–7.8 Hz; α-8.7–12.7 Hz; β-13.6–29.3 Hz; γ-30.2–99.6 Hz. WT: *n* = 6 males, 6 females; *Gnb1*$^{I80T}$: *n* = 6 males, 9 females

#### *Supplementary Figure 7*

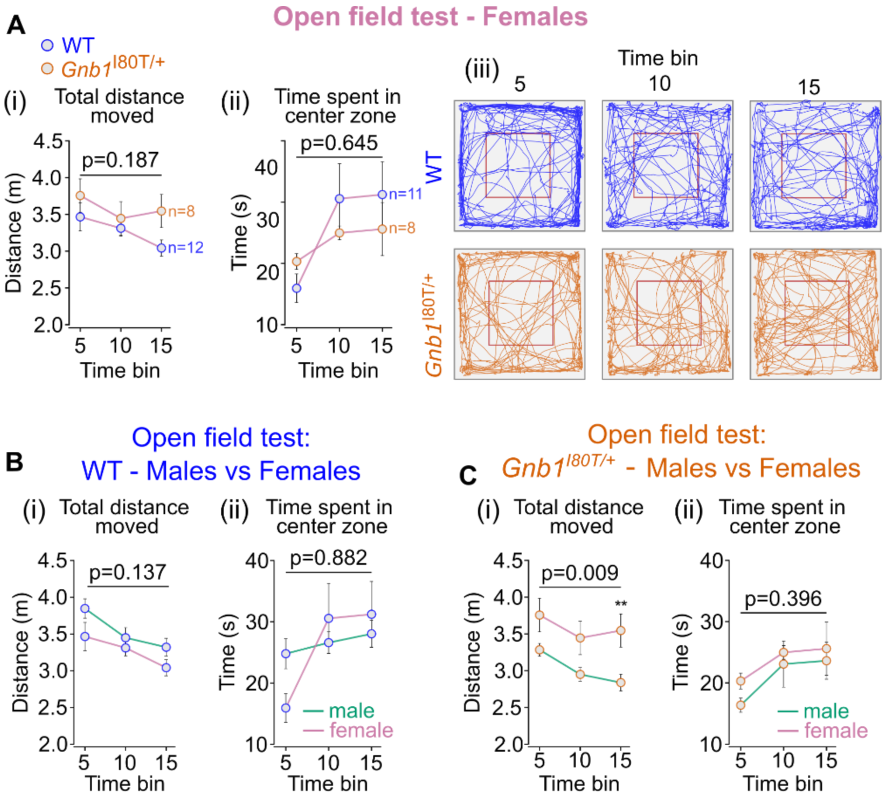


**Supplementary Figure 7. Behavioral assessment of *Gnb1*$^{I80T/+}$ females in open field and comparison of males versus females within genotype.** (**A**) Open field test conducted over a 15-minute period: (i) total distance moved, (ii) time spent in the center zone, and (iii) representative trace of WT and *Gnb1*$^{I80T/+}$ mice locomotion with center zone highlighted in red. Error bars represent mean ± SEM; sample sizes are indicated beside each line. **(B, C)** Analysis of sex effects in open field test. (**B**) Comparison of WT males and females; (**C**) Comparison of *Gnb1*$^{I80T/+}$ males and females. In B and C: (i) total distance moved and (ii) time spent in the center zone. WT: $n$ = 16 males, 13 females; *Gnb1*$^{I80T/+}$: $n$ = 13 males, 8 females. Error bars represent mean ± SEM; line colors distinguish sex. Statistics (A, B, C): two-way ANOVA with Šídák's multiple comparisons test. $^{**}p < 0.01$ for C(i).

*Supplementary Figure 8*

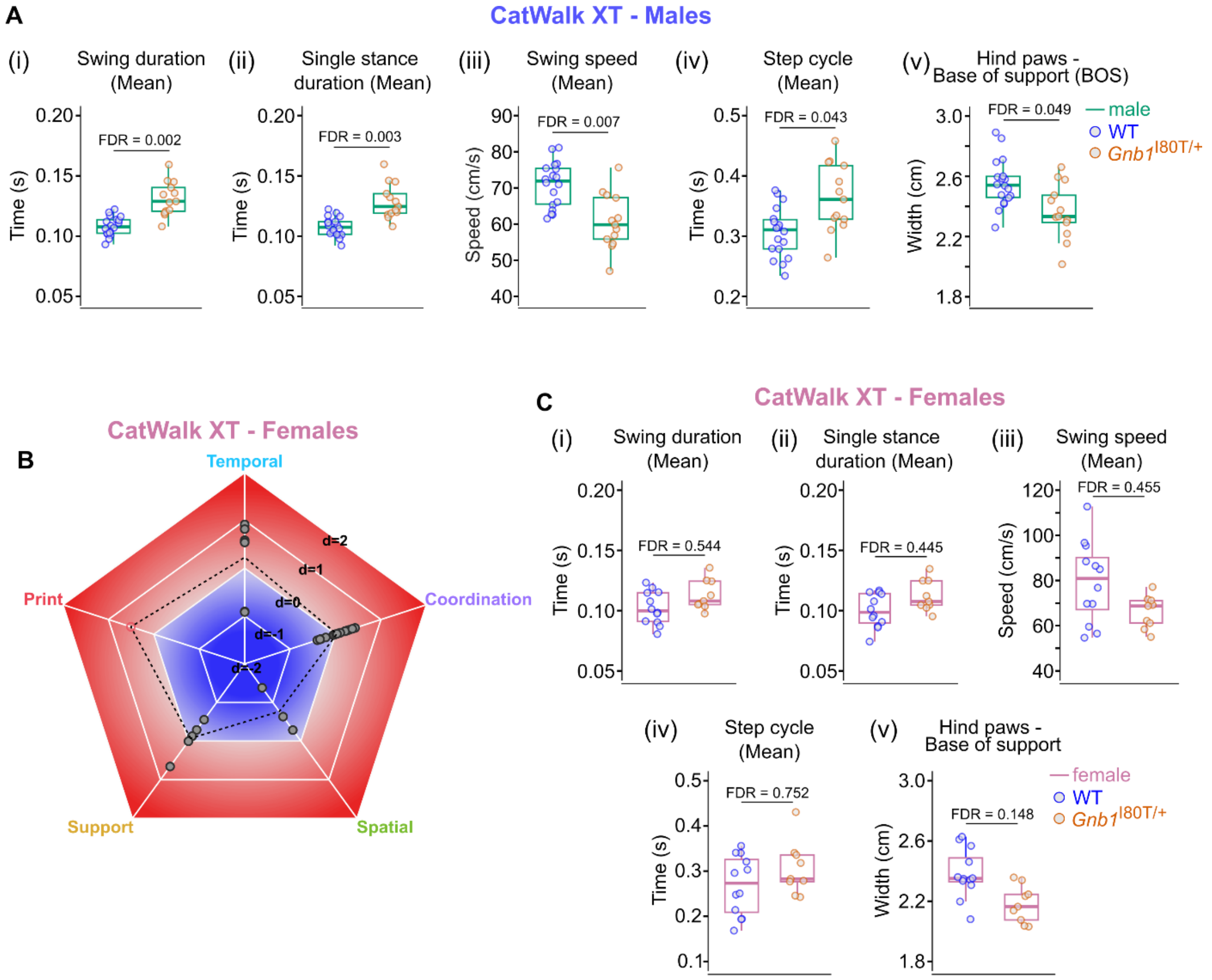


**Supplementary Figure 8. CatWalk XT test: Details of significantly altered gait parameters in males and parallel analysis in females.** (**A**) The five CatWalk XT parameters found to be significantly affected in males (Fig. 4B): (i) swing duration (mean airborne time of paws during a step cycle), (ii) single stance duration (mean time a paw supported weight without diagonal limb ground contact), (iii) swing speed (mean travel speed of paws during airborne phase), (iv) step cycle duration (mean total duration of one complete gait cycle), and (v) hind paw base of support (mean lateral distance between left and right hind paw placements). Parameters in (i–iv) were averaged across all four paws (see Supplementary Table 1a for full list of parameters). (**B**) CatWalk XT gait analysis for female mice depicted as a pentagon spider plot of Cohen's d values for all 36 parameters, categorized into five domains (Temporal, Spatial, Coordination, Support, and Print); compare with Fig. 4B for males. Each point represents one parameter; points are colored by domain for FDR-significant parameters (FDR < 0.05) and shown in grey for non-significant parameters; the dashed polygon connects the category mean Cohen's d per domain. (**C**) The five CatWalk XT parameters found to be significantly affected in males (Fig. 3B), plotted for females for comparison. See Supplementary Table 1b for full parameter list. In A – C: WT: *n* = 12 females, 18 males; *Gnb1*$^{I80T/+}$: *n* = 9 females, 13 males (sample sizes may vary per parameter; see Supplementary Table 1); boxes represent median and interquartile range; whiskers extend to 1.5× IQR; box outline colors distinguish sex; individual data points represent single animals colored by genotype. Statistics: (A, B, C) Welch's t-test or Mann-Whitney U test with Benjamini-Hochberg FDR correction.

#### *Supplementary Figure 9*

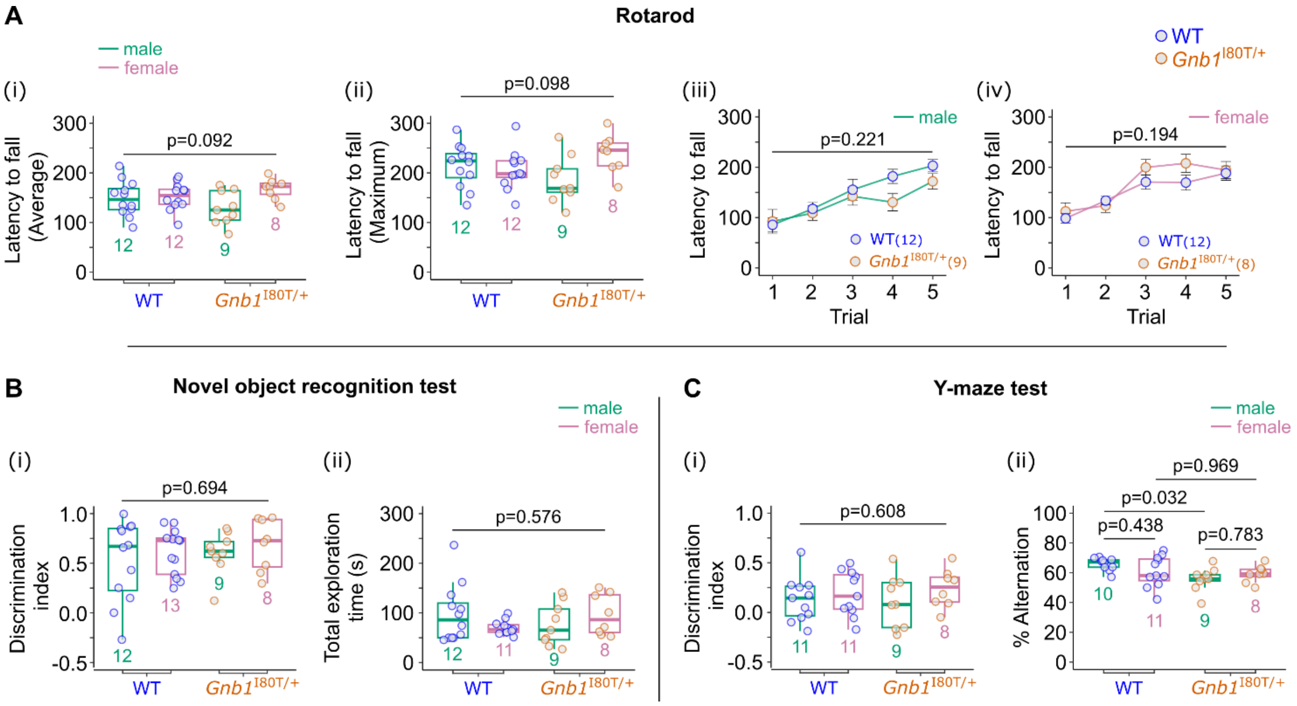


**Supplementary Figure 9. Summary of additional behavioral tests: comparison of WT and *Gnb1*$^{I80T/+}$, with sex stratification.** (**A**) Rotarod test (i, ii) average latency to fall and maximum latency to fall, (iii, iv) latency to fall per trial comparison of genotypes within male and female sex. (**B**) Novel object recognition test: (i) discrimination index (DI = time spent in close proximity to novel object / (time spent exploring novel object + time spent exploring familiar object)) and (ii) total exploration time (time spent exploring novel object + time spent exploring familiar object). (**C**) Y-maze test: (i) discrimination index (DI = time spent in novel arm / (time spent in novel arm + time spent in familiar arm)) and (ii) percentage alternation (percentage of successive entries into all three arms without repetition, out of total possible alternations). Throughout (**A** i, ii), (**B**), and (**C**): boxes represent median and interquartile range; whiskers extend to 1.5× IQR; box outline colors distinguish sex; individual data points represent single animals colored by genotype; sample sizes are indicated beside each box. For (**A** iii, iv): line colors distinguish sex; error bars represent mean ± SEM; sample sizes are indicated on labels. Statistics: (A i, ii) one-way ANOVA; (A iii, iv) two-way repeated measures ANOVA; (B i) one-way ANOVA; (B ii) Kruskal-Wallis test; (C i) one-way ANOVA; (C ii) one-way ANOVA with Tukey's multiple comparisons test.

*Supplementary Figure 10*

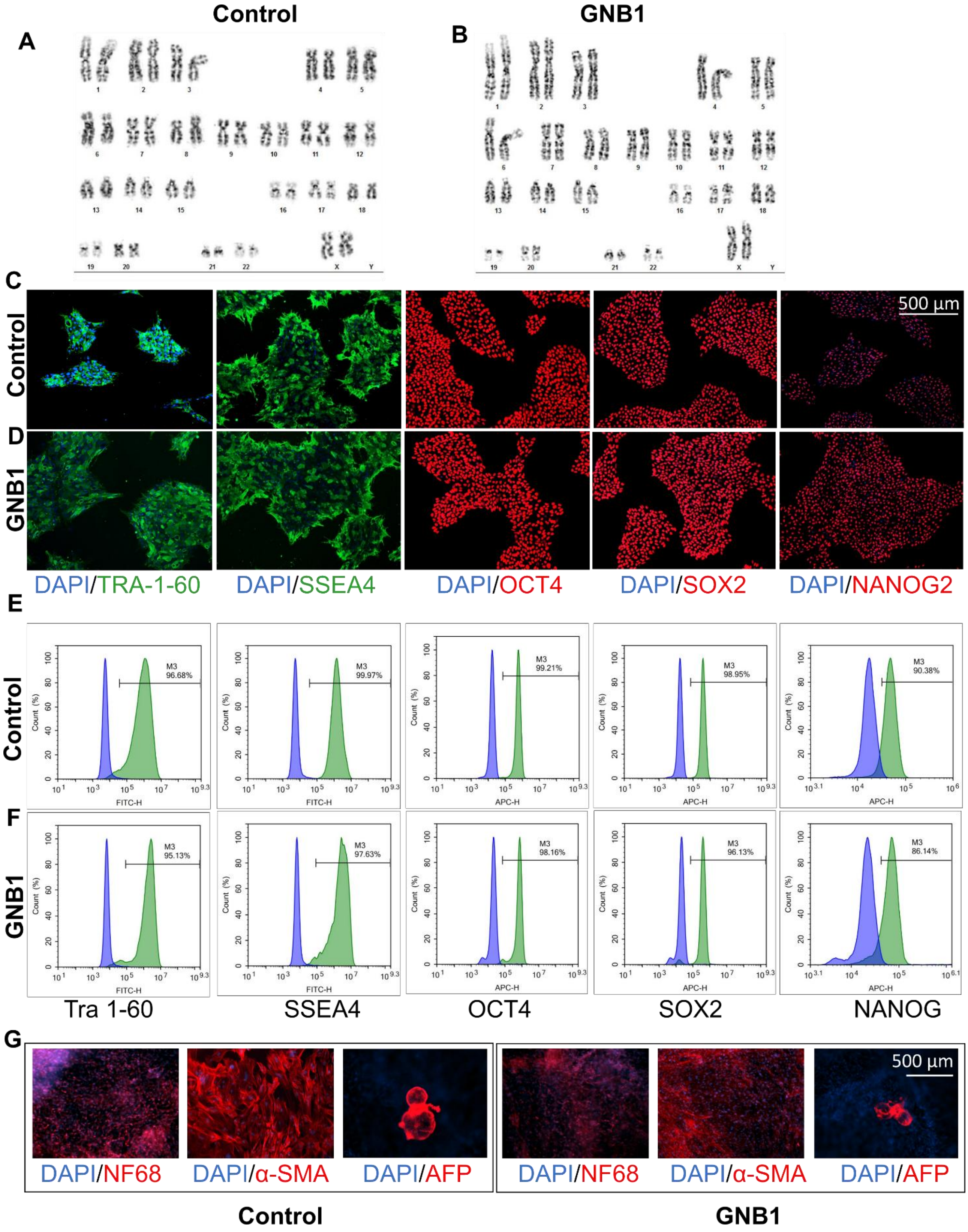


**Supplementary Figure 10. Generation and characterization of patient-derived induced pluripotent stem cell (iPSC) lines carrying the GNB1$^{I80T/+}$ variant. (A, B)** Representative G-banding karyotypes in control and patient-derived iPSC lines. Primary fibroblasts were reprogrammed by electroporation of non-integrating episomal plasmids encoding OCT4, SOX2, KLF4, L-MYC, shP53, LIN28, and SV40LT, as previously described (*1*). **(C,D)** Immunocytochemical characterization of pluripotency markers in generated iPSC lines, showing expression of TRA-1-60, SSEA-4, OCT4,

SOX2, and NANOG. Nuclei were counterstained with DAPI (blue). **(E, F)** Quantification of pluripotency marker expression by fluorescence-activated cell sorting (FACS). Histograms show expression of TRA-1-60, SSEA-4, OCT4, SOX2, and NANOG. Marker-positive populations ranged from 71–99%. **(G)** Immunocytochemistry for expression of lineage-specific markers: neurofilament light chain (NF-L; ectoderm), α-smooth muscle actin (α-SMA; mesoderm), and α-fetoprotein (AFP; endoderm). Nuclei were counterstained with DAPI (blue). Scale bar, 500 µm.

***Supplementary Figure 11***

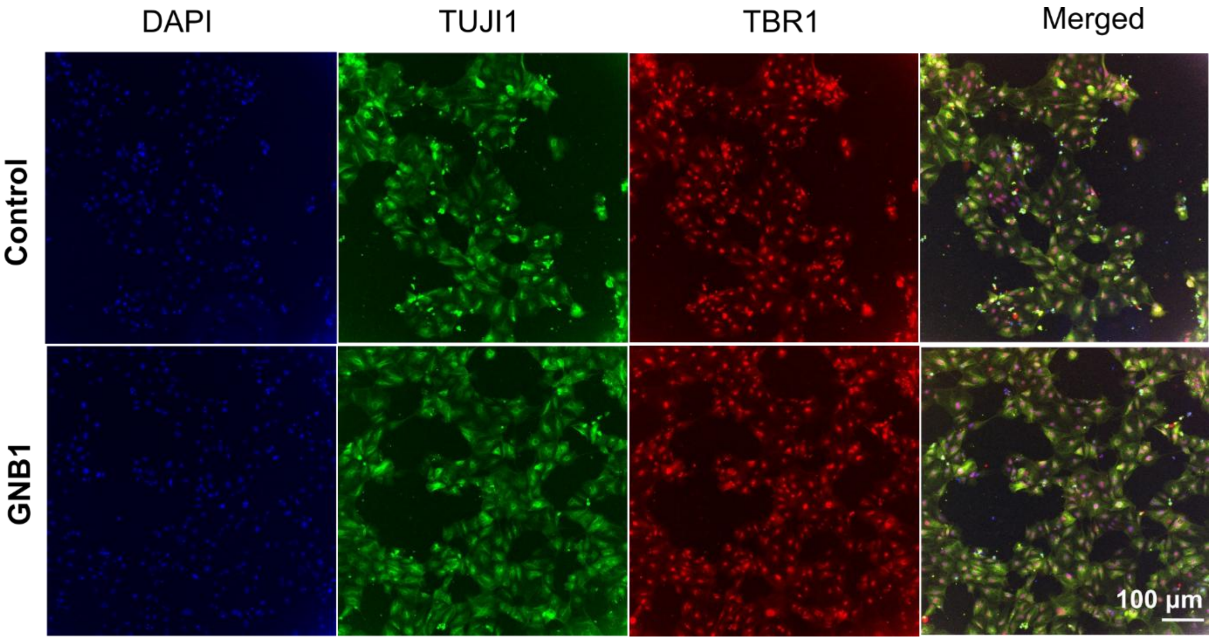


**Supplementary Figure 11. Validation of cortical neuronal differentiation from control and GNB1$^{I80T}$-derived iPSCs.** Representative immunofluorescence images of cortical neuron cultures differentiated from control and GNB1 patient-derived iPSC lines at day 13 of differentiation. Cells were stained for TUJ1 (green) to identify neuronal cells and TBR1 (red) to confirm cortical neuronal identity. Nuclei were counterstained with DAPI (blue). Merged images demonstrate co-expression of neuronal and cortical markers in both conditions, confirming successful differentiation into cortical neurons. Scale bar, 100 µm.

*Supplementary Figure 12*

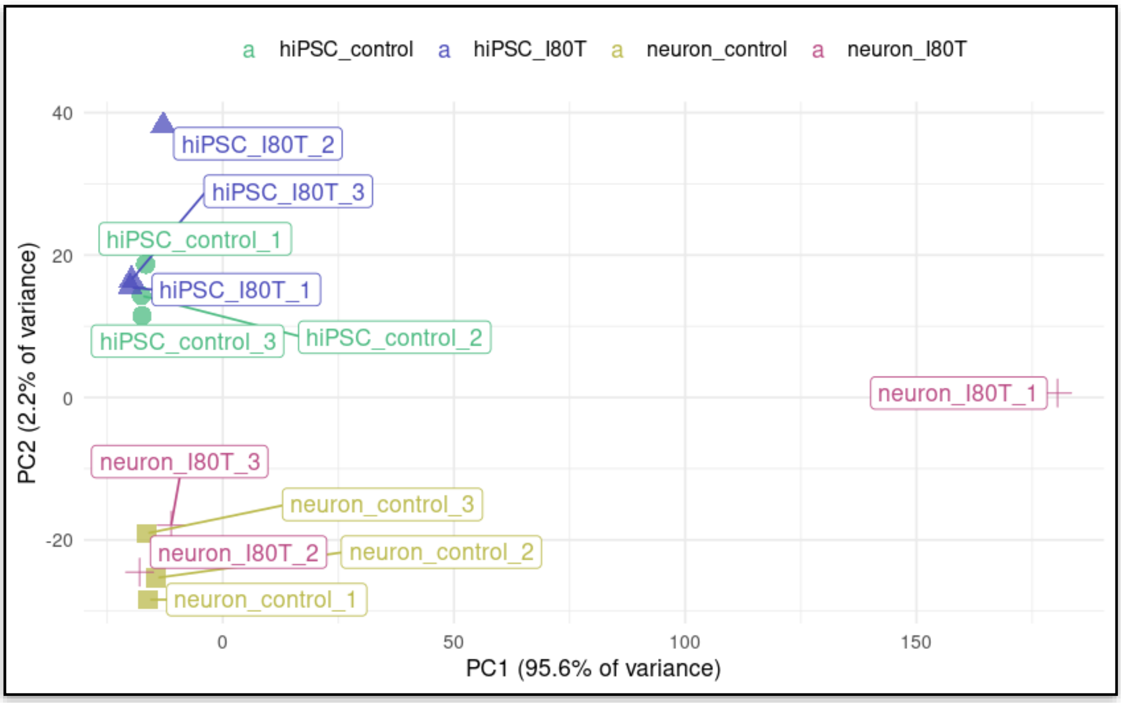


**Supplementary Figure 12. Principal component analysis (PCA) of hiPSCs and hiPSC-derived neurons.** PCA was performed on hiPSC samples (hiPSC_control and hiPSC_I80T) and hiPSC-derived neuron samples (neuron_control and neuron_I80T). PC1 explains 95.6% of the variance and PC2 explains 2.2% of the variance. The analysis reveals one outlier within the hiPSC-derived neuron group.

### Supplementary Tables

#### *Supplementary Table 1a*

Supplementary Table 1a. The Catwalk parameters analyzed (males).

| Parameter | P | Test | WT Mean | WT SD | I80T Mean | I80T SD | WT outliers | I80T outliers | FDR |
|---|---|---|---|---|---|---|---|---|---|
| **N: WT = 18; Gnb1-I80T/+ = 13** | | | | | | | | | |
| Avg_Swing_(s)_Mean | 5.6E-05 | Welch | 0.11 | 0.01 | 0.13 | 0.01 | 0 | 0 | 0.002 |
| Avg_SingleStance_(s)_Mean | 1.9E-04 | Welch | 0.11 | 0.01 | 0.13 | 0.01 | 0 | 0 | 0.003 |
| Avg_SwingSpeed_(cm/s)_Mean | 0.001 | Welch | 70.98 | 6.19 | 60.70 | 7.64 | 0 | 0 | 0.007 |
| Avg_StepCycle_(s)_Mean | 0.005 | Welch | 0.31 | 0.04 | 0.36 | 0.06 | 0 | 0 | 0.043 |
| BOS_HindPaws_Mean_(cm) | 0.007 | Welch | 2.55 | 0.16 | 2.37 | 0.18 | 0 | 0 | 0.049 |
| Run_Average_Speed_(cm/s)_Mean | 0.020 | Welch | 23.81 | 3.52 | 20.09 | 4.44 | 0 | 0 | 0.119 |
| Avg_Stand_(s)_Mean | 0.023 | Welch | 0.20 | 0.03 | 0.23 | 0.04 | 0 | 0 | 0.119 |
| StepSequence_RegularityIndex_(%) | 0.265 | MW | 98.09 | 1.46 | 96.63 | 3.37 | 0 | 0 | 0.734 |
| Couplings_LF->RH_CStat_Mean | 0.291 | Welch | 8.20 | 3.16 | 9.25 | 2.28 | 0 | 0 | 0.734 |
| Couplings_LH->RH_CStat_Mean | 0.185 | Welch | 49.08 | 2.38 | 50.31 | 2.54 | 0 | 0 | 0.734 |
| Couplings_LF->RF_CStat_Mean | 0.284 | Welch | 50.12 | 2.14 | 51.09 | 2.60 | 0 | 0 | 0.734 |
| Couplings_RH->LH_CStat_Mean | 0.227 | Welch | 49.88 | 2.35 | 48.70 | 2.81 | 0 | 0 | 0.734 |
| PhaseDispersions_LF->RH_CStat_Mean | 0.335 | Welch | 8.21 | 3.15 | 9.22 | 2.59 | 0 | 0 | 0.734 |
| PhaseDispersions_LH->RH_CStat_Mean | 0.229 | Welch | 49.65 | 2.33 | 50.76 | 2.57 | 0 | 0 | 0.734 |
| Support_Single_(%) | 0.192 | MW | 1.07 | 1.36 | 0.26 | 0.41 | 0 | 2 | 0.734 |
| Support_Lateral_(%) | 0.347 | MW | 2.42 | 1.36 | 2.83 | 1.30 | 1 | 2 | 0.734 |
| Avg_DutyCycle_(%)_Mean | 0.335 | Welch | 62.98 | 2.73 | 62.11 | 2.16 | 0 | 0 | 0.734 |
| Couplings_RF->LH_CStat_Mean | 0.386 | Welch | 8.69 | 3.53 | 7.51 | 3.51 | 1 | 1 | 0.738 |
| Couplings_RF->LF_CStat_Mean | 0.390 | Welch | 49.49 | 2.59 | 48.74 | 2.21 | 0 | 0 | 0.738 |
| PhaseDispersions_LF->RF_CStat_Mean | 0.449 | Welch | 50.51 | 2.20 | 51.15 | 2.40 | 0 | 0 | 0.809 |
| PhaseDispersions_RF->LH_CStat_Mean | 0.482 | Welch | 8.57 | 3.70 | 7.63 | 3.45 | 0 | 1 | 0.827 |
| PhaseDispersions_LF->LH_CStat_Mean | 0.519 | Welch | 58.84 | 2.68 | 58.04 | 3.76 | 0 | 0 | 0.849 |
| Couplings_RH->RF_CStat_Mean | 0.556 | Welch | 41.16 | 3.13 | 41.76 | 2.51 | 0 | 0 | 0.871 |
| Avg_StrideLength_(cm)_Mean | 0.621 | Welch | 7.18 | 0.45 | 7.08 | 0.58 | 0 | 0 | 0.894 |
| Couplings_LF->LH_CStat_Mean | 0.612 | Welch | 59.03 | 2.92 | 58.36 | 3.92 | 0 | 0 | 0.894 |
| Couplings_RF->RH_CStat_Mean | 0.984 | Welch | 58.13 | 3.25 | 58.11 | 3.22 | 0 | 0 | 0.984 |
| Couplings_LH->RF_CStat_Mean | 0.931 | Welch | 91.44 | 3.87 | 91.55 | 3.14 | 0 | 1 | 0.984 |
| Couplings_RH->LF_CStat_Mean | 0.967 | Welch | 91.09 | 2.60 | 91.06 | 2.22 | 1 | 0 | 0.984 |
| Couplings_LH->LF_CStat_Mean | 0.740 | Welch | 40.27 | 2.89 | 40.71 | 4.00 | 0 | 0 | 0.984 |
| PhaseDispersions_RF->RH_CStat_Mean | 0.841 | Welch | 58.07 | 3.17 | 57.83 | 3.12 | 0 | 0 | 0.984 |
| BOS_FrontPaws_Mean_(cm) | 0.922 | Welch | 1.63 | 0.07 | 1.63 | 0.06 | 0 | 1 | 0.984 |
| Support_Diagonal_(%) | 0.909 | Welch | 43.43 | 7.41 | 43.12 | 7.26 | 0 | 0 | 0.984 |
| Support_Three_(%) | 0.847 | Welch | 47.35 | 4.83 | 46.90 | 7.14 | 2 | 0 | 0.984 |
| Support_Girdle_(%) | 0.830 | MW | 0.56 | 0.84 | 0.45 | 0.64 | 1 | 1 | 0.984 |
| Support_Four_(%) | 0.904 | MW | 4.73 | 4.03 | 4.16 | 2.86 | 0 | 0 | 0.984 |
| Avg_PrintArea_(cm²)_Mean | 0.838 | Welch | 0.31 | 0.05 | 0.31 | 0.04 | 0 | 0 | 0.984 |

#### *Supplementary Table 1b*

Supplementary Table 1b. The Catwalk parameters analyzed (females).

| Parameter | P | Test | WT Mean | WT SD | I80T Mean | I80T SD | WT outliers | I80T outliers | FDR |
|---|---|---|---|---|---|---|---|---|---|
| **N: WT = 12; Gnb1-I80T/+ = 9** | | | | | | | | | |
| BOS_HindPaws_Mean_(cm) | 0.004 | Welch | 2.38 | 0.16 | 2.18 | 0.12 | 0 | 0 | 0.148 |
| Run_Average_Speed_(cm/s)_Mean | 0.038 | Welch | 30.92 | 10.36 | 23.18 | 5.12 | 0 | 0 | 0.445 |
| Avg_SwingSpeed_(cm/s)_Mean | 0.035 | Welch | 79.30 | 17.97 | 66.06 | 7.22 | 0 | 0 | 0.445 |
| Avg_SingleStance_(s)_Mean | 0.049 | Welch | 0.10 | 0.01 | 0.11 | 0.01 | 0 | 0 | 0.445 |
| Avg_Swing_(s)_Mean | 0.076 | Welch | 0.10 | 0.01 | 0.11 | 0.01 | 0 | 0 | 0.544 |
| Avg_StrideLength_(cm)_Mean | 0.132 | Welch | 7.31 | 1.02 | 6.79 | 0.45 | 0 | 0 | 0.699 |
| Support_Three_(%) | 0.136 | Welch | 35.97 | 14.01 | 44.27 | 10.41 | 0 | 0 | 0.699 |
| Support_Diagonal_(%) | 0.209 | Welch | 52.62 | 14.73 | 45.13 | 11.66 | 0 | 0 | 0.752 |
| Avg_StepCycle_(s)_Mean | 0.192 | Welch | 0.27 | 0.07 | 0.31 | 0.06 | 0 | 0 | 0.752 |
| Avg_Stand_(s)_Mean | 0.23 | Welch | 0.17 | 0.05 | 0.20 | 0.05 | 0 | 0 | 0.752 |
| Avg_DutyCycle_(%)_Mean | 0.169 | MW | 58.65 | 5.02 | 61.07 | 3.55 | 0 | 0 | 0.752 |
| StepSequence_RegularityIndex_(%) | 0.311 | Welch | 96.25 | 3.18 | 97.43 | 1.97 | 0 | 0 | 0.860 |
| Avg_PrintArea_(cm²)_Mean | 0.291 | Welch | 0.23 | 0.04 | 0.26 | 0.06 | 0 | 0 | 0.860 |
| Couplings_LF->RH_CStat_Mean | 0.508 | MW | 7.22 | 2.99 | 7.95 | 2.63 | 0 | 0 | 0.923 |
| Couplings_RH->RF_CStat_Mean | 0.363 | Welch | 42.18 | 3.58 | 40.83 | 3.02 | 0 | 0 | 0.923 |
| Couplings_LH->LF_CStat_Mean | 0.451 | Welch | 41.65 | 4.15 | 40.51 | 2.61 | 0 | 0 | 0.923 |
| PhaseDispersions_LF->LH_CStat_Mean | 0.464 | Welch | 57.77 | 4.14 | 58.78 | 1.85 | 0 | 0 | 0.923 |
| PhaseDispersions_LF->RF_CStat_Mean | 0.508 | MW | 50.44 | 2.73 | 50.47 | 1.85 | 0 | 0 | 0.923 |
| BOS_FrontPaws_Mean_(cm) | 0.513 | Welch | 1.62 | 0.15 | 1.58 | 0.13 | 0 | 0 | 0.923 |
| Support_Single_(%) | 0.502 | Welch | 1.55 | 1.65 | 1.12 | 1.25 | 0 | 0 | 0.923 |
| Couplings_RF->RH_CStat_Mean | 0.59 | Welch | 57.53 | 3.67 | 58.27 | 2.46 | 0 | 0 | 0.951 |
| Couplings_LF->LH_CStat_Mean | 0.628 | Welch | 58.33 | 4.68 | 59.12 | 2.58 | 0 | 0 | 0.951 |
| Couplings_RH->LF_CStat_Mean | 0.634 | Welch | 91.64 | 3.29 | 90.92 | 3.43 | 0 | 0 | 0.951 |
| PhaseDispersions_RF->RH_CStat_Mean | 0.62 | Welch | 57.18 | 3.33 | 57.82 | 2.46 | 0 | 0 | 0.951 |
| Couplings_RF->LH_CStat_Mean | 1 | MW | 15.84 | 26.48 | 8.89 | 3.05 | 0 | 0 | 1.000 |
| Couplings_LH->RF_CStat_Mean | 0.972 | MW | 83.67 | 26.16 | 91.02 | 1.48 | 0 | 0 | 1.000 |
| Couplings_LH->RH_CStat_Mean | 0.905 | Welch | 49.56 | 3.33 | 49.43 | 1.63 | 0 | 0 | 1.000 |
| Couplings_LF->RF_CStat_Mean | 0.754 | MW | 50.08 | 2.66 | 49.99 | 2.46 | 0 | 0 | 1.000 |
| Couplings_RH->LH_CStat_Mean | 0.981 | Welch | 50.17 | 3.51 | 50.20 | 2.32 | 0 | 0 | 1.000 |
| Couplings_RF->LF_CStat_Mean | 0.866 | Welch | 49.24 | 3.07 | 49.45 | 2.49 | 0 | 0 | 1.000 |
| PhaseDispersions_RF->LH_CStat_Mean | 0.808 | MW | 16.04 | 26.21 | 9.20 | 2.36 | 0 | 0 | 1.000 |
| PhaseDispersions_LF->RH_CStat_Mean | 0.816 | Welch | 7.91 | 2.69 | 8.18 | 2.57 | 0 | 0 | 1.000 |
| PhaseDispersions_LH->RH_CStat_Mean | 0.942 | Welch | 49.96 | 2.90 | 50.03 | 1.83 | 0 | 0 | 1.000 |
| Support_Lateral_(%) | 0.781 | Welch | 5.43 | 2.80 | 5.10 | 2.57 | 0 | 0 | 1.000 |
| Support_Girdle_(%) | 0.937 | MW | 0.38 | 0.55 | 0.32 | 0.46 | 0 | 0 | 1.000 |
| Support_Four_(%) | 0.802 | MW | 4.04 | 4.54 | 4.07 | 4.82 | 0 | 0 | 1.000 |

#### *Supplementary Table 2.*

Supplementary Table 2. The 323 overlapping gene set (concordantly changed differentially expressed genes from *Gnb1*$^{I80T/+}$ mouse cortex and patient's hiPSC-derived neurons.

| Mouse cortex | | | | | hiPSC-derived neurons | | | | |
|---|---|---|---|---|---|---|---|---|---|
| **Genes** | **FDR.x** | **fold Change.x** | **log2Fold Change.x** | **P-Value** | **SYMBOL.y** | **FDR.y** | **fold Change.y** | **log2Fold Change.y** | **P-Value** |
| CASP8AP2 | 0.007664 | 1.62 | 0.70 | 0.00057 | CASP8AP2 | 1.637E-06 | 1.87E+07 | 24.16 | 9.331E-08 |
| ADAMTS4 | 0.000302 | 1.69 | 0.76 | 1.2E-05 | ADAMTS4 | 2.182E-24 | 73.62 | 6.20 | 1.714E-26 |
| RGS7BP | 0.004995 | 1.14 | 0.18 | 0.00033 | RGS7BP | 5.269E-18 | 44.18 | 5.47 | 7.152E-20 |
| PAQR9 | 0.011777 | 1.21 | 0.28 | 0.00098 | PAQR9 | 1.577E-14 | 41.88 | 5.39 | 2.846E-16 |
| MOBP | 2.3E-17 | 1.34 | 0.42 | 6.1E-20 | MOBP | 6.039E-08 | 40.02 | 5.32 | 2.645E-09 |
| DLGAP2 | 0.001945 | 1.34 | 0.42 | 0.00011 | DLGAP2 | 1.313E-05 | 24.13 | 4.59 | 9.147E-07 |
| GRM7 | 9E-05 | 1.39 | 0.48 | 2.8E-06 | GRM7 | 0.0016081 | 21.69 | 4.44 | 0.0002147 |
| COL19A1 | 0.002341 | 1.38 | 0.46 | 0.00013 | COL19A1 | 0.0269509 | 19.61 | 4.29 | 0.0065893 |
| ATP2B2 | 2.07E-14 | 1.34 | 0.42 | 7.2E-17 | ATP2B2 | 3.018E-05 | 17.72 | 4.15 | 2.298E-06 |
| HKDC1 | 0.011582 | 1.86 | 0.89 | 0.00096 | HKDC1 | 0.0033045 | 17.58 | 4.14 | 0.0005046 |
| SCN1A | 9.35E-19 | 1.45 | 0.54 | 1.9E-21 | SCN1A | 2.106E-10 | 16.61 | 4.05 | 6.105E-12 |
| KCNT1 | 0.000117 | 1.47 | 0.56 | 3.9E-06 | KCNT1 | 4.272E-10 | 15.54 | 3.96 | 1.266E-11 |
| SCN4B | 0.013844 | 1.23 | 0.30 | 0.0012 | SCN4B | 5.265E-09 | 15.16 | 3.92 | 1.895E-10 |
| CFAP74 | 0.004699 | 2.90 | 1.54 | 0.00031 | CFAP74 | 0.000242 | 12.34 | 3.62 | 2.362E-05 |
| AATK | 3.71E-08 | 1.32 | 0.40 | 4.3E-10 | AATK | 6.288E-09 | 11.63 | 3.54 | 2.276E-10 |
| CD93 | 0.021549 | 1.87 | 0.91 | 0.00213 | CD93 | 0.0407273 | 11.43 | 3.52 | 0.0112357 |
| TMEM72 | 1.1E-12 | 1.77 | 0.83 | 5.3E-15 | TMEM72 | 0.0295446 | 11.43 | 3.51 | 0.0073953 |
| TBC1D30 | 2.46E-10 | 1.41 | 0.50 | 1.8E-12 | TBC1D30 | 4.606E-14 | 11.02 | 3.46 | 8.795E-16 |
| CACNA1G | 0.000865 | 1.46 | 0.55 | 4.1E-05 | CACNA1G | 1.53E-10 | 10.93 | 3.45 | 4.329E-12 |
| ADARB2 | 0.012607 | 1.28 | 0.35 | 0.00107 | ADARB2 | 5.174E-07 | 10.14 | 3.34 | 2.647E-08 |
| KCNK9 | 4.22E-05 | 1.46 | 0.54 | 1.2E-06 | KCNK9 | 0.0007765 | 9.97 | 3.32 | 9.177E-05 |
| MAG | 9.11E-10 | 1.49 | 0.58 | 7.2E-12 | MAG | 0.003826 | 9.62 | 3.27 | 0.0006024 |
| MKX | 0.002701 | 1.95 | 0.96 | 0.00016 | MKX | 3.235E-09 | 9.32 | 3.22 | 1.128E-10 |
| GPR158 | 0.042674 | 1.12 | 0.16 | 0.00531 | GPR158 | 0.0003268 | 8.63 | 3.11 | 3.349E-05 |
| RGS11 | 0.005837 | 1.80 | 0.85 | 0.0004 | RGS11 | 8.485E-05 | 7.95 | 2.99 | 7.273E-06 |
| GABRB2 | 0.01391 | 1.18 | 0.24 | 0.00121 | GABRB2 | 0.0004768 | 6.70 | 2.74 | 5.195E-05 |
| CARNS1 | 0.039197 | 2.20 | 1.13 | 0.00472 | CARNS1 | 0.0058139 | 6.61 | 2.73 | 0.0009953 |
| GPR26 | 5.74E-08 | 1.35 | 0.43 | 7E-10 | GPR26 | 7.699E-05 | 6.14 | 2.62 | 6.516E-06 |
| PLXNB3 | 0.00656 | 1.58 | 0.66 | 0.00047 | PLXNB3 | 0.0084989 | 5.66 | 2.50 | 0.0015776 |
| PCDHA7 | 0.02159 | 2.09 | 1.06 | 0.00214 | PCDHA7 | 4.209E-05 | 5.65 | 2.50 | 3.336E-06 |
| CABP1 | 0.001475 | 1.29 | 0.37 | 7.7E-05 | CABP1 | 0.0002018 | 5.46 | 2.45 | 1.93E-05 |
| UNKL | 0.040761 | 1.38 | 0.46 | 0.00498 | UNKL | 4.869E-19 | 5.33 | 2.41 | 6.029E-21 |
| KCNJ9 | 0.000319 | 1.29 | 0.37 | 1.3E-05 | KCNJ9 | 0.0197164 | 5.25 | 2.39 | 0.0044509 |
| ARC | 1.05E-29 | 1.55 | 0.63 | 7.9E-33 | ARC | 0.0022657 | 5.24 | 2.39 | 0.0003214 |
| TRPC3 | 0.006855 | 1.97 | 0.98 | 0.00049 | TRPC3 | 0.0072476 | 4.88 | 2.29 | 0.0012974 |
| SCN2A | 1.72E-05 | 1.17 | 0.22 | 4.1E-07 | SCN2A | 0.0001756 | 4.75 | 2.25 | 1.651E-05 |
| BRD2 | 0.002128 | 1.29 | 0.36 | 0.00012 | BRD2 | 1.608E-31 | 4.63 | 2.21 | 8.555E-34 |
| SPTBN4 | 0.02302 | 1.24 | 0.31 | 0.00231 | SPTBN4 | 1.573E-05 | 4.56 | 2.19 | 1.119E-06 |
| CX3CL1 | 0.027069 | 1.13 | 0.18 | 0.00286 | CX3CL1 | 0.0043849 | 4.55 | 2.19 | 0.0007107 |
| PDE4A | 0.009564 | 1.34 | 0.42 | 0.00075 | PDE4A | 0.0076589 | 4.54 | 2.18 | 0.0013889 |
| TLE2 | 0.010677 | 2.47 | 1.31 | 0.00086 | TLE2 | 1.153E-06 | 4.48 | 2.16 | 6.356E-08 |
| SPRED3 | 0.024673 | 1.41 | 0.50 | 0.00252 | SPRED3 | 0.0110373 | 4.43 | 2.15 | 0.0021728 |
| ABCA2 | 0.001121 | 1.28 | 0.36 | 5.5E-05 | ABCA2 | 0.0002907 | 4.20 | 2.07 | 2.919E-05 |

***Supplementary Table 2 continued***

| | | | | | | | | | |
|---|---|---|---|---|---|---|---|---|---|
| GRIN2A | 1.61E-07 | 1.24 | 0.31 | 2.1E-09 | GRIN2A | 1.266E-05 | 3.94 | 1.98 | 8.777E-07 |
| HECW1 | 0.014024 | 1.25 | 0.32 | 0.00122 | HECW1 | 0.0125445 | 3.90 | 1.96 | 0.0025456 |
| NOS1AP | 0.010214 | 1.38 | 0.47 | 0.00082 | NOS1AP | 0.0091316 | 3.83 | 1.94 | 0.0017236 |
| SLC24A2 | 1.54E-05 | 1.20 | 0.26 | 3.6E-07 | SLC24A2 | 0.0013847 | 3.82 | 1.94 | 0.0001801 |
| SYTL2 | 0.001115 | 1.31 | 0.39 | 5.4E-05 | SYTL2 | 1.413E-06 | 3.69 | 1.88 | 7.938E-08 |
| GRIP2 | 0.014703 | 1.85 | 0.89 | 0.0013 | GRIP2 | 0.0182456 | 3.30 | 1.72 | 0.0040381 |
| TTLL7 | 0.016543 | 1.11 | 0.15 | 0.00151 | TTLL7 | 0.0151508 | 3.26 | 1.71 | 0.0031965 |
| MACROD2 | 0.006028 | 1.47 | 0.56 | 0.00042 | MACROD2 | 0.0234444 | 3.10 | 1.63 | 0.0055385 |
| GABBR2 | 0.001423 | 1.16 | 0.21 | 7.4E-05 | GABBR2 | 0.0212835 | 3.10 | 1.63 | 0.0048883 |
| ADGRB1 | 0.001221 | 1.35 | 0.43 | 6.1E-05 | ADGRB1 | 0.04739 | 3.09 | 1.63 | 0.0136107 |
| VIPR1 | 1.61E-05 | 1.93 | 0.95 | 3.8E-07 | VIPR1 | 0.0001756 | 3.07 | 1.62 | 1.649E-05 |
| DLGAP3 | 0.048941 | 1.15 | 0.20 | 0.00637 | DLGAP3 | 0.0002033 | 3.03 | 1.60 | 1.945E-05 |
| TENM4 | 0.041841 | 1.22 | 0.29 | 0.00516 | TENM4 | 0.0038987 | 2.99 | 1.58 | 0.0006154 |
| LDLRAD4 | 0.048871 | 1.32 | 0.40 | 0.00635 | LDLRAD4 | 0.0083802 | 2.92 | 1.55 | 0.0015514 |
| HCN1 | 2.15E-06 | 1.32 | 0.40 | 3.8E-08 | HCN1 | 0.039091 | 2.88 | 1.52 | 0.0106557 |
| PIP5K1C | 0.001071 | 1.21 | 0.27 | 5.2E-05 | PIP5K1C | 2.516E-11 | 2.84 | 1.50 | 6.456E-13 |
| NAT8L | 0.049855 | 1.15 | 0.20 | 0.00653 | NAT8L | 0.0002883 | 2.76 | 1.46 | 2.882E-05 |
| SEL1L3 | 0.00014 | 1.32 | 0.40 | 4.9E-06 | SEL1L3 | 2.806E-17 | 2.74 | 1.45 | 4.061E-19 |
| RYR2 | 4.04E-13 | 1.43 | 0.52 | 1.8E-15 | RYR2 | 0.0051346 | 2.70 | 1.44 | 0.0008589 |
| ADAM22 | 0.000127 | 1.20 | 0.26 | 4.4E-06 | ADAM22 | 0.000429 | 2.67 | 1.42 | 4.592E-05 |
| FMNL1 | 0.000117 | 1.56 | 0.64 | 3.9E-06 | FMNL1 | 0.0080021 | 2.61 | 1.39 | 0.0014639 |
| ATP6 | 1.23E-05 | 2.09 | 1.07 | 2.8E-07 | ATP6 | 0.0026467 | 2.60 | 1.38 | 0.0003869 |
| SATB2 | 0.001013 | 1.56 | 0.64 | 4.9E-05 | SATB2 | 0.0101189 | 2.57 | 1.36 | 0.0019573 |
| KLF10 | 0.000697 | 1.45 | 0.54 | 3.2E-05 | KLF10 | 1.9E-18 | 2.56 | 1.35 | 2.494E-20 |
| SPEG | 0.003918 | 1.34 | 0.42 | 0.00025 | SPEG | 1.84E-09 | 2.53 | 1.34 | 6.12E-11 |
| SPRY4 | 0.018129 | 1.58 | 0.66 | 0.0017 | SPRY4 | 0.0007587 | 2.48 | 1.31 | 8.929E-05 |
| MCF2L | 2.76E-06 | 1.39 | 0.47 | 5E-08 | MCF2L | 0.0020186 | 2.45 | 1.29 | 0.0002796 |
| UNC13A | 2.02E-06 | 1.32 | 0.40 | 3.6E-08 | UNC13A | 0.0085817 | 2.41 | 1.27 | 0.0015968 |
| KRI1 | 0.002214 | 1.83 | 0.87 | 0.00013 | KRI1 | 0.0047571 | 2.32 | 1.22 | 0.0007834 |
| ANK1 | 0.000292 | 1.55 | 0.63 | 1.1E-05 | ANK1 | 0.0098888 | 2.29 | 1.19 | 0.0019039 |
| CROCC | 0.000195 | 2.44 | 1.29 | 7.1E-06 | CROCC | 0.0001698 | 2.26 | 1.18 | 1.584E-05 |
| WDR90 | 0.012501 | 2.11 | 1.08 | 0.00106 | WDR90 | 0.0059241 | 2.25 | 1.17 | 0.0010213 |
| XKR6 | 0.038679 | 1.62 | 0.70 | 0.00464 | XKR6 | 0.016869 | 2.24 | 1.17 | 0.0036697 |
| CACNA1A | 5.09E-05 | 1.42 | 0.51 | 1.5E-06 | CACNA1A | 0.0092437 | 2.23 | 1.16 | 0.0017484 |
| SRSF1 | 0.01834 | 1.18 | 0.24 | 0.00173 | SRSF1 | 0.0349864 | 2.15 | 1.10 | 0.0092515 |
| SHANK2 | 0.000667 | 1.30 | 0.38 | 3E-05 | SHANK2 | 0.0024375 | 2.13 | 1.09 | 0.0003509 |
| TMEM151B | 0.016065 | 1.17 | 0.23 | 0.00146 | TMEM151B | 0.0032252 | 2.13 | 1.09 | 0.0004898 |
| PLEKHG5 | 0.001013 | 1.61 | 0.69 | 4.9E-05 | PLEKHG5 | 0.0033819 | 2.09 | 1.07 | 0.0005198 |
| NDOR1 | 0.001488 | 2.02 | 1.01 | 7.8E-05 | NDOR1 | 0.004558 | 2.06 | 1.04 | 0.0007427 |
| RPAP1 | 0.017347 | 1.55 | 0.63 | 0.0016 | RPAP1 | 0.0001759 | 2.05 | 1.04 | 1.656E-05 |
| TMEM178B | 7.9E-06 | 1.27 | 0.34 | 1.7E-07 | TMEM178B | 0.0049041 | 2.05 | 1.03 | 0.000813 |
| CAMTA2 | 0.020927 | 1.30 | 0.38 | 0.00206 | CAMTA2 | 0.0457369 | 2.04 | 1.03 | 0.0130177 |
| ITPR1 | 0.039946 | 1.13 | 0.18 | 0.00484 | ITPR1 | 0.0052963 | 2.04 | 1.03 | 0.000888 |
| BCOR | 0.001642 | 1.46 | 0.55 | 8.7E-05 | BCOR | 0.000107 | 2.02 | 1.01 | 9.419E-06 |
| RNPC3 | 0.046367 | 1.34 | 0.42 | 0.00592 | RNPC3 | 0.0172103 | 1.98 | 0.99 | 0.0037637 |
| GTPBP2 | 0.002704 | 1.45 | 0.53 | 0.00016 | GTPBP2 | 1.113E-05 | 1.98 | 0.99 | 7.635E-07 |
| CSNK1D | 0.004303 | 1.42 | 0.51 | 0.00028 | CSNK1D | 0.0102423 | 1.96 | 0.97 | 0.0019888 |
| DOT1L | 0.00034 | 2.10 | 1.07 | 1.4E-05 | DOT1L | 0.0043945 | 1.96 | 0.97 | 0.0007137 |
| PFKFB3 | 0.000575 | 1.71 | 0.78 | 2.5E-05 | PFKFB3 | 0.003769 | 1.96 | 0.97 | 0.0005912 |

***Supplementary Table 2 continued***

| | | | | | | | | | |
|---|---|---|---|---|---|---|---|---|---|
| KCNF1 | 0.000863 | 1.27 | 0.35 | 4.1E-05 | KCNF1 | 0.0063629 | 1.94 | 0.96 | 0.0011163 |
| SCN8A | 7.82E-07 | 1.34 | 0.42 | 1.2E-08 | SCN8A | 0.0175234 | 1.92 | 0.94 | 0.0038469 |
| ANKRD10 | 0.008268 | 1.33 | 0.41 | 0.00062 | ANKRD10 | 1.782E-05 | 1.90 | 0.92 | 1.278E-06 |
| RNF166 | 0.00835 | 1.62 | 0.70 | 0.00063 | RNF166 | 0.0005565 | 1.89 | 0.92 | 6.22E-05 |
| TSC2 | 0.021333 | 1.26 | 0.34 | 0.0021 | TSC2 | 0.0002975 | 1.88 | 0.91 | 2.996E-05 |
| RSRP1 | 0.031407 | 1.23 | 0.30 | 0.00346 | RSRP1 | 0.006173 | 1.80 | 0.84 | 0.0010734 |
| CCNL1 | 0.00041 | 1.52 | 0.60 | 1.7E-05 | CCNL1 | 7.624E-05 | 1.80 | 0.84 | 6.445E-06 |
| CLK4 | 0.009178 | 1.29 | 0.37 | 0.00071 | CLK4 | 0.0203863 | 1.79 | 0.84 | 0.0046417 |
| BICDL1 | 0.000705 | 1.42 | 0.51 | 3.2E-05 | BICDL1 | 0.0363753 | 1.77 | 0.83 | 0.0097222 |
| MTCL1 | 4.13E-09 | 1.60 | 0.68 | 3.8E-11 | MTCL1 | 0.0014152 | 1.77 | 0.82 | 0.0001847 |
| TBC1D24 | 0.000637 | 1.50 | 0.58 | 2.8E-05 | TBC1D24 | 0.0032705 | 1.77 | 0.82 | 0.0004976 |
| MBP | 1.27E-19 | 1.34 | 0.42 | 2.5E-22 | MBP | 0.0005709 | 1.77 | 0.82 | 6.41E-05 |
| DGKH | 0.000143 | 1.27 | 0.34 | 5E-06 | DGKH | 0.0214736 | 1.75 | 0.81 | 0.0049416 |
| SYNE1 | 5.06E-09 | 1.24 | 0.31 | 4.7E-11 | SYNE1 | 0.0459297 | 1.69 | 0.75 | 0.0130931 |
| VEGFA | 0.000414 | 1.44 | 0.53 | 1.7E-05 | VEGFA | 0.0001823 | 1.68 | 0.75 | 1.723E-05 |
| BANP | 0.004061 | 1.51 | 0.59 | 0.00026 | BANP | 0.0400106 | 1.66 | 0.73 | 0.0109867 |
| CCDC88C | 0.018937 | 1.56 | 0.64 | 0.0018 | CCDC88C | 0.0139485 | 1.65 | 0.72 | 0.0028908 |
| TMEM132B | 0.04219 | 1.15 | 0.20 | 0.00521 | TMEM132B | 0.0377595 | 1.64 | 0.71 | 0.0101914 |
| MSANTD2 | 0.042232 | 1.70 | 0.77 | 0.00522 | MSANTD2 | 0.0092479 | 1.62 | 0.70 | 0.0017502 |
| CIZ1 | 0.024807 | 1.62 | 0.70 | 0.00254 | CIZ1 | 0.0076742 | 1.62 | 0.69 | 0.0013924 |
| FOXP1 | 0.028433 | 1.23 | 0.29 | 0.00303 | FOXP1 | 0.0041995 | 1.60 | 0.68 | 0.0006743 |
| ARHGEF2 | 0.000293 | 1.28 | 0.36 | 1.1E-05 | ARHGEF2 | 0.0054248 | 1.59 | 0.67 | 0.0009148 |
| PRPF38B | 0.000124 | 1.64 | 0.72 | 4.2E-06 | PRPF38B | 0.0148116 | 1.58 | 0.66 | 0.0031109 |
| FRMD6 | 7.83E-06 | 1.69 | 0.75 | 1.7E-07 | FRMD6 | 0.000403 | 1.57 | 0.65 | 4.254E-05 |
| PPP1R12C | 4.82E-05 | 1.39 | 0.48 | 1.4E-06 | PPP1R12C | 0.0352197 | 1.56 | 0.65 | 0.009322 |
| VPS8 | 0.033992 | 1.29 | 0.36 | 0.00382 | VPS8 | 0.009578 | 1.56 | 0.64 | 0.0018312 |
| CDC37L1 | 5.4E-07 | 1.32 | 0.40 | 8.3E-09 | CDC37L1 | 0.0021142 | 1.54 | 0.63 | 0.0002966 |
| OSBPL3 | 8.02E-20 | 2.32 | 1.21 | 1.5E-22 | OSBPL3 | 0.0210022 | 1.50 | 0.58 | 0.0048118 |
| DUSP11 | 1.45E-17 | 2.06 | 1.04 | 3.7E-20 | DUSP11 | 0.0308106 | 1.45 | 0.54 | 0.0078134 |
| PGGT1B | 0.024692 | 1.46 | 0.54 | 0.00252 | PGGT1B | 0.0430971 | 1.44 | 0.52 | 0.0120713 |
| CHIC1 | 0.004443 | 1.26 | 0.33 | 0.00029 | CHIC1 | 0.01951 | 1.43 | 0.51 | 0.0043927 |
| TIPARP | 0.037995 | 1.34 | 0.42 | 0.00452 | TIPARP | 0.0017011 | 1.42 | 0.51 | 0.0002288 |
| RIMS2 | 3.41E-07 | 1.33 | 0.41 | 4.9E-09 | RIMS2 | 0.0264193 | 1.42 | 0.51 | 0.0064328 |
| PACS2 | 1.62E-06 | 1.47 | 0.55 | 2.8E-08 | PACS2 | 0.0053556 | 1.42 | 0.51 | 0.0008998 |
| ANKRD11 | 0.017175 | 1.21 | 0.27 | 0.00158 | ANKRD11 | 0.0245813 | 1.42 | 0.51 | 0.0058694 |
| SNTB2 | 0.001585 | 1.44 | 0.53 | 8.4E-05 | SNTB2 | 0.0151992 | 1.40 | 0.49 | 0.003209 |
| GLS | 2.02E-08 | 1.31 | 0.39 | 2.2E-10 | GLS | 0.0461534 | 1.39 | 0.47 | 0.0131683 |
| CDK13 | 0.000152 | 1.42 | 0.51 | 5.4E-06 | CDK13 | 0.0322915 | 1.34 | 0.42 | 0.0083212 |
| METTL17 | 0.00282 | 1.75 | 0.80 | 0.00017 | METTL17 | 0.042506 | 1.34 | 0.42 | 0.0118613 |
| LUC7L2 | 4.96E-12 | 1.51 | 0.60 | 2.8E-14 | LUC7L2 | 0.0393655 | 1.28 | 0.36 | 0.0107564 |
| LITAF | 0.04035 | 1.49 | 0.58 | 0.0049 | LITAF | 0.0152451 | 1.27 | 0.34 | 0.0032212 |
| G3BP2 | 0.00905 | 0.87 | -0.20 | 0.0007 | G3BP2 | 0.0242621 | 0.83 | -0.28 | 0.0057715 |
| RPL23A | 0.001368 | 0.84 | -0.25 | 7E-05 | RPL23A | 0.0499191 | 0.81 | -0.30 | 0.0145729 |
| ADD3 | 0.01187 | 0.85 | -0.24 | 0.00099 | ADD3 | 0.0155708 | 0.81 | -0.31 | 0.0033107 |
| RPS15 | 5.81E-05 | 0.79 | -0.34 | 1.7E-06 | RPS15 | 0.00975 | 0.78 | -0.35 | 0.0018723 |
| ARPC5 | 0.000344 | 0.75 | -0.42 | 1.4E-05 | ARPC5 | 0.0195854 | 0.78 | -0.35 | 0.0044165 |
| UBC | 0.011853 | 0.85 | -0.23 | 0.00099 | UBC | 6.29E-05 | 0.78 | -0.35 | 5.183E-06 |
| PRPF31 | 0.031019 | 0.75 | -0.41 | 0.00339 | PRPF31 | 0.0379638 | 0.77 | -0.37 | 0.0102654 |
| SLC39A6 | 0.00271 | 0.70 | -0.51 | 0.00016 | SLC39A6 | 0.0219934 | 0.77 | -0.37 | 0.0051071 |

***Supplementary Table 2 continued***

| | | | | | | | | | |
|---|---|---|---|---|---|---|---|---|---|
| KLHL9 | 0.020927 | 0.86 | -0.21 | 0.00206 | KLHL9 | 0.0046812 | 0.77 | -0.37 | 0.0007661 |
| MLLT11 | 0.002377 | 0.80 | -0.31 | 0.00014 | MLLT11 | 0.0202468 | 0.77 | -0.38 | 0.0045978 |
| SUB1 | 0.03265 | 0.88 | -0.18 | 0.00363 | SUB1 | 0.0459747 | 0.77 | -0.38 | 0.0131105 |
| SKP1 | 0.001389 | 0.86 | -0.21 | 7.1E-05 | SKP1 | 0.0059077 | 0.77 | -0.38 | 0.0010179 |
| RPL13A | 4.64E-07 | 0.79 | -0.34 | 6.9E-09 | RPL13A | 0.0037235 | 0.77 | -0.39 | 0.0005832 |
| DDOST | 0.040584 | 0.81 | -0.30 | 0.00495 | DDOST | 0.0085306 | 0.76 | -0.39 | 0.0015843 |
| RBM3 | 0.000816 | 0.74 | -0.43 | 3.8E-05 | RBM3 | 0.0133629 | 0.75 | -0.41 | 0.0027417 |
| SELENOW | 4.76E-05 | 0.82 | -0.29 | 1.4E-06 | SELENOW | 0.0430236 | 0.75 | -0.41 | 0.01204 |
| PTMA | 0.014043 | 0.88 | -0.19 | 0.00123 | PTMA | 0.0024321 | 0.75 | -0.42 | 0.0003499 |
| PSMB1 | 0.014604 | 0.81 | -0.30 | 0.00129 | PSMB1 | 0.0121034 | 0.75 | -0.42 | 0.0024347 |
| MDH2 | 0.006132 | 0.80 | -0.32 | 0.00043 | MDH2 | 0.0181328 | 0.75 | -0.42 | 0.0040029 |
| NACA | 0.032895 | 0.86 | -0.21 | 0.00366 | NACA | 0.0023342 | 0.75 | -0.42 | 0.0003326 |
| UBB | 1.09E-09 | 0.76 | -0.40 | 9E-12 | UBB | 0.0111393 | 0.75 | -0.42 | 0.0021962 |
| PPP1CB | 0.002209 | 0.86 | -0.22 | 0.00012 | PPP1CB | 0.0062732 | 0.75 | -0.42 | 0.0010964 |
| VPS35L | 0.002284 | 0.74 | -0.44 | 0.00013 | VPS35L | 0.0178344 | 0.75 | -0.42 | 0.0039285 |
| SLC7A11 | 0.005745 | 0.80 | -0.33 | 0.0004 | SLC7A11 | 0.0171146 | 0.74 | -0.43 | 0.0037358 |
| AP2M1 | 7.04E-05 | 0.84 | -0.25 | 2.2E-06 | AP2M1 | 0.0001252 | 0.74 | -0.44 | 1.128E-05 |
| USP22 | 0.001455 | 0.81 | -0.31 | 7.5E-05 | USP22 | 0.039975 | 0.74 | -0.44 | 0.0109685 |
| PKM | 3.98E-05 | 0.87 | -0.21 | 1.1E-06 | PKM | 0.0028767 | 0.74 | -0.44 | 0.000427 |
| RPL41 | 1.63E-05 | 0.80 | -0.33 | 3.9E-07 | RPL41 | 0.0110108 | 0.74 | -0.44 | 0.0021665 |
| MDH1 | 0.002102 | 0.84 | -0.26 | 0.00012 | MDH1 | 0.0008432 | 0.73 | -0.45 | 0.0001009 |
| RPL3 | 3.3E-184 | 0.20 | -2.35 | 3E-188 | RPL3 | 0.0019457 | 0.73 | -0.45 | 0.0002677 |
| TMEM47 | 0.018937 | 0.84 | -0.25 | 0.0018 | TMEM47 | 0.0012355 | 0.73 | -0.46 | 0.0001581 |
| USP11 | 0.001273 | 0.79 | -0.33 | 6.4E-05 | USP11 | 0.0404896 | 0.73 | -0.46 | 0.0111517 |
| DCTN2 | 0.018476 | 0.84 | -0.26 | 0.00175 | DCTN2 | 0.0030497 | 0.73 | -0.46 | 0.0004566 |
| ENSA | 0.011451 | 0.88 | -0.19 | 0.00094 | ENSA | 0.0012211 | 0.72 | -0.48 | 0.0001559 |
| MAP2K1 | 2.3E-06 | 0.79 | -0.35 | 4.1E-08 | MAP2K1 | 0.0282057 | 0.72 | -0.48 | 0.0069946 |
| FKBP8 | 0.010975 | 0.83 | -0.27 | 0.00089 | FKBP8 | 0.0002871 | 0.71 | -0.49 | 2.866E-05 |
| RANGAP1 | 0.023577 | 0.85 | -0.23 | 0.00238 | RANGAP1 | 0.0008992 | 0.71 | -0.49 | 0.0001089 |
| MIF | 0.032977 | 0.78 | -0.36 | 0.00368 | MIF | 0.0011828 | 0.70 | -0.51 | 0.0001493 |
| KIFAP3 | 0.017927 | 0.88 | -0.19 | 0.00167 | KIFAP3 | 0.0223958 | 0.70 | -0.51 | 0.0052321 |
| COX4I1 | 0.000856 | 0.85 | -0.23 | 4E-05 | COX4I1 | 0.0014832 | 0.70 | -0.51 | 0.0001954 |
| PPP1CA | 0.037355 | 0.87 | -0.20 | 0.00441 | PPP1CA | 0.0040166 | 0.70 | -0.51 | 0.0006379 |
| SEC23A | 0.026773 | 0.87 | -0.20 | 0.00282 | SEC23A | 2.567E-05 | 0.70 | -0.52 | 1.922E-06 |
| RPL13 | 9.69E-05 | 0.76 | -0.39 | 3.1E-06 | RPL13 | 8.976E-05 | 0.69 | -0.53 | 7.718E-06 |
| MATN2 | 9.79E-05 | 0.58 | -0.78 | 3.2E-06 | MATN2 | 0.0286295 | 0.69 | -0.53 | 0.0071317 |
| MAGED1 | 0.002556 | 0.87 | -0.21 | 0.00015 | MAGED1 | 0.0006434 | 0.69 | -0.53 | 7.358E-05 |
| TIMM8B | 0.046367 | 0.82 | -0.29 | 0.00592 | TIMM8B | 0.0242861 | 0.69 | -0.53 | 0.0057808 |
| GNB1 | 0.009263 | 0.87 | -0.20 | 0.00072 | GNB1 | 0.0020042 | 0.69 | -0.54 | 0.000277 |
| GNAS | 0.002753 | 0.89 | -0.17 | 0.00016 | GNAS | 0.0001154 | 0.69 | -0.54 | 1.028E-05 |
| TMSB10 | 0.02159 | 0.79 | -0.35 | 0.00214 | TMSB10 | 0.000101 | 0.69 | -0.54 | 8.794E-06 |
| RPS29 | 0.006021 | 0.78 | -0.36 | 0.00042 | RPS29 | 0.0083123 | 0.68 | -0.55 | 0.0015367 |
| UQCRC1 | 0.006777 | 0.84 | -0.24 | 0.00049 | UQCRC1 | 0.0005936 | 0.68 | -0.55 | 6.706E-05 |
| PRDX5 | 0.000348 | 0.77 | -0.37 | 1.4E-05 | PRDX5 | 0.0003972 | 0.68 | -0.55 | 4.183E-05 |
| DDAH1 | 0.043863 | 0.85 | -0.24 | 0.0055 | DDAH1 | 0.0001583 | 0.68 | -0.55 | 1.465E-05 |
| MRPL20 | 0.028856 | 0.74 | -0.44 | 0.00309 | MRPL20 | 0.0320652 | 0.68 | -0.56 | 0.0082407 |
| TOMM20 | 0.005967 | 0.86 | -0.22 | 0.00041 | TOMM20 | 2.587E-06 | 0.68 | -0.56 | 1.54E-07 |
| RAN | 0.043589 | 0.87 | -0.20 | 0.00546 | RAN | 0.0037511 | 0.68 | -0.57 | 0.0005881 |
| AP2S1 | 0.042476 | 0.83 | -0.27 | 0.00527 | AP2S1 | 0.0030845 | 0.67 | -0.57 | 0.0004639 |

***Supplementary Table 2 continued***

| | | | | | | | | | |
|---|---|---|---|---|---|---|---|---|---|
| PPP2R1A | 0.001862 | 0.87 | -0.20 | 0.0001 | PPP2R1A | 0.0062889 | 0.67 | -0.57 | 0.0010998 |
| PEBP1 | 7.27E-10 | 0.74 | -0.43 | 5.7E-12 | PEBP1 | 4.167E-07 | 0.67 | -0.58 | 2.115E-08 |
| PKP2 | 0.005611 | 0.46 | -1.11 | 0.00038 | PKP2 | 9.061E-07 | 0.67 | -0.59 | 4.9E-08 |
| ERH | 0.0192 | 0.60 | -0.73 | 0.00184 | ERH | 0.0003916 | 0.67 | -0.59 | 4.114E-05 |
| PQBP1 | 0.025317 | 0.76 | -0.39 | 0.00261 | PQBP1 | 0.033213 | 0.67 | -0.59 | 0.0086323 |
| NOTCH2 | 0.030497 | 0.74 | -0.44 | 0.00333 | NOTCH2 | 6.781E-05 | 0.66 | -0.59 | 5.675E-06 |
| YWHAZ | 0.00252 | 0.84 | -0.25 | 0.00015 | YWHAZ | 0.0003215 | 0.66 | -0.60 | 3.276E-05 |
| PJA2 | 0.039187 | 0.91 | -0.14 | 0.00472 | PJA2 | 1.089E-06 | 0.66 | -0.60 | 5.97E-08 |
| TUBB2A | 2.86E-14 | 0.73 | -0.44 | 1E-16 | TUBB2A | 8.438E-06 | 0.66 | -0.60 | 5.635E-07 |
| TSPAN13 | 0.010953 | 0.86 | -0.23 | 0.00089 | TSPAN13 | 0.0010961 | 0.66 | -0.60 | 0.0001367 |
| EID1 | 0.000774 | 0.84 | -0.25 | 3.6E-05 | EID1 | 1.709E-05 | 0.65 | -0.61 | 1.221E-06 |
| DYNLL1 | 2.64E-07 | 0.76 | -0.39 | 3.7E-09 | DYNLL1 | 3.409E-05 | 0.65 | -0.62 | 2.633E-06 |
| COX5B | 0.008106 | 0.74 | -0.43 | 0.00061 | COX5B | 0.0001632 | 0.65 | -0.62 | 1.516E-05 |
| CALM3 | 0.022943 | 0.90 | -0.15 | 0.0023 | CALM3 | 5.36E-05 | 0.65 | -0.63 | 4.333E-06 |
| LDHB | 0.047806 | 0.85 | -0.23 | 0.00617 | LDHB | 1.195E-05 | 0.64 | -0.63 | 8.243E-07 |
| PSMB5 | 0.012939 | 0.68 | -0.56 | 0.0011 | PSMB5 | 0.0004334 | 0.64 | -0.63 | 4.651E-05 |
| CNN3 | 0.029969 | 0.80 | -0.33 | 0.00325 | CNN3 | 1.609E-08 | 0.64 | -0.65 | 6.151E-10 |
| YWHAH | 5.15E-05 | 0.85 | -0.23 | 1.5E-06 | YWHAH | 1.608E-08 | 0.64 | -0.65 | 6.141E-10 |
| PGAM1 | 0.018546 | 0.88 | -0.18 | 0.00175 | PGAM1 | 1.555E-07 | 0.63 | -0.66 | 7.345E-09 |
| SDCBP | 0.000187 | 0.81 | -0.30 | 6.8E-06 | SDCBP | 0.0090205 | 0.63 | -0.67 | 0.0016975 |
| ELOB | 0.006806 | 0.78 | -0.36 | 0.00049 | ELOB | 2.275E-05 | 0.63 | -0.67 | 1.679E-06 |
| CALM2 | 0.00106 | 0.89 | -0.16 | 5.1E-05 | CALM2 | 5.78E-07 | 0.63 | -0.68 | 2.992E-08 |
| MZT1 | 0.024547 | 0.83 | -0.26 | 0.0025 | MZT1 | 0.0005642 | 0.62 | -0.68 | 6.324E-05 |
| RTN3 | 0.002619 | 0.89 | -0.17 | 0.00015 | RTN3 | 0.0001796 | 0.62 | -0.68 | 1.694E-05 |
| PSMA2 | 0.037843 | 0.83 | -0.26 | 0.00449 | PSMA2 | 5.657E-06 | 0.62 | -0.69 | 3.648E-07 |
| PRXL2A | 0.007675 | 0.79 | -0.34 | 0.00057 | PRXL2A | 2.529E-07 | 0.61 | -0.70 | 1.247E-08 |
| YWHAE | 0.003592 | 0.83 | -0.27 | 0.00022 | YWHAE | 0.0005548 | 0.61 | -0.70 | 6.19E-05 |
| SUMO2 | 0.027035 | 0.85 | -0.24 | 0.00285 | SUMO2 | 7.856E-06 | 0.61 | -0.71 | 5.203E-07 |
| FAM131A | 0.000224 | 0.82 | -0.29 | 8.3E-06 | FAM131A | 0.0069186 | 0.61 | -0.71 | 0.0012295 |
| ARF1 | 0.000275 | 0.81 | -0.31 | 1.1E-05 | ARF1 | 2.352E-05 | 0.61 | -0.71 | 1.744E-06 |
| SPAG7 | 0.015507 | 0.67 | -0.58 | 0.00138 | SPAG7 | 3.336E-05 | 0.61 | -0.72 | 2.563E-06 |
| YWHAB | 0.000122 | 0.82 | -0.29 | 4.1E-06 | YWHAB | 0.0058528 | 0.61 | -0.72 | 0.0010046 |
| ORAI2 | 0.0023 | 0.70 | -0.51 | 0.00013 | ORAI2 | 0.0053182 | 0.61 | -0.72 | 0.0008928 |
| PPIB | 0.026594 | 0.78 | -0.35 | 0.00278 | PPIB | 1.198E-05 | 0.60 | -0.73 | 8.273E-07 |
| ATP6V0E2 | 0.03265 | 0.87 | -0.19 | 0.00363 | ATP6V0E2 | 0.0090914 | 0.60 | -0.74 | 0.0017151 |
| CLU | 2.4E-13 | 0.75 | -0.41 | 1.1E-15 | CLU | 0.002862 | 0.59 | -0.75 | 0.0004244 |
| RABAC1 | 0.00213 | 0.74 | -0.44 | 0.00012 | RABAC1 | 0.0008638 | 0.59 | -0.76 | 0.0001041 |
| PDXP | 0.034106 | 0.85 | -0.24 | 0.00384 | PDXP | 0.0026643 | 0.58 | -0.77 | 0.0003898 |
| SWI5 | 0.031673 | 0.81 | -0.30 | 0.00349 | SWI5 | 0.0062635 | 0.58 | -0.79 | 0.0010944 |
| PSMC3 | 0.002082 | 0.82 | -0.29 | 0.00012 | PSMC3 | 7.728E-08 | 0.58 | -0.79 | 3.455E-09 |
| NDN | 3.36E-09 | 0.69 | -0.53 | 3E-11 | NDN | 0.0009312 | 0.58 | -0.79 | 0.0001133 |
| MRFAP1 | 0.000554 | 0.83 | -0.28 | 2.4E-05 | MRFAP1 | 6.134E-05 | 0.58 | -0.80 | 5.045E-06 |
| HSBP1 | 0.026202 | 0.85 | -0.23 | 0.00272 | HSBP1 | 1.045E-06 | 0.57 | -0.81 | 5.703E-08 |
| NAGA | 0.026222 | 0.48 | -1.07 | 0.00273 | NAGA | 0.0010103 | 0.57 | -0.81 | 0.0001241 |
| RPLP1 | 0.000106 | 0.75 | -0.41 | 3.5E-06 | RPLP1 | 4.905E-05 | 0.57 | -0.82 | 3.929E-06 |
| PTPA | 0.016874 | 0.79 | -0.35 | 0.00154 | PTPA | 4.294E-10 | 0.56 | -0.83 | 1.277E-11 |
| RPL8 | 0.000184 | 0.78 | -0.36 | 6.6E-06 | RPL8 | 3.699E-08 | 0.56 | -0.83 | 1.54E-09 |
| TUBA1A | 1.16E-08 | 0.81 | -0.30 | 1.2E-10 | TUBA1A | 7.348E-06 | 0.56 | -0.85 | 4.838E-07 |
| PRDX2 | 0.001262 | 0.79 | -0.34 | 6.3E-05 | PRDX2 | 1.203E-06 | 0.56 | -0.85 | 6.682E-08 |

*Supplementary Table 2 continued*

| | | | | | | | | | |
|---|---|---|---|---|---|---|---|---|---|
| CRIM1 | 0.000344 | 0.73 | -0.45 | 1.4E-05 | CRIM1 | 1.237E-06 | 0.55 | -0.86 | 6.88E-08 |
| COX8A | 0.037129 | 0.86 | -0.22 | 0.00437 | COX8A | 1.069E-05 | 0.55 | -0.86 | 7.298E-07 |
| ALDH4A1 | 0.045635 | 0.70 | -0.51 | 0.00579 | ALDH4A1 | 0.0043223 | 0.55 | -0.87 | 0.0006987 |
| POLR2F | 0.043497 | 0.66 | -0.61 | 0.00544 | POLR2F | 0.0004057 | 0.54 | -0.89 | 4.293E-05 |
| NSG1 | 0.009491 | 0.80 | -0.32 | 0.00074 | NSG1 | 5.796E-05 | 0.54 | -0.90 | 4.726E-06 |
| POLR2G | 0.007635 | 0.71 | -0.49 | 0.00056 | POLR2G | 0.0005488 | 0.54 | -0.90 | 6.102E-05 |
| RRP1 | 0.001088 | 0.76 | -0.40 | 5.3E-05 | RRP1 | 0.0001017 | 0.53 | -0.93 | 8.862E-06 |
| CTSL | 0.002036 | 0.77 | -0.38 | 0.00011 | CTSL | 3.29E-07 | 0.53 | -0.93 | 1.647E-08 |
| AHNAK | 0.036725 | 0.80 | -0.33 | 0.0043 | AHNAK | 0.0011316 | 0.52 | -0.94 | 0.0001416 |
| ID4 | 0.005593 | 0.80 | -0.32 | 0.00038 | ID4 | 8.536E-11 | 0.51 | -0.97 | 2.335E-12 |
| SNRPN | 0.011901 | 0.87 | -0.20 | 0.00099 | SNRPN | 2.346E-11 | 0.51 | -0.97 | 5.948E-13 |
| CST3 | 6.65E-14 | 0.75 | -0.42 | 2.7E-16 | CST3 | 9.825E-08 | 0.51 | -0.97 | 4.48E-09 |
| SPOPL | 0.035215 | 0.75 | -0.41 | 0.00405 | SPOPL | 2.411E-07 | 0.50 | -1.00 | 1.181E-08 |
| APOE | 1.01E-18 | 0.69 | -0.53 | 2.1E-21 | APOE | 1.319E-20 | 0.50 | -1.00 | 1.456E-22 |
| GPRIN1 | 0.047361 | 0.85 | -0.24 | 0.00609 | GPRIN1 | 0.0033871 | 0.49 | -1.02 | 0.0005209 |
| NDUFAF8 | 0.038511 | 0.54 | -0.89 | 0.00461 | NDUFAF8 | 0.0001093 | 0.49 | -1.03 | 9.675E-06 |
| PTPN14 | 0.016892 | 0.49 | -1.02 | 0.00154 | PTPN14 | 0.0035535 | 0.49 | -1.03 | 0.0005516 |
| ARL6IP1 | 0.034486 | 0.88 | -0.19 | 0.00391 | ARL6IP1 | 1.71E-18 | 0.49 | -1.03 | 2.211E-20 |
| TIMP3 | 0.002036 | 0.79 | -0.34 | 0.00011 | TIMP3 | 4.886E-08 | 0.48 | -1.05 | 2.089E-09 |
| TUSC3 | 4.93E-05 | 0.79 | -0.34 | 1.4E-06 | TUSC3 | 3.718E-13 | 0.48 | -1.06 | 7.765E-15 |
| TUBA1B | 0.000235 | 0.85 | -0.24 | 8.8E-06 | TUBA1B | 1.609E-10 | 0.47 | -1.08 | 4.592E-12 |
| FANCE | 0.029969 | 0.51 | -0.97 | 0.00325 | FANCE | 1.82E-06 | 0.47 | -1.09 | 1.046E-07 |
| DSC3 | 0.021357 | 0.28 | -1.85 | 0.0021 | DSC3 | 0.0015792 | 0.47 | -1.10 | 0.0002105 |
| CPE | 1.68E-10 | 0.79 | -0.33 | 1.2E-12 | CPE | 3.99E-12 | 0.46 | -1.11 | 9.264E-14 |
| ARHGEF26 | 0.015596 | 0.62 | -0.69 | 0.0014 | ARHGEF26 | 0.0257007 | 0.46 | -1.12 | 0.0062108 |
| NNAT | 3.46E-45 | 0.35 | -1.52 | 1.6E-48 | NNAT | 4.805E-05 | 0.46 | -1.13 | 3.845E-06 |
| USP46 | 0.031385 | 0.86 | -0.21 | 0.00345 | USP46 | 1.749E-06 | 0.46 | -1.13 | 1.002E-07 |
| FXYD6 | 5.21E-06 | 0.64 | -0.64 | 1E-07 | FXYD6 | 4.118E-08 | 0.46 | -1.13 | 1.732E-09 |
| NCALD | 0.010902 | 0.87 | -0.21 | 0.00088 | NCALD | 0.0005542 | 0.45 | -1.14 | 6.178E-05 |
| ALDOC | 3.28E-14 | 0.73 | -0.44 | 1.2E-16 | ALDOC | 0.0009167 | 0.45 | -1.16 | 0.0001112 |
| GEMIN4 | 0.024882 | 0.41 | -1.28 | 0.00255 | GEMIN4 | 0.0001388 | 0.41 | -1.27 | 1.265E-05 |
| ALDH6A1 | 0.010775 | 0.78 | -0.35 | 0.00087 | ALDH6A1 | 9.081E-11 | 0.41 | -1.27 | 2.497E-12 |
| SLC44A5 | 0.006363 | 0.63 | -0.67 | 0.00045 | SLC44A5 | 0.0004748 | 0.40 | -1.32 | 5.161E-05 |
| SYP | 1.59E-07 | 0.84 | -0.25 | 2.1E-09 | SYP | 0.0019175 | 0.40 | -1.33 | 0.000263 |
| KCTD12 | 0.019252 | 0.78 | -0.37 | 0.00184 | KCTD12 | 2.305E-08 | 0.39 | -1.34 | 9.205E-10 |
| SPON1 | 0.033627 | 0.77 | -0.38 | 0.00376 | SPON1 | 0.0439716 | 0.39 | -1.37 | 0.0123819 |
| TMEM35A | 0.04332 | 0.76 | -0.39 | 0.00541 | TMEM35A | 0.0183549 | 0.38 | -1.39 | 0.0040669 |
| ANGPT1 | 0.025846 | 0.55 | -0.86 | 0.00268 | ANGPT1 | 0.002084 | 0.38 | -1.41 | 0.0002912 |
| PKIA | 0.004228 | 0.84 | -0.24 | 0.00027 | PKIA | 5.78E-07 | 0.37 | -1.43 | 2.99E-08 |
| THBD | 0.042674 | 0.71 | -0.50 | 0.00531 | THBD | 0.0054813 | 0.37 | -1.44 | 0.0009265 |
| PLPP3 | 0.00032 | 0.82 | -0.29 | 1.3E-05 | PLPP3 | 1.544E-19 | 0.36 | -1.46 | 1.84E-21 |
| H2AZ1 | 0.002888 | 0.79 | -0.33 | 0.00017 | H2AZ1 | 2.043E-19 | 0.36 | -1.46 | 2.459E-21 |
| DNER | 7.04E-05 | 0.77 | -0.38 | 2.2E-06 | DNER | 0.0013438 | 0.36 | -1.47 | 0.0001739 |
| MCTP1 | 0.010176 | 0.80 | -0.32 | 0.00081 | MCTP1 | 0.0159999 | 0.36 | -1.48 | 0.0034321 |
| CNIH2 | 8.29E-07 | 0.70 | -0.52 | 1.3E-08 | CNIH2 | 0.0128917 | 0.36 | -1.48 | 0.0026269 |
| CADM1 | 0.001841 | 0.83 | -0.27 | 9.9E-05 | CADM1 | 1.333E-09 | 0.35 | -1.51 | 4.349E-11 |
| UBALD2 | 0.004611 | 0.55 | -0.86 | 0.00031 | UBALD2 | 0.0014473 | 0.35 | -1.53 | 0.0001898 |
| SGK1 | 0.000103 | 0.65 | -0.61 | 3.4E-06 | SGK1 | 3.699E-19 | 0.34 | -1.56 | 4.525E-21 |
| GSTA4 | 0.005188 | 0.74 | -0.44 | 0.00035 | GSTA4 | 8.339E-12 | 0.34 | -1.56 | 2.019E-13 |

***Supplementary Table 2 continued***

| | | | | | | | | | |
|---|---|---|---|---|---|---|---|---|---|
| APLP1 | 0.028949 | 0.90 | -0.15 | 0.00311 | APLP1 | 9.284E-05 | 0.34 | -1.57 | 8.032E-06 |
| TUBB2B | 0.007966 | 0.78 | -0.35 | 0.00059 | TUBB2B | 3.311E-37 | 0.34 | -1.57 | 1.366E-39 |
| KCNAB1 | 0.019404 | 0.83 | -0.27 | 0.00186 | KCNAB1 | 0.0007989 | 0.33 | -1.59 | 9.486E-05 |
| OLFM1 | 0.012276 | 0.87 | -0.21 | 0.00103 | OLFM1 | 7.646E-08 | 0.33 | -1.61 | 3.411E-09 |
| DACT2 | 0.008466 | 0.71 | -0.50 | 0.00064 | DACT2 | 1.21E-05 | 0.33 | -1.62 | 8.363E-07 |
| GPC4 | 0.018081 | 0.47 | -1.08 | 0.00169 | GPC4 | 1.069E-08 | 0.31 | -1.68 | 4.004E-10 |
| PLPPR4 | 0.031172 | 0.90 | -0.16 | 0.00342 | PLPPR4 | 7.614E-06 | 0.31 | -1.68 | 5.028E-07 |
| GPM6A | 2.4E-05 | 0.88 | -0.19 | 6.1E-07 | GPM6A | 3.29E-05 | 0.30 | -1.75 | 2.522E-06 |
| NR6A1 | 0.018426 | 0.41 | -1.30 | 0.00174 | NR6A1 | 2.389E-15 | 0.29 | -1.77 | 4.004E-17 |
| CNTN1 | 0.014604 | 0.91 | -0.14 | 0.00129 | CNTN1 | 1.565E-10 | 0.28 | -1.82 | 4.452E-12 |
| SNCA | 1.14E-06 | 0.78 | -0.37 | 1.9E-08 | SNCA | 2.825E-14 | 0.27 | -1.91 | 5.253E-16 |
| RAB3B | 0.02159 | 0.66 | -0.61 | 0.00214 | RAB3B | 4.493E-21 | 0.26 | -1.93 | 4.691E-23 |
| RARRES2 | 0.010115 | 0.45 | -1.14 | 0.0008 | RARRES2 | 1.478E-12 | 0.26 | -1.95 | 3.278E-14 |
| FBLN1 | 0.006325 | 0.46 | -1.13 | 0.00044 | FBLN1 | 1.425E-12 | 0.26 | -1.96 | 3.153E-14 |
| H4C4 | 0.010689 | 0.73 | -0.44 | 0.00086 | H4C4 | 4.721E-21 | 0.25 | -1.99 | 4.953E-23 |
| AMPH | 0.033947 | 0.86 | -0.21 | 0.00381 | AMPH | 5.513E-07 | 0.25 | -2.00 | 2.829E-08 |
| ATP1A3 | 0.005036 | 0.88 | -0.19 | 0.00034 | ATP1A3 | 4.155E-08 | 0.24 | -2.05 | 1.754E-09 |
| TMEM158 | 0.021421 | 0.66 | -0.60 | 0.00211 | TMEM158 | 0.0001122 | 0.23 | -2.13 | 9.967E-06 |
| COL1A2 | 0.018931 | 0.73 | -0.46 | 0.0018 | COL1A2 | 3.473E-72 | 0.23 | -2.13 | 4.144E-75 |
| PPP1R1A | 0.0111 | 0.74 | -0.43 | 0.0009 | PPP1R1A | 0.0001108 | 0.22 | -2.17 | 9.839E-06 |
| ZWINT | 2.12E-10 | 0.79 | -0.33 | 1.6E-12 | ZWINT | 6.78E-24 | 0.19 | -2.40 | 5.63E-26 |
| SEZ6 | 0.00453 | 0.77 | -0.37 | 0.0003 | SEZ6 | 0.0454874 | 0.19 | -2.43 | 0.0129331 |
| TGFBR3 | 0.000513 | 0.48 | -1.05 | 2.2E-05 | TGFBR3 | 1.544E-16 | 0.18 | -2.50 | 2.365E-18 |
| GUCY1B1 | 0.027015 | 0.87 | -0.20 | 0.00284 | GUCY1B1 | 1.432E-08 | 0.17 | -2.53 | 5.446E-10 |
| VAT1L | 0.008017 | 0.69 | -0.54 | 0.0006 | VAT1L | 4.791E-51 | 0.16 | -2.68 | 1.191E-53 |
| VTN | 0.000451 | 0.72 | -0.46 | 1.9E-05 | VTN | 3.578E-09 | 0.13 | -2.89 | 1.258E-10 |
| RTN1 | 0.000521 | 0.84 | -0.25 | 2.2E-05 | RTN1 | 3.921E-12 | 0.13 | -2.96 | 9.067E-14 |
| SYNPR | 2.93E-06 | 0.65 | -0.62 | 5.4E-08 | SYNPR | 0.0205605 | 0.12 | -3.05 | 0.0046916 |
| NRP2 | 0.006496 | 0.67 | -0.58 | 0.00046 | NRP2 | 2.41E-29 | 0.10 | -3.26 | 1.498E-31 |
| NDNF | 0.00049 | 0.52 | -0.95 | 2.1E-05 | NDNF | 1.326E-10 | 0.08 | -3.67 | 3.693E-12 |
| SLC13A4 | 2.93E-06 | 0.51 | -0.97 | 5.4E-08 | SLC13A4 | 1.678E-19 | 0.08 | -3.70 | 2.011E-21 |
| GPC3 | 0.004103 | 0.29 | -1.77 | 0.00026 | GPC3 | 2.86E-74 | 0.07 | -3.88 | 3.128E-77 |
| LYPD1 | 1.77E-07 | 0.55 | -0.87 | 2.4E-09 | LYPD1 | 6.074E-86 | 0.07 | -3.93 | 3.926E-89 |
| SST | 0.002687 | 0.79 | -0.34 | 0.00016 | SST | 4.194E-14 | 0.01 | -7.27 | 7.966E-16 |
| GNG3 | 0.017878 | 0.77 | -0.38 | 0.00166 | GNG3 | 1.994E-06 | 0.00 | -8.08 | 1.16E-07 |

***Supplementary Table 3.***

Supplementary Table 3. Gene Ontology (GO) Molecular Function analysis of the 323 overlapping gene set detects significant enrichment in 12 categories of genes (FDR<0.05). Full GO Molecular Function analysis results are in Data Set 4.

| Term | Overlap | P-value | Adjusted P-value | Odds Ratio | Combined Score | Genes |
|---|---|---|---|---|---|---|
| Sodium Channel Activity (GO:0005272) | 7/40 | 3E-06 | 0.00079 | 13.27 | 168.60 | KCNK9;GPRIN1;SCN8A;SCN2A;SCN4B; SCN1A;HCN1 |
| Adenylate Cyclase Regulator Activity (GO:0010854) | 4/8 | 4.3E-06 | 0.00079 | 62.07 | 766.47 | GRM7;GNAS;CALM3;CALM2 |
| Cyclase Activator Activity (GO:0010853) | 3/5 | 4E-05 | 0.00488 | 92.82 | 939.93 | GNAS;CALM3;CALM2 |
| Ion Channel Regulator Activity (GO:0099106) | 9/119 | 0.00013 | 0.01192 | 5.13 | 45.91 | YWHAE;KCNF1;ENSA;PKP2;KCNAB1; CABP1;SGK1;SCN4B;YWHAH |
| Voltage-Gated Sodium Channel Activity (GO:0005248) | 4/18 | 0.00017 | 0.01221 | 17.72 | 154.18 | SCN8A;SCN2A;SCN4B;SCN1A |
| Phosphoserine Residue Binding (GO:0050815) | 3/9 | 0.00032 | 0.01953 | 30.93 | 248.90 | YWHAE;YWHAB;YWHAZ |
| Calcium Channel Inhibitor Activity (GO:0019855) | 3/11 | 0.00061 | 0.03211 | 23.20 | 171.55 | YWHAE;ITPR1;CALM2 |
| Calcium Channel Regulator Activity (GO:0005246) | 5/45 | 0.00075 | 0.03409 | 7.77 | 55.95 | YWHAE;ITPR1;CABP1;SGK1;CALM2 |
| Voltage-Gated Monoatomic Ion Channel Activity (GO:0005244) | 4/28 | 0.00098 | 0.03805 | 10.33 | 71.55 | GRIN2A;GPRIN1;SCN1A;HCN1 |
| Protein Kinase A Catalytic Subunit Binding (GO:0034236) | 3/13 | 0.00104 | 0.03805 | 18.56 | 127.46 | RYR2;PKIA;PJA2 |
| Protein Phosphatase Regulator Activity (GO:0019888) | 6/75 | 0.0013 | 0.04315 | 5.41 | 35.99 | YWHAB;PPP2R1A;ENSA;CALM3; PPP1R12C;CALM2 |
| Potassium Channel Regulator Activity (GO:0015459) | 5/53 | 0.00158 | 0.04809 | 6.47 | 41.75 | YWHAE;KCNF1;ENSA;KCNAB1;SGK1 |

#### *Supplementary Table 4*

Supplementary Table 4. DisGeNet analysis of disease links of the human genes in the 323 overlapping gene set with FDR<0.05. The full analysis is in Data Set 5.

| Term | Overlap | P-value | Adjusted P-value | Odds Ratio | Combined Score | Genes |
|---|---|---|---|---|---|---|
| Epilepsy | 52/1177 | 3.34E-11 | 1.43E-07 | 3.16 | 76.33 | YWHAE;GABRB2;RYR2;NRP2;CLU;PQBP1;KCNT1;TUBA1A;GRM7;PPP2R1A;SCN1A;COX8A;BRD2;MAP2K1;SNRPN;TSC2;KCNAB1;MIF;DDOST;TGFBR3;TUBB2B;TUBB2A;SCN8A;TBC1D24;SELENOW;SGK1;NAT8L;SATB2;ANKRD11;ATP6;ADAM22;SEZ6;CACNA1A;ATP1A3;CACNA1G;CNN3;PTPA;GRIN2A;UBB;APOE;GABBR2;MDH2;NAGA;ALDH4A1;CCDC88C;GNB1;GNAS;CALM3;BCOR;SCN2A;CALM2;HCN1 |
| Schizophrenia | 67/1923 | 1.06E-09 | 2.28E-06 | 2.51 | 51.92 | YWHAE;GABRB2;RPL3;TENM4;TUSC3;YWHAB;PEBP1;CLU;GLS;SYNE1;RIMS2;GRM7;PDE4A;DLGAP3;DLGAP2;TMSB10;YWHAH;USP46;MAP2K1;TSC2;CSNK1D;MIF;SYP;RANGAP1;YWHAZ;FOXP1;ARC;TUBB2B;MAG;NOS1AP;UQCRC1;GEMIN4;ALDOC;KCTD12;ARHGEF2;SHANK2;DGKH;GPM6A;RTN3;RTN1;ARHGEF26;SATB2;ATP6;SRSF1;PRDX2;GRIN2A;NDOR1;MOBP;RPL13;MBP;APOE;GABBR2;MDH1;SPRY4;ATP2B2;VEGFA;SST;CCDC88C;GNB1;PLXNB3;GNAS;CALM3;FXYD6;SCN2A;CALM2;XKR6;HCN1 |
| Seizures | 47/1174 | 7.94E-09 | 1.13E-05 | 2.80 | 52.28 | YWHAE;RYR2;NRP2;SATB2;ANKRD11;ATP6;ADAM22;CACNA1A;ATP1A3;CLU;PQBP1;GRIN2A;GNG3;KCNT1;TUBA1A;UBB;GRM7;PPP2R1A;APOE;SCN1A;COX8A;BRD2;GABBR2;MAP2K1;CIZ1;SNRPN;ANGPT1;MDH2;TSC2;NAGA;ANK1;DDOST;ALDH4A1;TGFBR3;TUBB2B;TUBB2A;SST;SCN8A;CCDC88C;GNB1;TBC1D24;GNAS;AMPH;BCOR;SCN2A;NAT8L;HCN1 |
| Autistic Disorder | 32/677 | 6.15E-08 | 5.49E-05 | 3.24 | 53.88 | YWHAE;GABRB2;RYR2;NRP2;MAGED1;ANKRD11;CACNA1A;CACNA1G;GRIN2A;DNER;PDE4A;MBP;APOE;MACROD2;DLGAP2;SCN1A;GABBR2;SNRPN;CADM1;TMEM132B;TSC2;NAGA;ATP2B2;MIF;YWHAZ;VEGFA;FOXP1;SST;SCN8A;ARHGEF2;SCN2A;SHANK2 |
| Undifferentiated carcinoma | 18/237 | 6.41E-08 | 5.49E-05 | 5.24 | 86.85 | RPL3;CADM1;MDH2;PGAM1;PEBP1;MIF;YWHAZ;VEGFA;CNN3;PRDX2;VTN;TUBB2B;PKM;PPP2R1A;ID4;NDN;SGK1;RAN |
| Animal Mammary Neoplasms | 14/145 | 1.01E-07 | 6.17E-05 | 6.76 | 108.90 | RPL3;MDH2;PGAM1;PEBP1;MIF;YWHAZ;CNN3;PRDX2;TUBB2B;PKM;ID4;NDN;SGK1;RAN |
| Carcinomatosis | 15/176 | 1.84E-07 | 9.84E-05 | 5.90 | 91.55 | RPL3;MDH2;PGAM1;PEBP1;MIF;YWHAZ;CNN3;VEGFA;PRDX2;TUBB2B;PKM;ID4;NDN;SGK1;RAN |
| Anaplastic carcinoma | 16/204 | 2.25E-07 | 0.000107 | 5.40 | 82.70 | RPL3;MDH2;PGAM1;PEBP1;MIF;YWHAZ;VEGFA;CNN3;PRDX2;TUBB2B;PKM;ID4;NDN;SGK1;TMSB10;RAN |
| Bipolar Disorder | 35/837 | 2.8E-07 | 0.00012 | 2.86 | 43.15 | YWHAE;GABRB2;PFKFB3;MCTP1;TENM4;ITPR1;CACNA1A;ATP1A3;FBLN1;ADD3;SYNE1;PTPA;VTN;GRIN2A;GRM7;UBC;PDE4A;APOE;MACROD2;TMSB10;YWHAH;USP46;TRPC3;PGAM1;RARRES2;CSNK1D;SYP;YWHAZ;VEGFA;SST;SCN8A;NOS1AP;KCTD12;CALM2;DGKH |
| Impaired cognition | 30/671 | 5.3E-07 | 0.000206 | 3.04 | 43.94 | YWHAE;RYR2;RTN3;SPON1;SATB2;ANKRD11;ITPR1;CACNA1A;ADD3;CLU;CX3CL1;CACNA1G;KCNT1;PDE4A;APOE;SCN1A;SNCA;TSC2;NAGA;SYP;YWHAZ;VEGFA;FOXP1;MAG;SST;SCN8A;GNAS;AMPH;SCN2A;SHANK2 |
| Alzheimer's Disease | 61/1982 | 5.93E-07 | 0.000211 | 2.15 | 30.86 | RYR2;SPON1;PEBP1;SLC7A11;CLU;CX3CL1;SYNE1;RIMS2;TUBA1B;TIMP3;PDE4A;COX8A;MAP2K1;APLP1;TSC2;CSNK1D;MIF;SYP;ANK1;YWHAZ;EID1;OLFM1;ARC;FRMD6;DDAH1;UQCRC1;GEMIN4;AMPH;PPIB;SHANK2;RTN3;PFKFB3;ITPR1;ADD3;CACNA1G;PTPA;CST3;PRDX2;GRIN2A;MOBP;UBB;RPL13;MBP;APOE;VAT1L;SNCA;ABCA2;CIZ1;TIPARP;VEGFA;PPP1CA;PPP1R1A;NACA;SST;CPE;CALM3;FXYD6;PRPF31;CALM2;RAN;HCN1 |
| Cardiac Arrest | 15/201 | 1.02E-06 | 0.000334 | 5.10 | 70.43 | RYR2;ITPR1;CACNA1A;DYNLL1;YWHAZ;CACNA1G;NOS1AP;PKP2;TIMP3;PDE4A;CALM3;APOE;SCN2A;CALM2;RAN |
| Seizures, Focal | 12/129 | 1.24E-06 | 0.00038 | 6.45 | 87.72 | GRIN2A;NRP2;KCNT1;SST;SCN8A;TBC1D24;TSC2;APOE;SCN2A;CLU;SCN1A;HCN1 |
| Depressive disorder | 31/741 | 1.39E-06 | 0.000396 | 2.84 | 38.25 | GPM6A;GPR26;MAGED1;COX4I1;ATP1A3;CLU;CACNA1G;SYNE1;GRIN2A;KCNT1;GRM7;PDE4A;APOE;SNCA;UNC13A;CADM1;SPRY4;MIF;SYP;ANK1;YWHAZ;VEGFA;TGFBR3;GNB1;NOS1AP;GEMIN4;GNAS;CALM3;NAT8L;CALM2;HCN1 |

| | | | | | | |
|---|---|---|---|---|---|---|
| Parkinson Disease | 39/1064 | 1.55E-06 | 0.000416 | 2.50 | 33.42 | RAB3B;PEBP1;CACNA1A;ATP1A3;CLU;CX3CL1;PQBP1;SYNE1;CST3;PRDX2;GRIN2A;PRDX5;TUBA1A;UBB;MCF2L;UBC;NDN;MBP;APOE;GPC4;SCN1A;SKP1;YWHAH;SNCA;TSC2;CSNK1D;RPL23A;SYP;YWHAZ;VEGFA;MAG;GSTA4;PSMA2;SST;SCN8A;CNTN1;SCN2A;SGK1;RAN |
| Mental Retardation | 41/1158 | 1.96E-06 | 0.000494 | 2.42 | 31.75 | KCNK9;TUSC3;SATB2;ANKRD11;USP11;ITPR1;ATP1A3;PQBP1;RIMS2;GRIN2A;TUBA1A;GRM7;PPP2R1A;GPC3;APOE;SCN1A;COX8A;BRD2;GABBR2;MAP2K1;SNRPN;CADM1;TSC2;NAGA;SYP;FANCE;ANK1;FOXP1;ALDH4A1;TUBB2B;SCN8A;CCDC88C;GNB1;TBC1D24;GNAS;PRPF31;BCOR;SCN2A;CDK13;SHANK2;HCN1 |
| Mammary Neoplasms, Experimental | 13/160 | 2.09E-06 | 0.000497 | 5.57 | 72.87 | RPL3;MDH2;PGAM1;PEBP1;MIF;YWHAZ;CNN3;PRDX2;TUBB2B;PKM;ID4;SGK1;RAN |
| Carcinoma, Spindle-Cell | 14/188 | 2.39E-06 | 0.000531 | 5.08 | 65.73 | RPL3;MDH2;PGAM1;PEBP1;MIF;YWHAZ;CNN3;PRDX2;TUBB2B;PKM;ID4;NDN;SGK1;RAN |
| Neuroblastoma | 53/1698 | 2.48E-06 | 0.000531 | 2.15 | 27.77 | VIPR1;RRP1;PEBP1;CLU;CX3CL1;RIMS2;RBM3;CTSL;DNER;TMSB10;SCN1A;SKP1;COX8A;MAP2K1;SNRPN;PLEKHG5;APLP1;MIF;SYP;DYNLL1;YWHAZ;FOXP1;TGFBR3;ALDOC;ARHGEF2;SGK1;PTMA;DSC3;NOTCH2;DOT1L;SRSF1;ITPR1;PTPA;PRDX2;SDCBP;VTN;PRDX5;UBB;NNAT;GPC3;NDN;APOE;SNCA;ABCA2;CADM1;TIPARP;PTPN14;VEGFA;SST;GNAS;CALM3;CALM2;RAN |
| Epilepsy, Temporal Lobe | 13/181 | 8.14E-06 | 0.001658 | 4.87 | 57.07 | GABRB2;GABBR2;CACNA1A;VEGFA;GRIN2A;SST;MBP;CALM3;APOE;SCN2A;CALM2;SCN1A;HCN1 |
| Genetic Diseases, Inborn | 23/507 | 9.3E-06 | 0.001809 | 3.04 | 35.22 | RYR2;GABBR2;NRP2;MAP2K1;SATB2;ANKRD11;ITPR1;TSC2;CACNA1A;ATP1A3;PQBP1;FOXP1;GRIN2A;COL1A2;KCNT1;TUBA1A;SCN8A;GNB1;TBC1D24;GNAS;SCN2A;CDK13;SCN1A |
| Amyotrophic Lateral Sclerosis | 27/660 | 1.03E-05 | 0.00191 | 2.74 | 31.52 | SLC7A11;ADARB2;PTPA;CST3;RIMS2;GRIN2A;MOBP;SUMO2;TIMP3;APOE;KIFAP3;SNCA;COX8A;UNC13A;SNRPN;MDH1;TIPARP;OSBPL3;SYP;RANGAP1;VEGFA;SST;SCN8A;GEMIN4;AMPH;SGK1;RAN |
| Astrocytoma | 29/741 | 1.11E-05 | 0.001975 | 2.63 | 29.98 | NOTCH2;PFKFB3;PEBP1;CLU;RPS15;CST3;RIMS2;VTN;PRDX5;MBP;APOE;SNCA;COX8A;BRD2;TLE2;MAP2K1;SNRPN;CD93;ANGPT1;TSC2;YWHAZ;VEGFA;PPP1R1A;SST;SUB1;ID4;CALM3;CALM2;MATN2 |
| Carcinoma | 14/216 | 1.19E-05 | 0.002016 | 4.37 | 49.53 | RPL3;MDH2;PGAM1;PEBP1;MIF;YWHAZ;CNN3;PRDX2;TUBB2B;PKM;ID4;NDN;SGK1;RAN |
| Neurodegenerative Disorders | 29/745 | 1.22E-05 | 0.002016 | 2.61 | 29.54 | ATP6;ITPR1;CACNA1A;CLU;CX3CL1;PQBP1;PTPA;CST3;RIMS2;UBB;UBC;TIMP3;PDE4A;MBP;APOE;SNCA;UNC13A;SNRPN;TIPARP;APLP1;MIF;YWHAZ;VEGFA;SST;SCN8A;GEMIN4;CALM3;SGK1;CALM2 |
| Osteoarthrosis Deformans | 9/89 | 1.37E-05 | 0.002169 | 7.02 | 78.63 | SDCBP;SEC23A;COL1A2;PPP2R1A;PSMB1;ATP1A3;CLU;RAN;GLS |
| Learning Disabilities | 8/72 | 2.07E-05 | 0.003083 | 7.78 | 83.94 | GPM6A;TSC2;CALM3;SYP;RANGAP1;CALM2;CACNA1G;VEGFA |
| Global developmental delay | 37/1102 | 2.09E-05 | 0.003083 | 2.26 | 24.36 | YWHAE;RYR2;SATB2;ANKRD11;ATP6;ITPR1;CACNA1A;ATP1A3;ADD3;PQBP1;PPP1CB;GRIN2A;PPP2R1A;NDN;MBP;SCN1A;MAP2K1;SNRPN;MDH2;TSC2;NAGA;MIF;FANCE;ANK1;DDOST;FOXP1;ALDH6A1;TUBB2B;MAG;TUBB2A;SUB1;SCN8A;GNB1;TBC1D24;SCN2A;CDK13;NAT8L |
| Mental and motor retardation | 35/1021 | 2.31E-05 | 0.003225 | 2.30 | 24.59 | YWHAE;SATB2;ANKRD11;ATP6;ITPR1;CACNA1A;ATP1A3;ADD3;PQBP1;TMEM47;PPP1CB;GRM7;PPP2R1A;NDN;SCN1A;MAP2K1;CIZ1;SNRPN;MDH2;TSC2;NAGA;FANCE;ANK1;DDOST;ALDH6A1;TUBB2B;MAG;TUBB2A;SCN8A;GNB1;TBC1D24;GEMIN4;SCN2A;CDK13;NAT8L |
| Mental Depression | 24/575 | 2.34E-05 | 0.003225 | 2.79 | 29.71 | GPM6A;GPR26;PFKFB3;MAGED1;COX4I1;ATP1A3;SYP;ANK1;CLU;YWHAZ;VEGFA;SYNE1;TGFBR3;GRIN2A;GRM7;NOS1AP;GEMIN4;PDE4A;CALM3;APOE;NAT8L;CALM2;SNCA;HCN1 |
| Central neuroblastoma | 49/1655 | 2.56E-05 | 0.003419 | 2.01 | 21.28 | RRP1;PEBP1;CLU;CX3CL1;RIMS2;RBM3;CTSL;DNER;TMSB10;SCN1A;SKP1;COX8A;MAP2K1;SNRPN;PLEKHG5;APLP1;MIF;SYP;DYNLL1;YWHAZ;FOXP1;TGFBR3;ALDOC;ARHGEF2;SGK1;PTMA;DSC3;NOTCH2;DOT1L;SRSF1;ITPR1;PTPA;PRDX2;VTN;PRDX5;UBB;GPC3;NDN;APOE;SNCA;ABCA2;CADM1;TIPARP;VEGFA;SST;GNAS;CALM3;CALM2;RAN |
| Medulloblastoma | 24/590 | 3.53E-05 | 0.004359 | 2.71 | 27.79 | NOTCH2;BRD2;NRP2;MAP2K1;CIZ1;CADM1;TSC2;SYP;DYNLL1;VEGFA;LDHB;COL1A2;PKM;SST;SUB1;SCN8A;DNER;NNAT;GPC3;GNAS;TIMP3;CALM3;CALM2;SNCA |
| Ataxia | 13/208 | 3.56E-05 | 0.004359 | 4.19 | 42.91 | COX8A;ATP6;ITPR1;CACNA1A;ATP1A3;ANK1;SYNE1;GRIN2A;SCN8A;CALM3;CALM2;SCN1A;HCN1 |
| Neurodevelopmental Disorders | 13/208 | 3.56E-05 | 0.004359 | 4.19 | 42.91 | GABBR2;SNRPN;SATB2;ANKRD11;ITPR1;TSC2;CACNA1A;ATP1A3;FOXP1;GRIN2A;SCN8A;SCN2A;SHANK2 |
| Pituitary Diseases | 14/240 | 3.85E-05 | 0.004572 | 3.90 | 39.64 | BRD2;VIPR1;NRP2;MIF;CLU;YWHAZ;VEGFA;RIMS2;PRDX2;UBC;GNAS;CPE;CALM3;CALM2 |

| | | | | | | |
|---|---|---|---|---|---|---|
| Spinocerebellar Ataxia Type 1 | 8/79 | 4.09E-05 | 0.004732 | 7.01 | 70.86 | RIMS2;PTPA;TRPC3;CCDC88C;CACNA1A;PQBP1;CACNA1G;VEGFA |
| Drug Resistant Epilepsy | 7/59 | 4.46E-05 | 0.005022 | 8.36 | 83.75 | GRIN2A;KCNT1;APOE;SGK1;SCN2A;SCN1A;CNN3 |
| Absence Epilepsy | 7/60 | 4.98E-05 | 0.005463 | 8.20 | 81.26 | KCNK9;SCN8A;CACNA1A;YWHAZ;SCN1A;CACNA1G;HCN1 |
| Aura | 7/61 | 5.55E-05 | 0.00579 | 8.05 | 78.89 | GRIN2A;SCN8A;SELENOW;TSC2;SCN2A;SCN1A;HCN1 |
| Awakening Epilepsy | 7/61 | 5.55E-05 | 0.00579 | 8.05 | 78.89 | GRIN2A;SCN8A;SELENOW;TSC2;SCN2A;SCN1A;HCN1 |
| Epilepsy, Cryptogenic | 7/63 | 6.84E-05 | 0.006973 | 7.76 | 74.43 | GRIN2A;SCN8A;SELENOW;TSC2;SCN2A;SCN1A;HCN1 |
| Sarcoma | 25/658 | 7.31E-05 | 0.007233 | 2.52 | 24.04 | YWHAE;VIPR1;COX4I1;SRSF1;CLU;CX3CL1;PTPA;RIMS2;GPC3;MAP2K1;CADM1;ANGPT1;TIPARP;TSC2;DYNLL1;ZWINT;VEGFA;FOXP1;TGFBR3;SUB1;SCN8A;CPE;CALM3;BCOR;CALM2 |
| Borderline Personality Disorder | 9/110 | 7.44E-05 | 0.007233 | 5.56 | 52.81 | GABRB2;TENM4;ANGPT1;SST;SCN8A;GNAS;SCN4B;DGKH;VEGFA |
| Hemiplegia, Crossed | 3/6 | 8.05E-05 | 0.007443 | 61.48 | 579.59 | TBC1D24;ATP1A3;CACNA1A |
| HYPOCALCIURIC HYPERCALCEMIA, FAMILIAL, TYPE III | 3/6 | 8.05E-05 | 0.007443 | 61.48 | 579.59 | AP2S1;CALM3;CALM2 |
| Epileptic encephalopathy | 20/468 | 8.17E-05 | 0.007443 | 2.83 | 26.67 | GABRB2;GABBR2;MAP2K1;CIZ1;MDH2;TSC2;CACNA1A;ATP1A3;NAGA;GRIN2A;TUBB2B;KCNT1;TUBA1A;SCN8A;CCDC88C;GNB1;TBC1D24;SCN2A;SCN1A;HCN1 |
| Prostate carcinoma | 77/3145 | 8.57E-05 | 0.007638 | 1.69 | 15.87 | RAB3B;VIPR1;ARF1;NRP2;TUSC3;PEBP1;CLU;CX3CL1;RIMS2;PPP1CB;RBM3;THBD;TUBA1B;GRM7;CTSL;DNER;SUMO2;SLC39A6;TIMP3;PDE4A;YWHAH;COX8A;BRD2;MAP2K1;SNRPN;TRPC3;TSC2;MIF;SYP;DYNLL1;YWHAZ;FOXP1;TGFBR3;PKM;SUB1;GEMIN4;ARHGEF2;SGK1;PTMA;PPP1R12C;DSC3;SEC23A;ARHGEF26;MAGED1;ATP6;ITPR1;FBLN1;PTPA;CST3;LDHB;PRDX2;VTN;PRDX5;PSMB1;RPL13;MBP;APOE;PCDHA7;GPR158;SNCA;ABCA2;ANGPT1;TIPARP;MDH2;CRIM1;ZWINT;VEGFA;NR6A1;TSPAN13;PPP1R1A;SST;ID4;GNAS;CNTN1;CALM3;CALM2;RAN |
| Romano-Ward Syndrome | 5/29 | 9.2E-05 | 0.008034 | 12.88 | 119.66 | NOS1AP;PKP2;CALM3;CALM2;SCN4B |
| Vascular Diseases | 18/401 | 0.0001 | 0.008543 | 2.97 | 27.37 | NOTCH2;ANGPT1;RARRES2;NDNF;CLU;YWHAZ;CX3CL1;DDOST;VEGFA;CST3;VTN;THBD;CTSL;ADGRB1;TIMP3;PDE4A;APOE;SGK1 |
| West Syndrome | 7/67 | 0.000102 | 0.008543 | 7.24 | 66.58 | RIMS2;KCNT1;SCN8A;TBC1D24;TSC2;SCN2A;SCN1A |
| Paroxysmal familial ventricular fibrillation | 4/16 | 0.000104 | 0.00858 | 20.55 | 188.41 | RYR2;PKP2;CALM3;CALM2 |
| Epilepsies, Partial | 6/48 | 0.000117 | 0.00927 | 8.85 | 80.08 | GRIN2A;KCNT1;TBC1D24;TSC2;SCN2A;SCN1A |
| Intellectual Disability | 64/2503 | 0.000118 | 0.00927 | 1.75 | 15.80 | KCNK9;TUSC3;LITAF;PQBP1;SYNE1;RIMS2;PPP1CB;KCNT1;TUBA1A;GRM7;PPP2R1A;SPEG;DLGAP2;SCN1A;COX8A;BRD2;MAP2K1;UNC13A;SNRPN;TSC2;SYP;ANK1;DDOST;FOXP1;TUBB2B;TUBB2A;SCN8A;TBC1D24;GEMIN4;ARHGEF2;SHANK2;DGKH;NOTCH2;MAGED1;SATB2;ANKRD11;USP11;ATP6;ITPR1;CACNA1A;ATP1A3;CACNA1G;PTPA;GRIN2A;GPC3;NDN;APOE;SNCA;GABBR2;CADM1;MDH2;NAGA;FANCE;ALDH4A1;COL1A2;CCDC88C;GNB1;PLXNB3;GNAS;PRPF31;BCOR;SCN2A;CDK13;HCN1 |
| Adenocarcinoma | 48/1712 | 0.000119 | 0.00927 | 1.89 | 17.07 | NOTCH2;VIPR1;NRP2;PFKFB3;RTN1;SATB2;ATP6;PEBP1;CLU;SYNE1;PTPA;LDHB;PRDX2;SDCBP;VTN;CTSL;UBC;NNAT;GPC3;TIMP3;APOE;TMSB10;SCN1A;SNCA;COX8A;KLF10;MAP2K1;CADM1;ANGPT1;PGAM1;TSC2;MIF;SYP;DYNLL1;VEGFA;FOXP1;TGFBR3;PKM;SST;SUB1;ID4;GNAS;CNTN1;CALM3;ARHGEF2;CALM2;PTMA;DSC3 |
| Myocardial Ischemia | 19/448 | 0.000136 | 0.010362 | 2.80 | 24.97 | COX8A;RYR2;PFKFB3;RTN1;ANGPT1;TIPARP;PEBP1;MIF;COX5B;CX3CL1;VEGFA;ARC;ALDH6A1;TUBA1A;PSMB5;ENSA;CALM3;APOE;CALM2 |
| Cerebral Ischemia | 14/275 | 0.000164 | 0.012329 | 3.37 | 29.37 | MAP2K1;ANGPT1;RPL13A;DYNLL1;ADD3;YWHAZ;CX3CL1;VEGFA;TIMP3;CALM3;MBP;APOE;SGK1;CALM2 |
| Infantile Severe Myoclonic Epilepsy | 5/33 | 0.000174 | 0.01275 | 11.03 | 95.49 | SCN8A;CACNA1A;SCN2A;CACNA1G;SCN1A |

| | | | | | | |
|---|---|---|---|---|---|---|
| Breast Carcinoma | 109/4963 | 0.000178 | 0.01275 | 1.56 | 13.43 | VIPR1;RPL3;PEBP1;RPL8;CLU;GLS;RIMS2;SYNPR;PPP2R1A;SLC39A6;MAP2K1;SNRPN;TSC2;CSNK1D;MIF;FOXP1;SUB1;HECW1;GEMIN4;AMPH;PPIB;DSC3;NOTCH2;PFKFB3;RTN1;ANKRD11;ITPR1;FBLN1;RPAP1;CACNA1G;LDHB;PRDX2;SDCBP;VTN;PRDX5;UBC;G3BP2;NDN;APOE;BANP;PRPF38B;CADM1;TIPARP;NSG1;FANCE;ZWINT;MLLT11;PPP1R1A;CCDC88C;GNB1;ID4;GNAS;CALM3;CALM2;YWHAE;ARF1;NRP2;KCNK9;COX4I1;SLC7A11;LITAF;CX3CL1;SYNE1;RBM3;TUBA1B;CTSL;MCF2L;DNER;SUMO2;TIMP3;ELOB;COX8A;KLF10;BRD2;RPL13A;DYNLL1;YWHAZ;TGFBR3;PKM;NOS1AP;ARHGEF2;SGK1;PTMA;PPP1R12C;AHNAK;DOT1L;SRSF1;ADAM22;PDXP;PTPA;CST3;GPC3;RPL13;SNCA;ANGPT1;USP22;ATP2B2;RPL23A;PTPN14;VEGFA;PPP1CA;NR6A1;TMEM158;TSPAN13;PSMC3;SST;PRPF31;RAN;HCN1 |
| Glioblastoma | 52/1937 | 0.000179 | 0.01275 | 1.81 | 15.63 | NRP2;GPR26;KCNK9;TUSC3;SLC7A11;ADARB2;CLU;GLS;RPS15;ADAMTS4;CTSL;DNER;TIMP3;DLGAP3;COX8A;BRD2;MAP2K1;CD93;MIF;SYP;FOXP1;PKM;SUB1;ADGRB1;ARHGEF2;PLPP3;SGK1;PPIB;PPP1R12C;NOTCH2;PFKFB3;ARHGEF26;PTPA;CST3;PRDX2;VTN;NDN;MBP;APOE;MDH1;ANGPT1;TIPARP;RARRES2;VEGFA;PPP1CA;PPP1R1A;SST;ID4;CPE;CALM3;BCOR;CALM2 |
| Infant, Premature | 14/278 | 0.000184 | 0.012892 | 3.33 | 28.66 | COX8A;NOTCH2;SNRPN;ANGPT1;VEGFA;MLLT11;RIMS2;COL1A2;SUB1;GNAS;TIMP3;CALM3;CCNL1;CALM2 |
| Carcinoid Tumor | 12/213 | 0.000189 | 0.013057 | 3.74 | 32.05 | SDCBP;MAP2K1;SST;SUB1;APLP1;GNAS;TIMP3;CPE;SYP;BCOR;CLU;VEGFA |
| Mental deterioration | 6/53 | 0.000205 | 0.013918 | 7.91 | 67.12 | TBC1D24;GNAS;ATP1A3;APOE;SCN1A;SNCA |
| Shy-Drager Syndrome | 7/75 | 0.000208 | 0.013918 | 6.39 | 54.15 | CST3;PRDX5;UBB;PDXP;MBP;APOE;SNCA |
| Alternating hemiplegia of childhood | 3/8 | 0.00022 | 0.01449 | 36.89 | 310.63 | TBC1D24;ATP1A3;CACNA1A |
| Cognitive delay | 31/966 | 0.000225 | 0.014619 | 2.13 | 17.87 | YWHAE;SATB2;ANKRD11;ATP6;ITPR1;CACNA1A;ATP1A3;ADD3;PQBP1;PPP1CB;PPP2R1A;NDN;SCN1A;MAP2K1;SNRPN;MDH2;TSC2;NAGA;FANCE;ANK1;DDOST;ALDH6A1;TUBB2B;MAG;TUBB2A;SCN8A;GNB1;TBC1D24;SCN2A;CDK13;NAT8L |
| Huntington Disease | 23/627 | 0.000236 | 0.015084 | 2.42 | 20.22 | ITPR1;CACNA1A;NSG1;VEGFA;FOXP1;RIMS2;GRIN2A;UBB;SST;UBC;SUMO2;NNAT;CNTN1;PDE4A;CALM3;MBP;APOE;SGK1;CALM2;PTMA;RAN;SKP1;SNCA |
| Hypoplasia of corpus callosum | 11/187 | 0.00024 | 0.015113 | 3.91 | 32.56 | YWHAE;TMEM47;ALDH6A1;TUBB2B;TUBA1A;KCNT1;TUBB2A;GRM7;PPP2R1A;MDH2;ANKRD11 |
| Epilepsy, Generalized | 6/55 | 0.000252 | 0.015427 | 7.58 | 62.81 | KCNT1;CACNA1A;SCN2A;SCN1A;CACNA1G;HCN1 |
| Brugada Syndrome (disorder) | 6/55 | 0.000252 | 0.015427 | 7.58 | 62.81 | KCNT1;NOS1AP;PKP2;CALM3;SCN4B;CALM2 |
| Status Epilepticus | 12/221 | 0.000266 | 0.016015 | 3.59 | 29.59 | GRIN2A;KCNT1;SST;SUB1;SCN8A;TSC2;ATP1A3;SCN2A;CX3CL1;SCN1A;VEGFA;HCN1 |
| Degenerative polyarthritis | 31/976 | 0.000269 | 0.016016 | 2.10 | 17.30 | SPON1;SEC23A;DOT1L;ATP1A3;CLU;GLS;ADAMTS4;SDCBP;VTN;PRDX5;PPP2R1A;CTSL;MCF2L;PSMB1;SUMO2;TIMP3;PDE4A;APOE;MAP2K1;SNRPN;ANGPT1;RARRES2;ANK1;YWHAZ;VEGFA;FOXP1;TGFBR3;COL1A2;DDAH1;SCN8A;RAN |
| Vertigo | 6/56 | 0.000279 | 0.016125 | 7.43 | 60.81 | RYR2;CACNA1A;NAGA;CALM3;SCN2A;CALM2 |
| Channelopathies | 6/56 | 0.000279 | 0.016125 | 7.43 | 60.81 | RYR2;ATP6;PKP2;CACNA1A;SCN1A;HCN1 |
| Tonic Seizures | 8/104 | 0.000285 | 0.016271 | 5.18 | 42.28 | NRP2;SST;TSC2;APOE;SCN2A;CLU;SCN1A;HCN1 |
| Pick Disease of the Brain | 8/105 | 0.000304 | 0.017141 | 5.13 | 41.51 | RIMS2;PRDX2;FMNL1;UNC13A;UBB;APOE;RAN;SNCA |
| Prolonged QT interval | 4/21 | 0.000322 | 0.017448 | 14.50 | 116.62 | GPC3;GPC4;CALM2;SCN4B |
| Dementia, familial Danish | 3/9 | 0.000326 | 0.017448 | 30.74 | 246.75 | CST3;SYP;CLU |
| EPILEPTIC ENCEPHALOPATHY, EARLY INFANTILE, 1 | 3/9 | 0.000326 | 0.017448 | 30.74 | 246.75 | TBC1D24;SCN2A;HCN1 |
| Hemiplegic migraine | 3/9 | 0.000326 | 0.017448 | 30.74 | 246.75 | ATP1A3;CACNA1A;SCN1A |
| Dystonia Disorders | 9/135 | 0.000352 | 0.01839 | 4.45 | 35.36 | RIMS2;YWHAE;CIZ1;SCN8A;MCF2L;GNB1;CACNA1A;ATP1A3;APOE |
| Motor Neuron Disease | 9/135 | 0.000352 | 0.01839 | 4.45 | 35.36 | RIMS2;COX8A;SNRPN;PLEKHG5;APOE;RAN;PQBP1;SNCA;VEGFA |

| | | | | | | |
|---|---|---|---|---|---|---|
| Malignant neoplasm of prostate | 76/3239 | 0.000372 | 0.019166 | 1.61 | 12.69 | RAB3B;VIPR1;ARF1;NRP2;ARL6IP1;TUSC3;PEBP1;CLU;CX3CL1;RIMS2;PPP1CB;RBM3;THBD;TUBA1B;CTSL;DNER;SUMO2;SLC39A6;TIMP3;PDE4A;YWHAH;COX8A;BRD2;MAP2K1;SNRPN;TRPC3;TSC2;MIF;SYP;DYNLL1;YWHAZ;FOXP1;TGFBR3;OLFM1;PKM;SUB1;GEMIN4;ARHGEF2;SGK1;PTMA;PPP1R12C;DSC3;ARHGEF26;ATP6;ITPR1;FBLN1;CNN3;PTPA;CST3;LDHB;PRDX2;VTN;PRDX5;RPL13;MBP;APOE;GPR158;SNCA;CIZ1;CADM1;ANGPT1;TIPARP;MDH2;CRIM1;ZWINT;VEGFA;NR6A1;TSPAN13;PPP1R1A;SST;ID4;GNAS;CNTN1;CALM3;CALM2;RAN |
| Severe depression | 4/22 | 0.000388 | 0.019525 | 13.69 | 107.56 | YWHAE;NOS1AP;APOE;CALM2 |
| Abnormal behavior | 16/372 | 0.000389 | 0.019525 | 2.83 | 22.21 | GPM6A;PLPPR4;RRP1;TSC2;LDLRAD4;PQBP1;GRIN2A;SCN8A;NOS1AP;NDN;APOE;SCN2A;DLGAP3;SHANK2;SCN1A;HCN1 |
| Convulsive Seizures | 8/109 | 0.000392 | 0.019525 | 4.92 | 38.61 | NRP2;SST;SCN8A;TSC2;APOE;CLU;SCN1A;HCN1 |
| Ventricular arrhythmia | 6/60 | 0.000407 | 0.020037 | 6.88 | 53.69 | RYR2;NOS1AP;PKP2;CALM3;SGK1;CALM2 |
| Autism Spectrum Disorders | 21/572 | 0.000434 | 0.020988 | 2.41 | 18.69 | RYR2;CADM1;TENM4;SATB2;ANKRD11;ITPR1;TSC2;CACNA1A;ATP2B2;MIF;FOXP1;CACNA1G;SYNE1;GRIN2A;GRM7;SUB1;MACROD2;SCN2A;SHANK2;SCN1A;HCN1 |
| Movement Disorders | 10/169 | 0.000436 | 0.020988 | 3.92 | 30.34 | YWHAE;GABBR2;ARC;UNC13A;SCN8A;ITPR1;CACNA1A;ATP1A3;SCN1A;SNCA |
| Hemiparesis | 6/61 | 0.000446 | 0.021194 | 6.75 | 52.10 | TGFBR3;TUBB2B;TSC2;ATP1A3;CACNA1A;SCN1A |
| Hereditary X-Linked Recessive Spastic Paraplegia | 3/10 | 0.00046 | 0.021652 | 26.34 | 202.41 | PRDX5;PDXP;PQBP1 |
| Electroencephalogram abnormal | 9/141 | 0.000485 | 0.022553 | 4.24 | 32.39 | YWHAE;GABBR2;MAP2K1;SNRPN;GNB1;TSC2;CACNA1A;SCN2A;SCN1A |
| Amyloidosis | 27/833 | 0.000494 | 0.022736 | 2.14 | 16.26 | SPON1;ITPR1;ADD3;CLU;CST3;RIMS2;VTN;PRDX5;UBB;UBC;MBP;APOE;SNCA;ABCA2;MAP2K1;APLP1;SYP;YWHAZ;VEGFA;PPP1CA;OLFM1;ARC;MAG;DDAH1;SST;CPE;PPIB |
| Paresis | 8/113 | 0.0005 | 0.022762 | 4.73 | 35.98 | RIMS2;RYR2;SNRPN;PKM;SCN8A;ATP6;CACNA1A;SYNE1 |
| Tonic - clonic seizures | 10/173 | 0.000525 | 0.023632 | 3.82 | 28.89 | NRP2;SST;SCN8A;GNB1;TSC2;APOE;SCN2A;CLU;SCN1A;HCN1 |
| Intracranial Aneurysm | 8/114 | 0.00053 | 0.02364 | 4.69 | 35.37 | TGFBR3;CST3;COL1A2;ANGPT1;TMEM132B;TIMP3;APOE;VEGFA |
| Visual seizure | 11/207 | 0.000569 | 0.025116 | 3.50 | 26.18 | VTN;ARC;NRP2;YWHAB;SST;TSC2;PDE4A;APOE;CLU;YWHAZ;HCN1 |
| Complex partial seizures | 8/116 | 0.000595 | 0.025778 | 4.60 | 34.17 | NRP2;SST;GNB1;TSC2;APOE;CLU;SCN1A;HCN1 |
| Cerebral Palsy | 7/89 | 0.000596 | 0.025778 | 5.29 | 39.30 | THBD;TUBB2B;TUBA1A;CTSL;SCN8A;APOE;ADD3 |
| Myoclonic Epilepsy | 5/43 | 0.00062 | 0.026243 | 8.13 | 60.02 | BRD2;SCN8A;CACNA1A;SCN2A;SCN1A |
| Abnormality of brainstem morphology | 3/11 | 0.000625 | 0.026243 | 23.05 | 170.04 | TGFBR3;NAGA;SCN1A |
| Neurodevelopmental delay | 3/11 | 0.000625 | 0.026243 | 23.05 | 170.04 | SCN2A;DDOST;SCN1A |
| Poor school performance | 30/985 | 0.000674 | 0.027987 | 2.01 | 14.66 | KCNK9;TUSC3;SATB2;ANKRD11;ITPR1;CACNA1A;ATP1A3;PQBP1;GRM7;PPP2R1A;SCN1A;GABBR2;MAP2K1;SNRPN;TSC2;NAGA;SYP;FANCE;ANK1;FOXP1;ALDH4A1;TUBB2B;SCN8A;CCDC88C;GNB1;GNAS;PRPF31;BCOR;CDK13;HCN1 |
| Pulmonary Stenosis | 6/66 | 0.000682 | 0.028057 | 6.19 | 45.12 | PPP1CB;NOTCH2;MAP2K1;GPC3;GPC4;BCOR |
| Ataxia, Truncal | 5/44 | 0.00069 | 0.028132 | 7.92 | 57.63 | SCN8A;ATP1A3;CACNA1A;NAT8L;HCN1 |
| Glioblastoma Multiforme | 27/854 | 0.00072 | 0.029063 | 2.08 | 15.05 | NOTCH2;NRP2;TUSC3;ADARB2;CLU;GLS;SYNE1;VTN;CTSL;DNER;DLGAP3;TMSB10;COX8A;BRD2;ANGPT1;TIPARP;MIF;VEGFA;PPP1CA;PKM;PPP1R1A;SUB1;ADGRB1;CALM3;SGK1;PPIB;CALM2 |
| Glioma | 55/2211 | 0.000752 | 0.03009 | 1.67 | 12.00 | NRP2;GPR26;PEBP1;SLC7A11;CLU;CX3CL1;GLS;RPS15;ADAMTS4;CTSL;DNER;DACT2;TIMP3;DLGAP3;COX8A;BRD2;MAP2K1;CD93;PGAM1;TSC2;RPL13A;MIF;RANGAP1;FOXP1;MAG;PKM;SUB1;SCN8A;ADGRB1;PKP2;ARHGEF2;SGK1;DSC3;NOTCH2;ARHGEF26;MAGED1;ADAM22;ADD3;PTPA;CST3;PRDX2;VTN;MBP;CADM1;ANGPT1;TIPARP;RARRES2;VEGFA;SST;ID4;PLXNB3;CNTN1;CPE;CALM3;CALM2 |
| Heart failure | 26/815 | 0.000791 | 0.031326 | 2.10 | 14.97 | RYR2;ITPR1;PEBP1;ATP1A3;CX3CL1;PTPA;CST3;PPP2R1A;DNER;SUMO2;TIMP3;APOE;TRPC3;ANGPT1;MDH2;DYNLL1;VEGFA;ARC;COL1A2;PPP1R1A;SCN8A;NOS1AP;PKP2;CALM3;SGK1;CALM2 |
| Unipolar Depression | 19/517 | 0.000798 | 0.031326 | 2.41 | 17.17 | YWHAE;GPM6A;GABBR2;MAP2K1;PFKFB3;CADM1;TENM4;RARRES2;PEBP1;MIF;VEGFA;SYNE1;GRM7;SST;GNB1;PDE4A;APOE;SGK1;DGKH |
| Childhood Absence Epilepsy | 3/12 | 0.000824 | 0.032059 | 20.49 | 145.49 | KCNK9;CACNA1A;CACNA1G |
| Spinocerebellar Ataxia Type 5 | 5/46 | 0.000849 | 0.032717 | 7.53 | 53.25 | TRPC3;CCDC88C;CACNA1A;ANK1;CACNA1G |

| | | | | | | |
|---|---|---|---|---|---|---|
| Pelizaeus-Merzbacher Disease | 4/27 | 0.000874 | 0.033391 | 10.72 | 75.46 | PRDX5;MAG;PDXP;MBP |
| Hemangioblastoma | 5/47 | 0.000937 | 0.034942 | 7.35 | 51.25 | VTN;SST;ELOB;CLU;VEGFA |
| Peripheral demyelinating neuropathy | 5/47 | 0.000937 | 0.034942 | 7.35 | 51.25 | PRDX5;MAG;ATP6;PDXP;LITAF |
| Mammary Neoplasms | 58/2387 | 0.000939 | 0.034942 | 1.63 | 11.36 | VIPR1;ARF1;TENM4;KCNK9;PEBP1;CLU;CX3CL1;GLS;SYNE1;RIMS2;RBM3;TUBA1B;PPP2R1A;CTSL;SLC39A6;TIMP3;TMSB10;COX8A;KLF10;MAP2K1;HSBP1;TSC2;CSNK1D;MIF;DYNLL1;ANK1;FOXP1;TUBB2B;PKM;SGK1;DSC3;NOTCH2;SRSF1;FBLN1;CST3;LDHB;PRDX2;SDCBP;VTN;GPC3;G3BP2;RPL13;NDN;APOE;CIZ1;CADM1;ANGPT1;ATP2B2;RPL23A;VEGFA;PPP1CA;MLLT11;COL1A2;TMEM158;TSPAN13;SST;ID4;GNAS |
| Bronchopulmonary Dysplasia | 10/188 | 0.000996 | 0.036537 | 3.50 | 24.19 | GABRB2;TENM4;ANGPT1;SST;SCN8A;GNAS;MIF;SCN4B;DGKH;VEGFA |
| Myoclonic Seizures | 9/156 | 0.001002 | 0.036537 | 3.81 | 26.30 | NRP2;SST;TBC1D24;TSC2;APOE;SCN2A;CLU;SCN1A;HCN1 |
| Generalized seizures | 8/126 | 0.001025 | 0.036537 | 4.21 | 28.98 | NRP2;SST;SCN8A;TSC2;APOE;CLU;SCN1A;HCN1 |
| Infantile Spasm | 5/48 | 0.001032 | 0.036537 | 7.18 | 49.36 | RIMS2;TBC1D24;TSC2;SCN2A;SCN1A |
| hearing impairment | 13/294 | 0.001046 | 0.036537 | 2.89 | 19.86 | NOTCH2;GABBR2;NAGA;FANCE;PQBP1;COL1A2;GRM7;GPC3;TBC1D24;GNAS;APOE;GPC4;BCOR |
| Cerebral Amyloid Angiopathy, Hereditary | 3/13 | 0.001058 | 0.036537 | 18.44 | 126.32 | CST3;SYP;APOE |
| Hallucinations, Visual | 3/13 | 0.001058 | 0.036537 | 18.44 | 126.32 | CACNA1A;APOE;SNCA |
| Macular retinal edema | 3/13 | 0.001058 | 0.036537 | 18.44 | 126.32 | SST;APOE;VEGFA |
| Frontotemporal Lobar Degeneration | 7/98 | 0.001059 | 0.036537 | 4.77 | 32.66 | RIMS2;CST3;APOE;SYP;RAN;SNCA;VEGFA |
| Malformations of Cortical Development | 7/99 | 0.001124 | 0.038473 | 4.72 | 32.03 | TUBB2B;TUBA1A;TUBB2A;TSC2;CDK13;DLGAP2;SCN1A |
| Neck webbing | 5/49 | 0.001135 | 0.038546 | 7.02 | 47.58 | PPP1CB;MAP2K1;GPC3;GPC4;BCOR |
| Long QT Syndrome | 6/73 | 0.001163 | 0.039098 | 5.54 | 37.43 | RYR2;KCNJ9;NOS1AP;CALM3;SCN4B;CALM2 |
| Ventricular Septal Defects | 11/226 | 0.001169 | 0.039098 | 3.19 | 21.55 | PPP1CB;RYR2;MAP2K1;GPC3;PRPF31;APOE;GPC4;BCOR;CDK13;PQBP1;VEGFA |
| Pervasive Development Disorder | 8/129 | 0.001193 | 0.039585 | 4.10 | 27.63 | YWHAE;GRM7;MACROD2;APOE;SCN2A;SHANK2;FOXP1;CACNA1G |
| Childhood onset | 5/50 | 0.001244 | 0.040349 | 6.86 | 45.88 | SNRPN;RPS29;PLEKHG5;SYNE1;SCN1A |
| Idiopathic generalized epilepsy | 5/50 | 0.001244 | 0.040349 | 6.86 | 45.88 | BRD2;SCN2A;SCN1A;CACNA1G;HCN1 |
| Lymphedema | 5/50 | 0.001244 | 0.040349 | 6.86 | 45.88 | YWHAE;MAP2K1;NAGA;PTPN14;VEGFA |
| Hemangioma | 7/101 | 0.001263 | 0.04065 | 4.61 | 30.80 | PTPA;THBD;NRP2;ANGPT1;TSC2;CLU;VEGFA |
| Malignant neoplasm of breast | 106/5054 | 0.001319 | 0.042123 | 1.45 | 9.64 | VIPR1;PEBP1;CLU;GLS;RIMS2;PPP2R1A;SLC39A6;MAP2K1;SNRPN;TSC2;CSNK1D;MIF;ANK1;FOXP1;SUB1;HECW1;GEMIN4;AMPH;PPIB;DSC3;NOTCH2;PFKFB3;RTN1;ANKRD11;ITPR1;FBLN1;RPAP1;CACNA1G;LDHB;PRDX2;SDCBP;VTN;PRDX5;UBC;G3BP2;NDN;APOE;BANP;PRPF38B;CADM1;TIPARP;FANCE;ZWINT;MLLT11;PPP1R1A;CCDC88C;GNB1;ID4;GNAS;CALM3;CALM2;YWHAE;ARF1;NRP2;KCNK9;COX4I1;SLC7A11;CX3CL1;SYNE1;RBM3;TUBA1B;CTSL;MCF2L;DNER;SUMO2;TIMP3;COX8A;KLF10;BRD2;RPL13A;DYNLL1;YWHAZ;TGFBR3;PKM;NOS1AP;ARHGEF2;SGK1;PTMA;AHNAK;DOT1L;SRSF1;ADAM22;PDXP;COL19A1;PTPA;CST3;GPC3;RPL13;POLR2F;SNCA;ANGPT1;USP22;ATP2B2;RPL23A;PTPN14;VEGFA;PPP1CA;NR6A1;TMEM158;TSPAN13;PSMC3;SST;FAM131A;PRPF31;RAN;HCN1 |
| Maxillary Sinus Squamous Cell Carcinoma | 3/14 | 0.001331 | 0.042197 | 16.76 | 110.99 | LDHB;PPP1CA;VEGFA |
| Mental deficiency | 30/1032 | 0.001401 | 0.044092 | 1.91 | 12.54 | KCNK9;TUSC3;SATB2;ANKRD11;ITPR1;ATP1A3;PQBP1;CACNA1G;PPP2R1A;SCN1A;GABBR2;MAP2K1;SNRPN;TSC2;NAGA;SYP;FANCE;ANK1;FOXP1;ALDH4A1;TUBB2B;SCN8A;CCDC88C;GNB1;GNAS;PRPF31;BCOR;CDK13;SHANK2;HCN1 |
| Epileptic Seizures | 7/103 | 0.001416 | 0.044236 | 4.52 | 29.64 | NRP2;SST;TSC2;APOE;CLU;SCN1A;HCN1 |
| Machado-Joseph Disease | 6/76 | 0.001434 | 0.044477 | 5.30 | 34.71 | CST3;UBB;GEMIN4;CACNA1A;APOE;SKP1 |

| | | | | | | |
|---|---|---|---|---|---|---|
| Lung Neoplasms | 33/1177 | 0.001478 | 0.044884 | 1.84 | 12.01 | NOTCH2;YWHAE;SRSF1;CACNA1A;CLU;GLS;PTPA;LDHB;SDCBP;CTSL;PGGT1B;GPC3;TIMP3;APOE;TMSB10;MAP2K1;CIZ1;CADM1;ANGPT1;KCNJ9;TSC2;MIF;ANK1;YWHAZ;VEGFA;PKM;GSTA4;RPS29;SST;SUB1;CPE;AMPH;PTMA |
| Gait, Unsteady | 5/52 | 0.001487 | 0.044884 | 6.57 | 42.76 | TRPC3;CCDC88C;ATP1A3;NAT8L;CACNA1G |
| Congenital long QT syndrome | 4/31 | 0.001489 | 0.044884 | 9.13 | 59.40 | NOS1AP;CALM3;CALM2;SCN4B |
| Early infantile epileptic encephalopathy with suppression bursts | 4/31 | 0.001489 | 0.044884 | 9.13 | 59.40 | KCNT1;SCN8A;SCN2A;SCN1A |
| Encephalopathies | 11/234 | 0.001544 | 0.046178 | 3.08 | 19.91 | PRDX2;GRIN2A;SST;SCN8A;TSC2;ATP1A3;APOE;SCN2A;SCN1A;VEGFA;SNCA |
| Generalized hypotonia | 21/633 | 0.001558 | 0.046178 | 2.17 | 14.00 | SNRPN;CACNA1A;NAGA;DDOST;FOXP1;PPP1CB;ALDH6A1;COL1A2;KCNT1;TUBA1A;TUBB2A;PPP2R1A;SCN8A;GNB1;GPC3;NDN;SPEG;GPC4;BCOR;CDK13;NAT8L |
| Dull intelligence | 28/947 | 0.001575 | 0.046178 | 1.94 | 12.50 | KCNK9;TUSC3;SATB2;ANKRD11;ITPR1;ATP1A3;PQBP1;PPP2R1A;SCN1A;GABBR2;MAP2K1;SNRPN;TSC2;NAGA;SYP;FANCE;ANK1;FOXP1;ALDH4A1;TUBB2B;SCN8A;CCDC88C;GNB1;GNAS;PRPF31;BCOR;CDK13;HCN1 |
| Low intelligence | 28/947 | 0.001575 | 0.046178 | 1.94 | 12.50 | KCNK9;TUSC3;SATB2;ANKRD11;ITPR1;ATP1A3;PQBP1;PPP2R1A;SCN1A;GABBR2;MAP2K1;SNRPN;TSC2;NAGA;SYP;FANCE;ANK1;FOXP1;ALDH4A1;TUBB2B;SCN8A;CCDC88C;GNB1;GNAS;PRPF31;BCOR;CDK13;HCN1 |
| Plaque, Amyloid | 11/235 | 0.001597 | 0.04629 | 3.06 | 19.72 | CST3;RIMS2;OLFM1;UBB;APLP1;MBP;APOE;SLC7A11;SYP;CLU;SNCA |
| Other specified types of schizophrenia, unspecified | 5/53 | 0.00162 | 0.04629 | 6.43 | 41.31 | GNAS;MBP;APOE;YWHAZ;YWHAH |
| Multiple Sclerosis, Relapsing-Remitting | 6/78 | 0.00164 | 0.04629 | 5.15 | 33.05 | PRDX5;PDXP;MBP;APOE;CX3CL1;VEGFA |
| Dyskinetic syndrome | 6/78 | 0.00164 | 0.04629 | 5.15 | 33.05 | MCF2L;SCN8A;APOE;SCN2A;SNCA;VEGFA |
| Ductal Carcinoma In Situ with Microinvasion | 3/15 | 0.001644 | 0.04629 | 15.36 | 98.49 | COX8A;PEBP1;PRPF31 |
| Middle East Respiratory Syndrome | 3/15 | 0.001644 | 0.04629 | 15.36 | 98.49 | VTN;CTSL;PPP1CA |
| Cognition Disorders | 11/237 | 0.001708 | 0.047775 | 3.03 | 19.34 | SST;SATB2;SCN8A;AMPH;APOE;SYP;YWHAZ;SHANK2;SCN1A;FOXP1;SNCA |
| Dizziness | 5/54 | 0.001763 | 0.048984 | 6.30 | 39.94 | RYR2;CACNA1A;NAGA;CALM3;CALM2 |

*Supplementary Table 5. Reagents details.*

| Antibodies used for immunocytochemistry/flow-cytometry | | | |
|---|---|---|---|
| | **Antibody** | **Dilution** | **Company Cat # and RRID** |
| Pluripotency Markers | Mouse anti OCT 3\4 | 1:100 | Santa Cruz Cat# sc-5279, RRID: AB_628051 |
| | Mouse anti TRA-1-60 | 1:50 | R & D Cat# MAB4770, RRID: AB_2119062 |
| | Mouse anti SSEA-4 | 1:200 ICC1:100 FACS | Santa Cruz Cat# SC-21704, RRID: AB_628289 |
| | Rabbit anti SOX2 | 1:200 | Abcam Cat# ab97959, RRID: AB_2341193 |
| | Rabbit anti NANOG | 1:150 | Abcam Cat# 21624, RRID: AB_446437 |
| | Mouse anti IgG3 Isotype | 1:100 | Bio-legend Cat# 330405, RRID: AB_1089207 |
| Differentiation Markers | Rabbit anti 68 kDa Neurofilament | 1:100 | Abcam Cat# ab52989, RRID: AB_869924 |
| | Rabbit anti α-Fetoprotein | 1:1 | ScyTek Cat# A00058, RRID: AB_651223 |
| | Rabbit anti α-smooth muscle actin | 1:100 | Abcam Cat# ab32575, RRID: AB_722538 |
| | Mouse anti Tubulin, beta III (TUJ1) | 1:200 | Sigma-Aldrich Cat# MAB1637, RRID:AB_2210524 |
| | Mouse anti Cardiac troponin T | 1:100 | Abcam Cat# ab8295, RRID:AB_306445 |
| | Rabbit anti SOX17 | 1:100 | Abcam Cat# ab 224637, RRID:AB_2801385 |
| Secondary antibodies | Alexa Fluor 488-conjugated Anti Mouse | 1:500 | Jackson Immuonoresearch inc. Cat# 715-545-150, RRID: AB_2340846 |
| | Alexa Fluor 594-conjugated Anti Rabbit | 1:500 | Jackson Immuonoresearch inc. Cat# 711-585-152, RRID: AB_2340621 |
| | Goat anti-Rabbit IgG (H + L) Secondary Antibody, Alexa Fluor 488 conjugate | 1:500 | invitrogen A-11034 |
| Episomal Plasmids | pCXLE-hOCT3/4-shp53 | Addgene #27077 | |
| | pCXLE-hSK | Addgene #27078 | |
| | pCXLE-hUL | Addgene #27080 | |
| | pEP4 E02S ET2K | Addgene #20927 | |
| | pCXWB-EBNA1 | Addgene #37624 | |

***Data sets***

Data sets 1-5 are openly available at Google drive (https://drive.google.com/drive/folders/1D-W3LL0gSn7yWZ8h1ynLq-y29KaCLDea?usp=sharing)

Data set 1: Undifferentiated hiPSC_differentially expressed genes.tsv
Data set 2: hiPSC-derived cortical neurons_differentially expressed genes.tsv
Data set 3: Mouse cortex_differentially expressed genes.tsv
Data set 4: Gene Ontology (GO) Molecular Function analysis.xlsx
Data set 5: DisGeNet analysis.xlsx
